# An LRH-1 agonist reverses hepatic metabolic dysfunction in obesity

**DOI:** 10.64898/2026.10.28.761933

**Authors:** Suzanne G. Mays, André R. Cuevas, Racheal M. Spurlin, Rohini Polavarapu, Chih-Yu Chen, Zhenxin Hou, Michael L. Cato, Rita Swinger, Elizabeth J. Millings, Autumn R. Flynn, Debipreeta Bhowmik, Henry Zecca, Wen-Yu Hsiao, Sarah M. Abraham, Alyssa M. Johnson, Nathan T. Jui, Eric A. Ortlund, John W. Calvert

## Abstract

Liver receptor homolog-1 (LRH-1; NR5A2) is a nuclear receptor that controls metabolic gene programs important for liver health. Here, we address two challenges with dissecting ligand-driven LRH-1 metabolism in vivo. First, we generate ME610, a potent LRH-1 agonist with favorable pharmacokinetic properties. Second, we engineer *Nr5a2^humLBD^* mice, edited to recapitulate the ligand response of the human receptor. ME610 dose-dependently induces LRH-1 target genes and reduces triglycerides in livers of *Nr5a2^humLBD^* mice. In a mouse model of diet-induced obesity (DIO), ME610 dramatically improves insulin sensitivity and reduces liver fat. Multi-omics integration of transcriptomics, lipidomics, and metabolomics data demonstrates a coordinated reversal of DIO-induced hepatic metabolic stress via changes in lipid, amino acid, and energy metabolism, with peroxisomal lipid metabolism emerging as a central process correlated to efficacy. These studies uncover new pathways through which LRH-1 regulates metabolism and demonstrate the therapeutic potential of synthetic LRH-1 agonists in obesity-associated metabolic diseases.

## Introduction

Metabolic dysfunction-associated steatotic liver disease (MASLD) is a chronic liver disease estimated to affect more than 32% of the world’s population^1^. Formerly known as non-alcoholic fatty liver disease, MASLD is associated with obesity, type II diabetes mellitus, or other symptoms of metabolic dysfunction such as insulin resistance or high circulating triglycerides (TGs)^1^. MASLD progression leads to metabolic dysfunction-associated steatohepatitis (MASH), liver injury, increased risk of hepatocellular carcinoma, and liver failure^1–3^. Obesity and insulin resistance are major drivers of MASLD, contributing to toxic accumulation of fatty acids (FAs) and other proinflammatory lipids in the liver. Aberrant lipid accumulation can cause mitochondrial dysfunction, reactive oxygen species production, and endoplasmic reticulum (ER)-stress, contributing to MASH development^2, 4^. In the early stages of MASLD, mitochondrial FA β-oxidation (FAO) is induced to compensate for lipid overload^5, 6^, increasing flux through the tricarboxylic acid (TCA) cycle and oxidative phosphorylation^7^. Peroxisomal FAO (PxFAO) is induced to ease the burden on the mitochondrial pathway but, paradoxically, can also contribute to mitochondrial dysfunction and oxidative stress^8, 9^. Therapeutics that ease metabolic stress in MASLD may prevent disease progression.

Liver receptor homolog-1 (LRH-1; gene symbol NR5A2) is a nuclear receptor that acts as a master regulator of metabolic programs relevant to MASLD. It is highly expressed in the liver, where it regulates de novo lipogenesis^10^, bile acid and cholesterol metabolism^11–14^, glucose capture^15^, one-carbon metabolism^16^, and the acute phase response^17^. It also modulates hepatic TG metabolism^18^ and ER-stress resolution^19^. Genetic and pharmacological studies demonstrate its potential as a therapeutic target. Although full knockout is not possible due to embryonic lethality^20^, LRH-1 haploinsufficiency promotes obesity in mice on high fat diet (HFD)^21^. Conditional LRH-1 knockout in hepatocytes of male mice causes steatosis that is exacerbated with HFD^22^. Importantly, a phospholipid LRH-1 agonist, dilauroylphosphatidylcholine (DLPC), improves insulin resistance and reduces hepatic steatosis in a mouse model of HFD-induced obesity (DIO)^10^. DLPC efficacy is associated with suppression of lipogenic genes in the liver, although the mechanism of action is not fully elucidated. More recently, the bacterial lipid vaccenic acid has been identified as an LRH-1 ligand with efficacy in a mouse DIO model^23^.

The discovery that LRH-1 activity is modulated by lipid ligands motivated the search for synthetic agonists with improved potency and specificity compared to natural lipids. These have been challenging to develop, in part due to the receptor’s hydrophobic binding pocket and a poor understanding of how ligands drive its activity. We have undertaken a structure-guided approach to improve weak modulators with the hexahydropentalene (6HP) scaffold (*e.g.,* RJW100)^24, 25^. We discovered that small modifications to the 6HP scaffold can cause unpredictable positioning in the pocket, helping to explain an unclear structure-activity relationship (SAR)^26^. LRH-1 crystal structures with RJW100 and DLPC revealed opportunities to anchor the compound in the pocket at two polar sites, greatly improving potency and SAR^27–32^. Extending the RJW100 hexyl “tail” and adding a terminal carboxylic acid increases activity by mimicking interactions made by DLPC (*e.g.,* agonist 10CA)^29, 31^. Strategic incorporation of a sulfamide promotes anchoring hydrogen bonding interactions by displacing a structural water molecule, increasing EC_50_ by over 100-fold (*e.g.,* 6N and 6N-10CA)^27, 31, 33^. While activity is greatly improved over earlier derivatives, hydrophobicity of these compounds may limit use in vivo. The long, hydrocarbon “tails” are key sites of hydrophobicity.

In these studies, we synthesize and evaluate a set of compounds with solubilizing ethers in the tails of the LRH-1 agonists 10CA and 6N-10CA. The monoether version of 6N-10CA (ME610) is more soluble than the parent compound and is equally active in reporter assays. ME610 shows striking efficacy in a DIO model with novel humanized LRH-1 mice, improving insulin resistance and reducing intrahepatic fat. Efficacy is associated with reversal of diet-induced metabolic programs, including an important role of PxFAO.

## Results

### A monoether variant of 6N-10CA is a potent LRH-1 agonist with improved solubility

We incorporated ethers within the agonists’ hydrocarbon “tails,” aiming to add polarity while avoiding charged or protonated species likely to be incompatible with the hydrophobic binding pocket. We synthesized monoether and diether variants based on two of our most active LRH-1 agonists: 10CA (**1**)^29^ and 6N-10CA (**2**)^31^ (Fig. 1a). To make these molecules, we first devised routes to generate ether-containing tails (Extended Data Fig. 1a-b) and incorporated them into the 6HP scaffold via the Whitby cyclization^28, 29, 31^, yielding compounds **3-6** (Fig. 1a).

**Figure 1.**
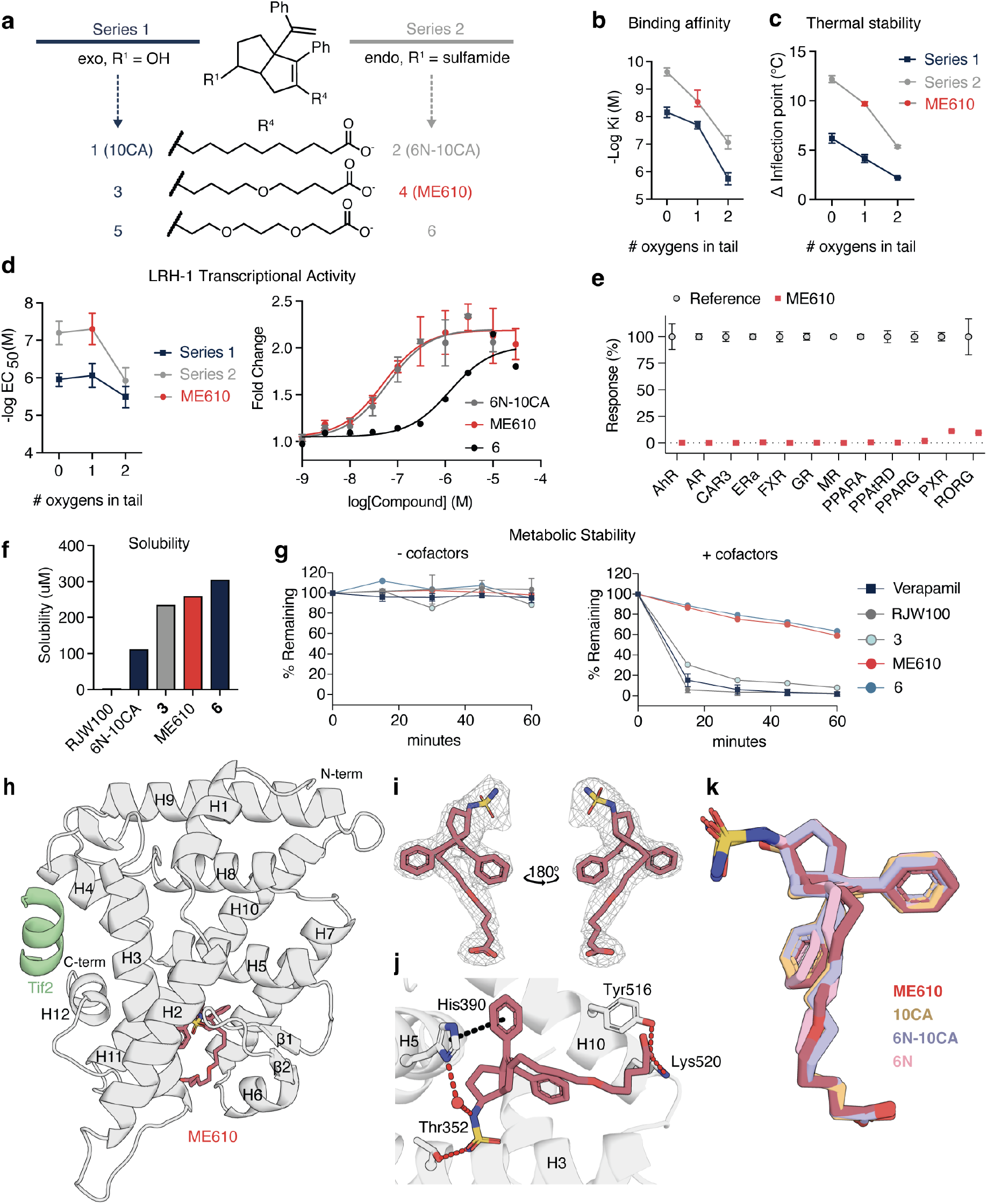
ME610 is similarly active to 6N-10CA and has improved solubility. **a.** Design strategy for monoether and diether variants of 10CA (Series 1: navy) or 6N-10CA (Series 2: grey). ME610 (*red*) is the monoether from Series 2. **b**. LRH-1 binding affinities for Series 1 and 2 compounds, plotted as a function of number of tail oxygen atoms. Compiled data from three experiments were used to calculate pKi, and error bars represent the 95% CI. **c.** Thermal shift assays. Purified LRH-1 LBD was incubated overnight with 10 µM compound, and the complex was heated to determine the inflection point, a measure of thermal stability. Values reported as the difference in inflection points for each compound *vs* controls (LRH-1 LBD incubated with DMSO). **d.** LRH-1 transcriptional activity measured by luciferase reporter assay. *Left,* pEC_50_ values for all compounds. *Right,* dose response curves for Series 2 compounds showing ME610 is equally activating to 6N-10CA. **e.** ME610 does not activate AhR or several other NRs in luciferase reporter assays. **f.** Ǫuantification of aqueous solubility for the indicated compounds, measured in PBS at room temperature. **g**. Metabolic stability of compounds in male mouse liver microsomes with cofactors *left* or without *right*. ME610 and compound 6 are more stabilizing than compound **3** and the LRH-1 agonist RJW100. Verapamil is used as a control. **h.** Overall structure of the LRH-1 LBD bound to ME610 and a fragment of the Tif2 coregulator (2.7 Å). **i.** Omit map (F_O_ -F_C_, contoured at 2 σ) showing that a single enantiomer of ME610 is bound in the structure. **j.** Close view of ME610 in the pocket showing key hydrogen bonding interactions (red dotted lines) and pi-stacking with residue H390 (black dotted line). A water molecule participating in a bridging interaction is shown as a red sphere. **k.** Superposition of ME610 with ligands in LRH-1 crystal structures with related compounds, including 6N (PDB 6OǪY)^27^, 10CA (PDB 7JYD)^28^, and 6N-10CA (PBD 7TT8)^31^ shows close similarity in binding poses.

In vitro evaluation of the ether-containing compounds reveals clear SAR (Fig. 1b-d; S1a-b). Derivatives of 6N-10CA are more potent, stabilizing, and active than 10CA derivatives, regardless of tail composition (Fig. 1b-d). Within each series, tail oxygen atoms progressively impair ligand binding affinity and reduce thermal stabilizing effects on purified LRH-1 LBD, suggesting some incompatibility with the pocket (Fig. 1b-c). The monoether version of 6N-10CA (**4**, hereafter “ME610”) is the most potent ether-containing derivative, with a 2.9 nM binding affinity and equal potency to 6N-10CA in cellular LRH-1 reporter assays (EC_50_ ∼50 nM for both). In contrast, the corresponding diether **6** has micromolar affinity and EC_50_ (Fig. 1b-d). ME610 also exhibits LRH-1 selectivity: it does not modulate the aryl hydrocarbon receptor (AhR) or a panel of nuclear receptors at a concentration yielding maximum LRH-1 activation (Fig. 1e).

As designed, tail ethers improve aqueous solubility and confer favorable ADME/PK profiles. Monoether and diether variants are 2-3-fold more soluble than 6N-10CA and 90-100-fold more soluble than RJW100 (Fig. 1f). ME610 and **6** exhibit greater metabolic stability in mouse liver microsomes relative to RJW100 (Fig. 1g). In rat PK studies, ME610 reaches a mean C_max_ of 2690 ng/ml when administered via intraperitoneal (i.p.) injection, equivalent to ∼11x the EC_90_ for activating LRH-1 in vitro (Table S1; Extended Data Fig. 1c). The C_max_ of **6** is higher than ME610 (5500 ng/ml) but does not reach its EC_90_. Similar trends are seen with AUC_inf_ (Extended Data Fig. 1c) and with oral administration (Table S1). Therefore, poor potency of **6** likely limits its ability to activate LRH-1 in vivo despite improved exposure, whereas ME610 reaches levels adequate to drive activity.

### ME610 maintains important interactions in the LRH-1 crystal structure

To understand how the ether affects ligand binding mode, we determined the x-ray crystal structure of LRH-1 LBD with ME610 (2.7Å resolution; Table S2; Fig. 1h). LRH-1 protein conformation is similar to the 6N-10CA-LRH-1 co-crystal structure (root mean square deviation (RMSD) of 0.2Å)^31^. The Tif2 coactivator peptide (Fig. 1h, *green*) is bound at the expected site at the activation function surface (AFS). Clear electron density around ME610 indicates a single enantiomer is bound (*1S,3aR,CR*; Fig. 1i). Superposition of ME610 and 6N-10CA ligands reveals similar tail trajectories, and both compounds make contacts important for activity (Tyr516 and Lys520)^28, 31^ via the terminal carboxylic acids (Fig. 1j-k). Like 6N-10CA and related agonists, ME610 forms hydrogen bonds with residue Thr352 and π-π stacking interactions with His390, also important for binding and activation (Fig. 1k)^34, 35^.

Molecular dynamics simulations provide insights into why ME610 maintains similar activity to 6N-10CA despite having lower affinity (Extended Data Fig. 1d-f; Fig. S1d-e). Both ME610 and **6** interact with Tyr516 and Lys520 for less simulation time than 6N-10CA, mirroring trends in binding affinities (Extended Data Fig. 1d). However, when Tif2 is added to the complex to model the active state, 6N-10CA and ME610 induce strong connectivity throughout the AFS while **6** does not, suggesting **6** has an impaired ability to activate LRH-1 via allosteric communication with the AFS (Extended Data Fig. 1e-f). These structural analyses, together with the activity and PK profiles, demonstrate the potential of ME610 as an LRH-1 agonist in vivo.

### Humanizing the LRH-1 ligand binding domain alters gene expression in the liver

We next generated a humanized LRH-1 mouse designed for testing ME610 and other modulators tailored to the human LRH-1 binding pocket. Mouse LRH-1 (mLRH-1) shares 86.5% sequence identity with the human ortholog (hLRH-1) but differs in amino acids H438-F443 near the mouth of the ligand binding pocket (Fig. 2a). Substituting this mouse sequence into hLRH-1 reduces flexibility in the vicinity^36^ that is important for ligand-driven activation^37, 38^. The phenylalanine (leucine in hLRH-1) may obstruct the binding pocket (Fig. 2a). In luciferase reporter assays, mLRH-1 displays higher basal activity than hLRH-1 (Fig. 2b) but is only weakly activated by ME610. Mutating H438-F443 to the human sequence restores ME610-induced activity in the reporter assay (Fig. 2c). We therefore used CRISPR-Cas9 to replace 18 bp encoding residues 438-443 with the human sequence in C57BL/6J mice. Mice homozygous for the edited region (hereafter *Nr5a2^humLBD^*) breed and develop normally. At 10 weeks of age, *Nr5a2^humLBD^* mice express similar LRH-1 protein levels to WT C57BL/6J mice in the liver, pancreas, ileum, and colon (Extended Data Fig. 2a-b). Body weights and organ weights are indistinguishable from WT mice (Extended Data Fig. 2c-e).

**Figure 2.**
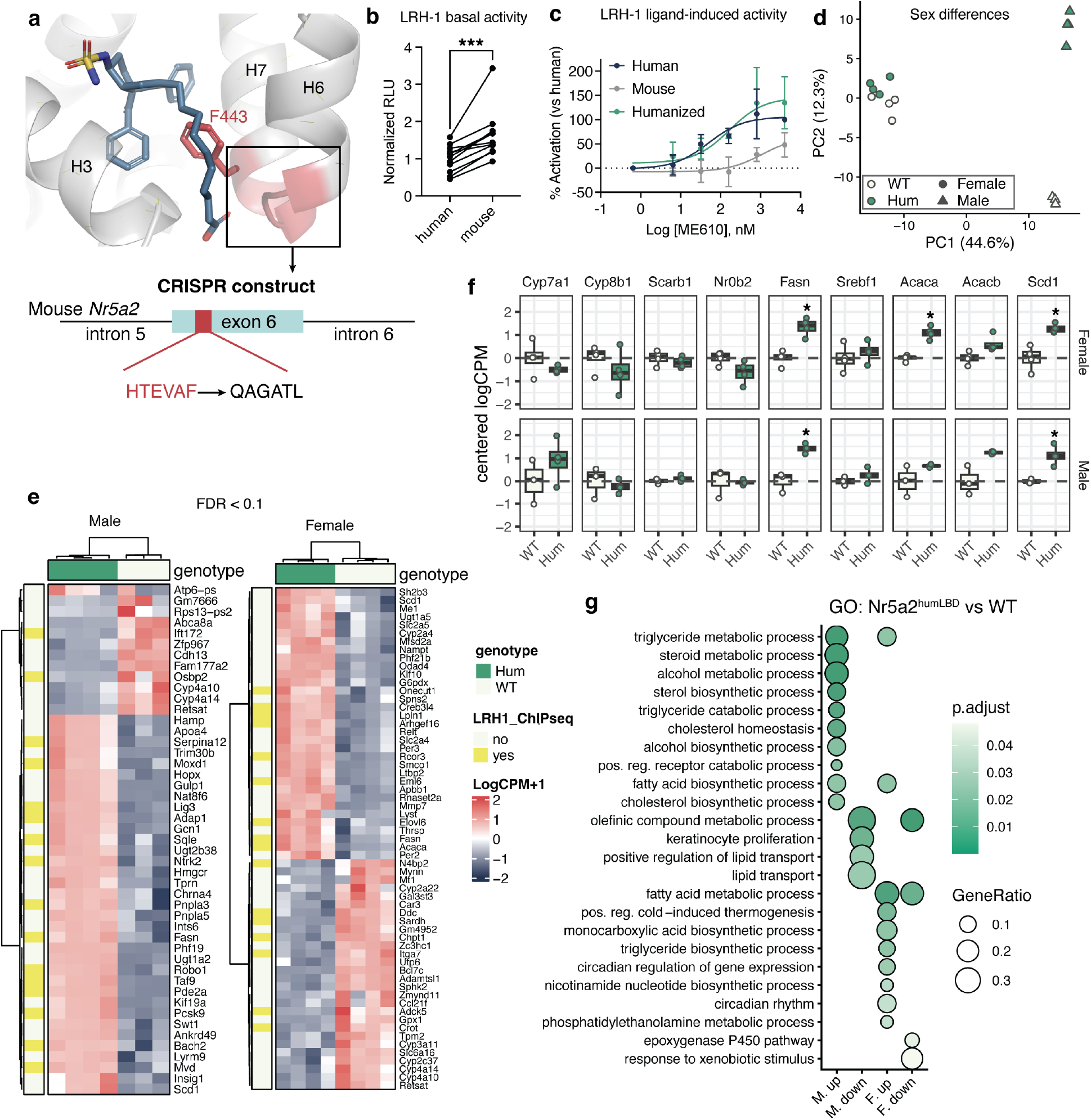
Generation of a humanized Lrh-1 mouse (*Nr5a2^humLBD^)* with increased ligand sensitivity. **a.** Structural model illustrating the rationale for making the humanized mouse. The mouse Lrh-1 protein structure (from PDB 1PK5)^55^ is superposed with a ligand-bound human LRH-1 structure (PDB 7TT8)^31^. Edited region is highlighted in red. Mouse Lrh-1 residue F443 on helix 7 (H7) (shown as sticks in red) may clash with bound ligands. H, helix. *Bottom,* schematic of the genetic locus of the edited region and the amino acid changes introduced. **b.** Reporter assay in Hela cells showing increased LRH-1 transcriptional activity at the SHP promoter in the presences of mLrh-1 *versus* hLRH-1. Each point represents the mean of three replicates from independent experiments. Error bars are SD, boxes show medians; whiskers show the range. *, p < 0.05 by two-tailed, unpaired Mann-Whitney test). **c.** Comparison of dose-response to ME610 for human, mouse, and humanized versions of LRH-1. Each point represents the mean of three replicates from three independent experiments. Error bars are SD. Values on the y-axis are expressed as % of Emax for the human curve. **d.** Principal component analysis from gene expression data in livers of *Nr5a2^humLBD^ versus* WT mice shows a major effect of sex in driving gene expression differences (PC1). **e.** Heatmaps from whole genome RNA-seq of mouse livers, showing distribution of genes with FDR < 0.1 for *Nr5a2^humLBD^ versus* WT male (*left*) and female (*right*) mice. **f.** Humanizing the pocket upregulates a subset of DLPC-controlled genes related to lipogenesis. Each point is the logCPM value for an individual mouse in the study (centered around the mean logCPM of WT mice for each gene). Error bars are SD. *, FDR < 0.1 from the RNA-seq statistical analysis. **g.** Dot plot showing the most significantly enriched gene ontology (GO) categories for each comparison. Dot size represents the GeneRatio and the shade represents adjusted p-values. Both male and female mice have upregulated genes involved in metabolic processes related to known LRH-1 functions.

Despite the phenotypic similarity, humanizing mLRH-1 alters hepatic gene expression profiles, determined by bulk mRNA sequencing (RNA-seq). We analyzed data from males and females separately after finding sex is a dominant source of variability (Fig. 2d). A small, interesting set of genes are differentially expressed in *Nr5a2^humLBD^* mice *versus* WT for each sex (FDR threshold < 0.1, Fig. 2e). These include direct LRH-1 targets, defined as genes with promoters bound by LRH-1 in published ChIP-seq datasets^39–41^ (Fig. 2e, yellow). Lipogenic genes modified by DLPC in mouse liver are differentially expressed: fatty acid synthase (*Fasn),* stearoyl-CoA desaturase 1 (*Scd1*), and acetyl-coA carboxylase 1 (*Acaca*). Upregulation of these genes in *Nr5a2^humLBD^* mice *versus* WT is the opposite change seen with DLPC and synthetic LRH-1 agonists^10, 28, 31^, consistent with higher WT mLRH-1 basal activity observed in vitro. Other DLPC targets do not change expression (Fig. 2f). Gene ontology (GO) analysis shows enrichment in categories related to TG and FA metabolism for males and females, cholesterol/steroid metabolism and lipid transport in males, and circadian rhythm, thermogenesis, and glucose transport in females (Fig. 2g). These data demonstrate that humanizing a small portion of the mLRH-1 LBD changes expression of genes relevant to liver health and suggest that the humanized receptor responds to endogenous ligands differently than the WT mouse receptor.

### ME610 modulates LRH-1 transcriptional targets and reduces triglycerides in the livers of *Nr5a2^humLBD^* mice

To investigate activity of ME610 in *Nr5a2^humLBD^* mice, we conducted a dose titration study, treating male mice twice daily by i.p. injection over 3 days (five total treatments). Four hours after the last treatment, livers were harvested for transcriptomics and lipidomics analyses. Doses of 3 mg/kg and 30 mg/kg were used for RNA-seq, with additional, lower doses included in the lipidomics analysis (Fig. 3a).

**Figure 3.**
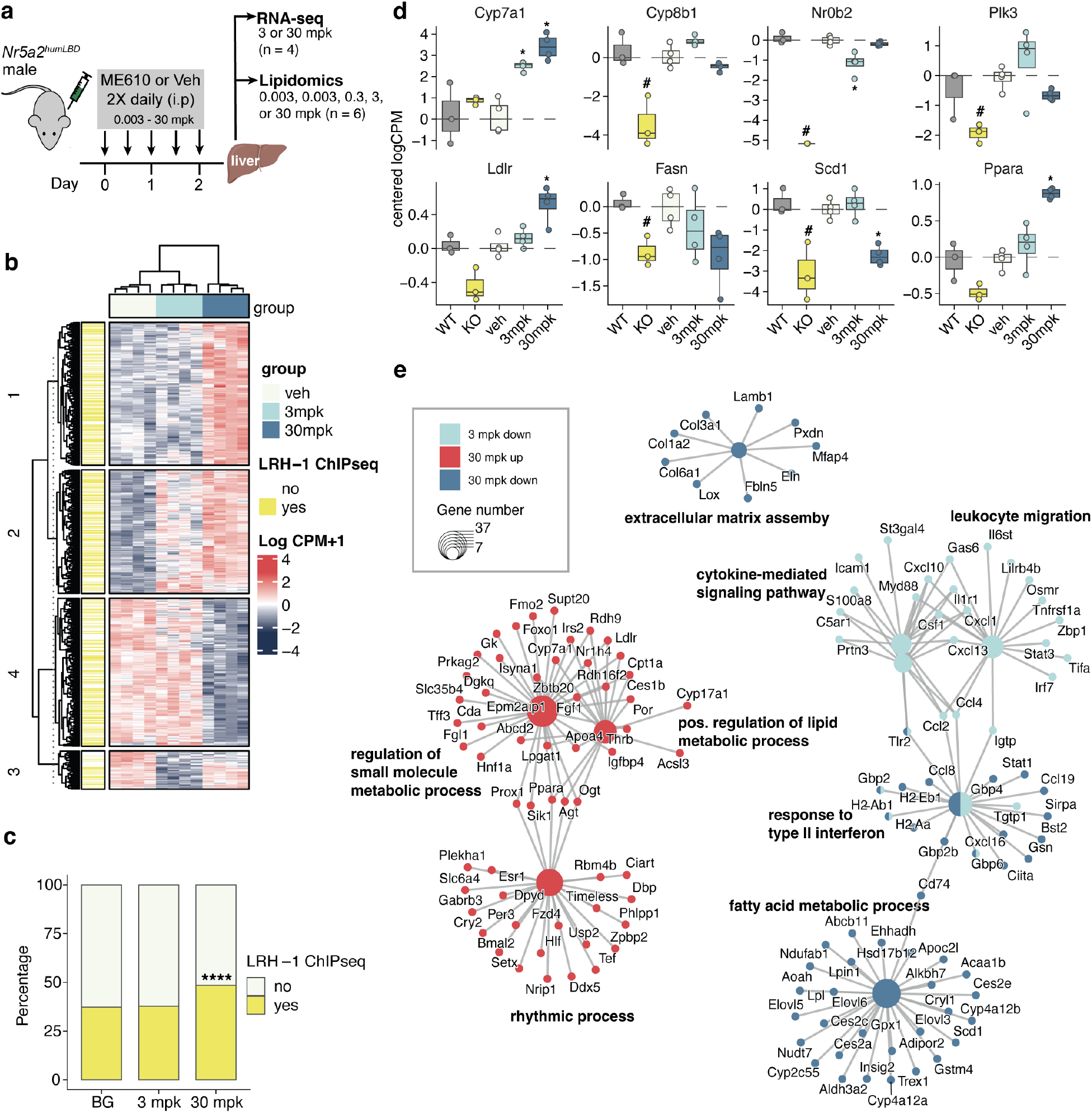
ME610 induces dose-dependent gene expression change in the livers of male *Nr5a2^humLBD^*mice. **a.** Schematic of experiment. Male *Nr5a2^humLBD^*mice were treated with vehicle or ME610 twice daily for two days. On the third day, they were given one treatment, and livers were harvested four hours later for use in RNAseq and lipidomics. **b.** Heatmap depicting scaled Log_2_ counts per million (CPM) + 1 values for the set of genes with FDR < 0.1 for either dose of ME610 *versus* vehicle controls. Columns are clustered by Euclidean distance and Ward’s agglomeration; rows are clustered by Kmeans. Left annotation indicates genes from published LRH-1 ChIP-seq datasets. **c.** Changing genes are enriched for LRH-1 ChIP-seq targets in the 30mpk group relative to background (BG, defined as all expressed genes in the dataset). ****, p < 0.001, Fisher’s exact test, Benjamin Hochberg correction for multiple comparisons. **d.** Comparison of gene expression changes in the livers of conditional Lrh-1 knockout (KO) mice *versus* in *Nr5a2^humLBD^* mice treated with ME610 at the indicated doses. KO mouse data was analyzed from a publicly available dataset, selecting data from healthy, adult male mice in which Lrh-1 was inducibly knocked out in hepatocytes^22^. Log_2_CPM values were centered on the “control” values for each dataset for visual comparison across experiments (“WT” in the KO study and “veh” in the ME610 study). Each point is a mouse in the study; boxes indicate the first quartile, median and third quartiles; whiskers are the range. *, FDR < 0.1 relative to controls. **e.** Top enriched GO categories for each comparison in the ME610 titration study plotted as a functional network, p.adj < 0.05. The top three categories for each comparison are shown. See associated source data for RNA-seq and GO statistics tables.

The RNA-seq data demonstrate modulation of LRH-1 target genes in the liver, with main effects on anti-inflammatory and lipid metabolic pathways. Groups are well-separated by principal component analysis (PCA, Extended Data Fig. 3a) and by unbiased clustering (Fig. 3b). Numbers of changing genes (FDR < 0.1) increase dose-dependently (Extended Data Fig. 3b-e). At the 30 mg/kg dose, changes to direct LRH-1 targets (from ChIP-seq datasets^39–41^) are enriched by 19% over a background of all expressed genes (Fig. 3c). Some LRH-1 target genes change in opposite directions than in published data from hepatocyte-specific *Nr5a2* knockout (KO) mice versus WT mice^22^, as might be expected for an agonist (*e.g., Ldlr, Plk3, and Ppara*, Fig. 3d). Others do not follow this pattern, changing in the same direction as KO (*e.g., Fasn, Scd1*) or changing in one condition but not the other (*e.g., Cyp8b1*). Interestingly, strong repression of *Nr0b2* occurs in the KO livers but not with ME610. This has potential for a widespread impact on LRH-1-controlled transcription, since Nr0b2 protein represses LRH-1 activity^14, 42^. Other reasons not to observe a strict reversal of the KO signature is that *Nr5a2^humLBD^* mice express the humanized receptor in all LRH-1-expressing cells from birth and were treated acutely with the agonist, different from the KO mice.

GO analysis reveals an anti-inflammatory gene signature at both dosage levels of ME610 (Fig. 3e). At 3 mg/kg, downregulated genes are highly enriched for cytokine signaling, involving genes such as *Stat3, Cxcl10, Tnfrsf1a,* and *Ccl2*. A Stat3 phosphatase (*Ptprd*)^43^ is upregulated, which could also attenuate Stat3 signaling. Downregulated genes at both doses are involved in type II interferon signaling, including *Stat1* and *Ccl1S*. We also observe repression of genes encoding guanylate binding proteins (*e.g. Gbp2, Gbp2b*), components of the inflammasome and of histocompatibility factors (*e.g. H2-Ab1* and *H2-Eb1*) at both doses. The 30 mg/kg dose has a strong GO signature related to lipid metabolism, with suppression of genes such as *Elovl5*, *Scd1, Abcb11, and Acaa1b* and upregulation of genes such as *Ldlr, Acsl3, Fgf1, Cpt1a, and Cyp7a1.* Suppression of extracellular matrix (ECM) assembly-related genes at 30 mg/kg may suggest an anti-fibrotic function^44^. Additionally, several circadian factors are upregulated, including *Cry1, Cry2*, *Per3,* and *Bmal2*.

In lipidomics data, a total of 202 lipids across multiple classes change expression with ME610 at any dose versus vehicle, with the most changes at 3 mg/kg (Fig. 4a-b; Extended Data Fig. 3f-g). ME610 induces a potent and broad reduction of TGs, evident at even the lowest dose and affecting most TG species (Fig. 4c-d). Diglyceride (DG) species, potentially derived from TG hydrolysis, begin to increase at doses ≥ 0.3 mg/kg. Unsaturated ether-linked PC (O-PC) species are consistently upregulated. Increases are also seen in certain sphingolipids. C18 dihydro-ceramide, representing 85% of the changing ceramide (Cer) species, decreases at lower doses and increases at higher doses. Sphingomyelin (SM) species, particularly with chain lengths of 32-36, increase at doses ≥ 0.3 mg/kg. Lipid class-based analysis (by lipid set enrichment analysis (LSEA)), confirms suppression of TG as a prominent effect, occurring at the lowest dose (0.003 mg/kg) and persisting for all treated groups (Fig. 4d). Other lipid classes with high sensitivity to ME610 in LSEA are O-PCs, LPCs, and SM (increased), as well as hexosylceramides (HexCer), Cer, and DG (decreased).

**Figure 4.**
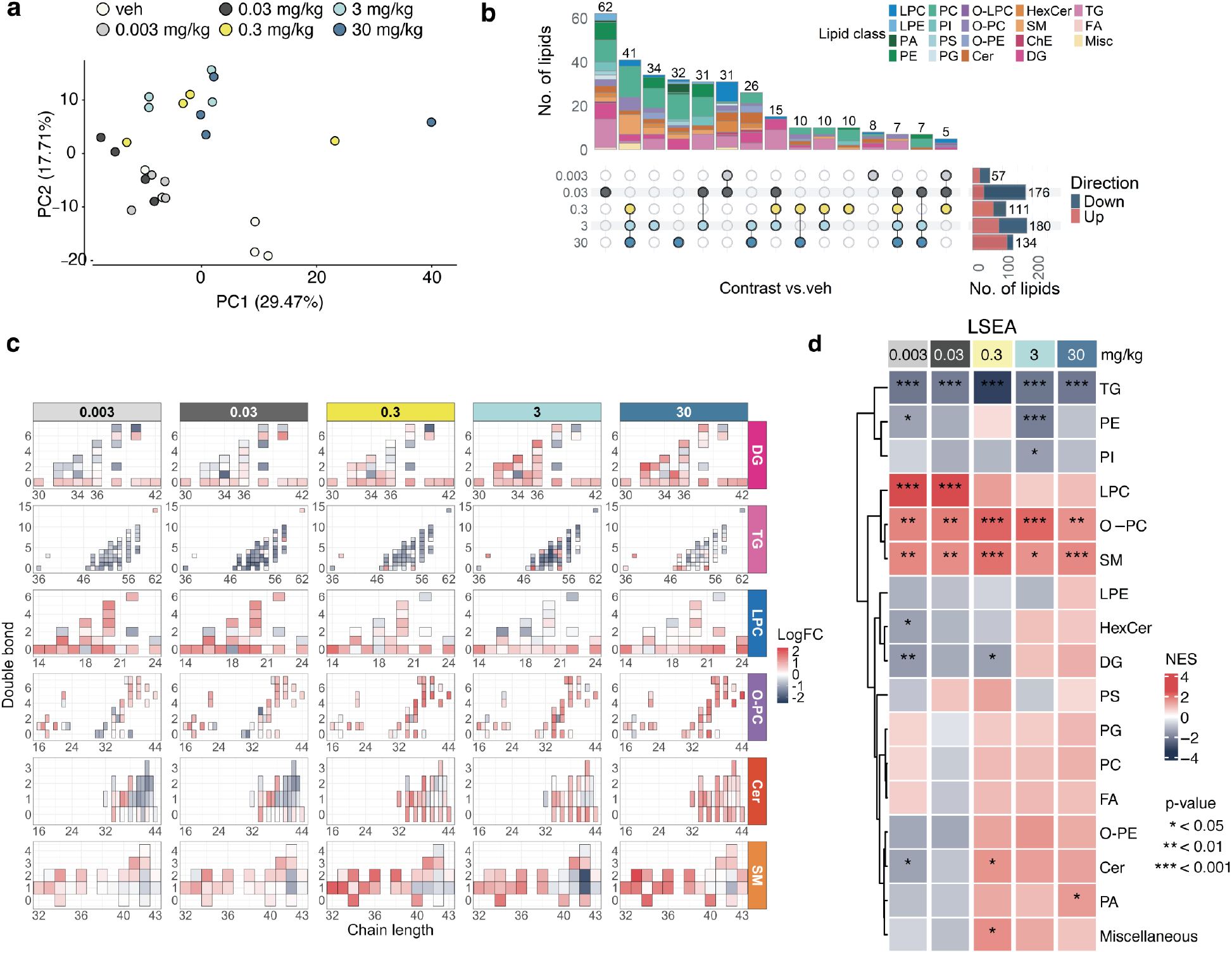
ME610 reduces hepatic triglycerides and remodels lipids in the livers of male *Nr5a2^humLBD^*mice. **a.** PCA from lipidomics dose titration study, showing group separation along PC2 (mice treated with doses 0.3 mg/kg and above cluster away from the other groups). **b.** Upset plot of changing lipids for each treatment group *versus* the vehicle-treated group (veh). Horizontal bars show numbers of upregulated (*red*) and downregulated (*dark blue*) lipid species at each dose (0.03 mg/kg and 3 mg/kg have the most changing lipids). Colors of vertical bars indicate proportions of lipid classes among the changing lipids. Overlap between changing lipids across various groups are indicated by the dots and lines below the plot. **c.** Heatmaps showing mean log fold changes (logFC) of individual diglycerides (DG), triglycerides (TG), lysophosphatidylcholines (LPC), ether-linked phosphatidylcholines (O-PC), ceramides (Cer), and sphingomyelins (SM). X-axes show total lengths of acyl chains, and y-axes show total numbers of double bonds in acyl chains. **d.** Heatmap depicting normalized enrichment scores (NES) from LSEA analysis. Abbreviations: phosphatidylethanolamine (PE); lysoPE (LPE); phosphatidylinositol (PI); hexosylceramides (HexCer); phosphatidylserine (PS); phosphatidylglycerol (PG); fatty acid (FA); ether-linked PE (O-PE); phosphatidic acid (PA). See associated source data for lipidomics and LSEA statistics tables.

The sensitivity of lipid changes is striking considering that many lipid metabolic genes change only at the highest dose. This suggests lipid changes may drive some transcriptional changes. For example, suppression of FA metabolic genes at 30 mg/ kg may reflect a compensatory response to TG depletion. Anti-inflammatory gene changes at the lower dose may be more important in driving lipid metabolism. The Ptprd-Stat3 pathway has been linked to TG reduction^43^.

### ME610 improves insulin resistance and reduces liver fat in obese mice

We next tested ME610 in a model of diet-induced obesity (DIO) in *Nr5a2^humLBD^* mice. Prior to conducting the study, we determined that gene expression changes in *Nr5a2^humLBD^*mouse livers (Fig. 2) do not alter hallmark phenotypic features of the model. *Nr5a2^humLBD^* mice gain weight and develop hepatic steatosis similarly to WT mice (Fig. S2a-e). Liver weights, insulin resistance, and adiposity in *Nr5a2^humLBD^* mice are indistinguishable from WT mice (Fig. S2f-w).

The DIO study contained three treatment groups, two receiving HFD and one receiving standard chow. We administered these diets for 12 weeks and then treated with ME610 or vehicle for four weeks while maintaining the diet (Fig. 5a). For rigorous assessment of agonist efficacy, we ensured the HFD cohort did not differ in body weight or performance in glucose tolerance tests (GTT) and insulin tolerance tests (ITT) at the onset of treatment (Extended Data Fig. 4). Males and females were analyzed separately due to differences in body weights and known differences in liver biology^45^.

**Figure 5.**
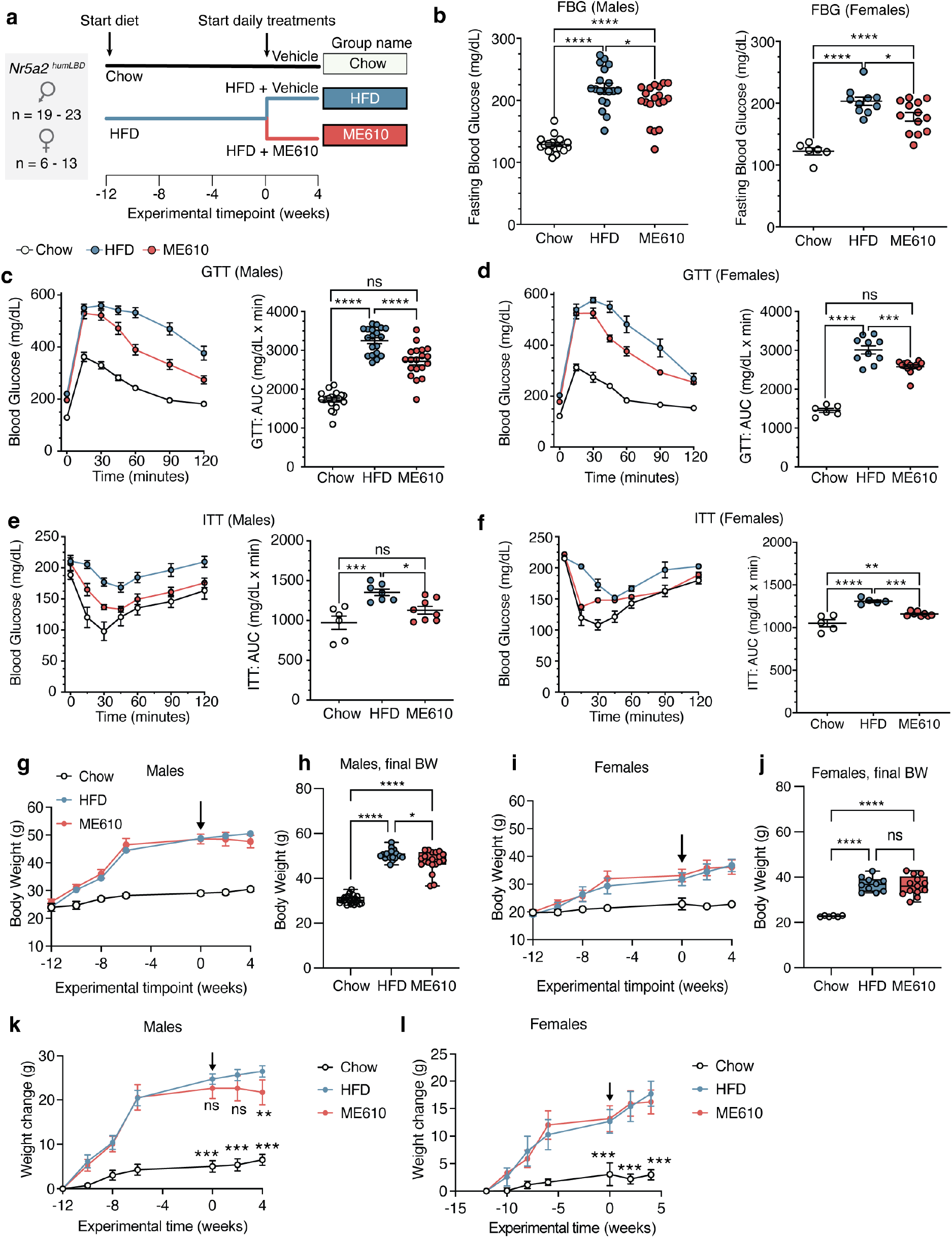
ME610 improves insulin sensitivity in a DIO mouse model. **a.** Experimental schematic. 8-week-old *Nr5a2^humLBD^* mice were fed HFD or chow for 12 weeks and then split into three treatment groups. Mice were given ME610 (3 mg/kg) or vehicle by daily i.p. injections for 4 weeks while continuing their respective diets. n = 19-20 (males); n = 6-13 (females). **b.** Fasting blood glucose improved in ME610 versus the HFD group. Each point is an individual mouse; error bars are SEM. **c-d.** Measurement of blood glucose over time (*left*) and quantification of resulting areas under the curve (AUC, *right*) for an oral glucose tolerance test in male (*c*) and female (*d*) mice. Left panels show mean +/-SEM, and right panels show AUC for individual mice. **e-f.** As in *c-d* but with the insulin tolerance test. **g.** Mean body weights +/-95% CI for male mice over time in the DIO study. Arrow indicates when daily i.p. treatments of vehicle or ME610 began. **h.** Ǫuantification of body weights of individual mice at the end of the study. *, p < 0.05; ****, p < 0.001 by one-way ANOVA, followed by Tukey’s test for multiple comparisons. **i-j.** Body weight data from female mice, presented in the same way as g-h. **k-l.** Plots of change in body weight over the course of the DIO experiment. Arrows indicate the start of treatments with vehicle or ME610. Weight change was calculated by subtracting the weight of each mouse from its own starting weight. Each point is the mean weight change +/-95% CI for n = 19-20 (males) and n = 6-13 (females). In *k-l,* statistical significance of weight change over time was calculated by two-way ANOVA with repeated measures, followed by Dunnet’s test for multiple comparisons. **, p < 0.01; ***, p < 0.0001 *versus* the HFD group.

ME610 had profound anti-diabetic effects in the DIO model, with modest sex differences. HFD induced higher fasting blood glucose (FBG) levels and areas under the curve (AUC) in GTT and ITT, while ME610 significantly improved glucose and insulin sensitivities (Fig. 5b-f). In males, ME610 suppressed further weight gain, resulting in lower body weights relative to vehicle-treated mice on HFD at the end of the study (Fig. 5g-h; k). In females, ME610 did not affect body weights (Fig. 5i-j; l). ME610 also improved liver fat and adiposity in the DIO model, decreasing size and quantities of hepatic lipid droplets *versus* vehicle-treated mice on HFD (Fig. 6a-b). Liver weights were decreased by ME610 in males but not females; however, female liver weights were less affected by HFD (Figure 6c). In both males and females, ME610 reduced epididymal white adipose tissue (eWAT) and inguinal WAT (iWAT) masses relative to vehicle (Fig. 6d-e) without altering anterior subcutaneous WAT (asWAT), interscapular brown adipose tissue (iBAT), or subscapular BAT (sBAT) (Fig. 6f-h).

**Figure 6.**
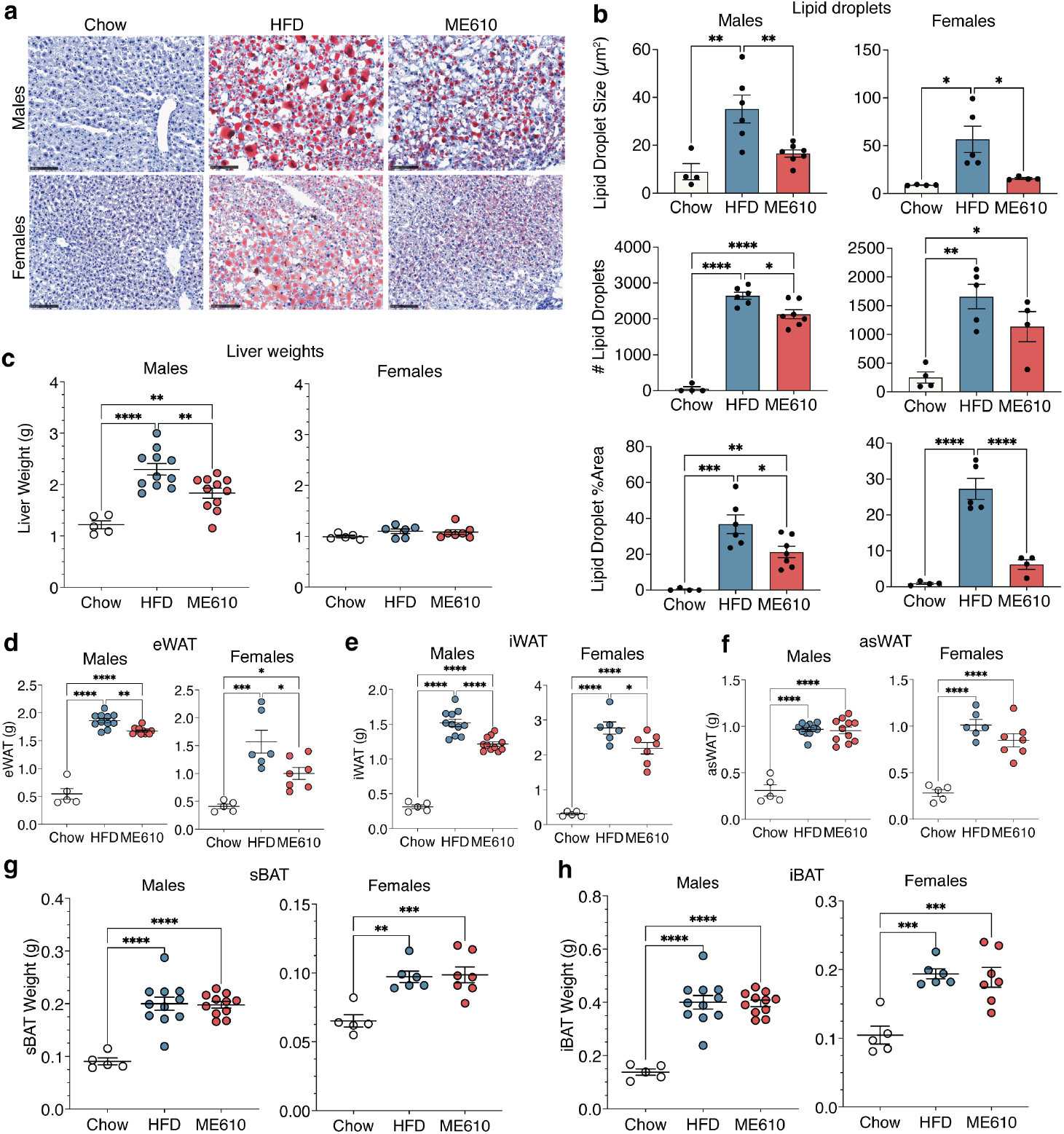
ME610 reduces intrahepatic fat and white adipose tissue mass. **a.** Representative sections from liver tissue in each group from male and female *Nr5a2^humLBD^* mice at the end of the DIO study. Sections were stained with Oil Red O (ORO). **b**. Ǫuantification of ORO staining in males (*left*) and females (*right*). n = 4 Chow; 5 HFD; 4-7 ME610. *Top,* lipid droplet size in individual mice. Bars are average +/-SEM for each group. *Middle,* lipid droplet count. Bars are average +/-SEM for each group. *Bottom,* mean lipid droplet size, expressed as percent of total area. Bars are average +/-SEM for each group. **c.** Post-treatment liver weights in males (*left*) and females (*right*). **d-g.** Post-treatment weights of adipose tissue in males (*left*) and females (*right*): eWAT (*d*), iWAT (*e*), asWAT (*f*), sBAT (*g*) and iBAT (*h*). *p < 0.05; **p < 0.01; ***p < 0.001; ****p < 0.0001 by one-way ANOVA with Tukey’s test for multiple comparisons.

### ME610 reverses DIO-induced metabolic programs and reduces hepatic triglycerides

To investigate metabolic changes associated with efficacy, we conducted transcriptomics with liver tissue from mice in the DIO study, as well as untargeted lipidomics and targeted metabolomics in liver and serum. Sex-specific expression patterns were observed in all platforms, including among chow-fed mice (Extended Data Fig. 5a-c; Extended Data Fig. 7a). Primary pairwise comparisons were HFD versus chow (“HFD-C”) to identify changes associated with diet, and HFD+ME610 versus HFD+vehicle (“ME610-H”) to identify treatment-associated changes in the context of DIO.

There is a strong signature of ME610 reversing HFD-induced metabolism across the ‘omics platforms. In RNA-seq, expression changes for ME610-H at FDR < 0.1 are mild (Extended Data Fig. 5d-e), but there are significant shifts in functionally related gene sets. KEGG and GSEA analyses show multiple metabolic pathways enriched in opposite directions for HFD-C versus ME610-H in categories related to amino acid, lipid, one-carbon/folate, and energy metabolism, and in PPAR signaling and peroxisome-related pathways (Fig. 7a; Extended Data Fig. 5f). The reversal predominantly reflects a return to baseline, as the enrichment is often lost in the ME610-C comparison. Categories related to ribosome and translation follow a different pattern, suppressed by HFD-C and further suppressed by ME610. Oxidative phosphorylation (OXPHOS), thermogenesis, and ROS-related functions are not enriched in HFD-C but are suppressed for ME610-H, with stronger effects in males than females. Additionally, there is positive enrichment of TNFα and NF-κb signaling pathways for ME610-H that is lost in ME610-C, suggesting normalization of these pathways.

**Figure 7.**
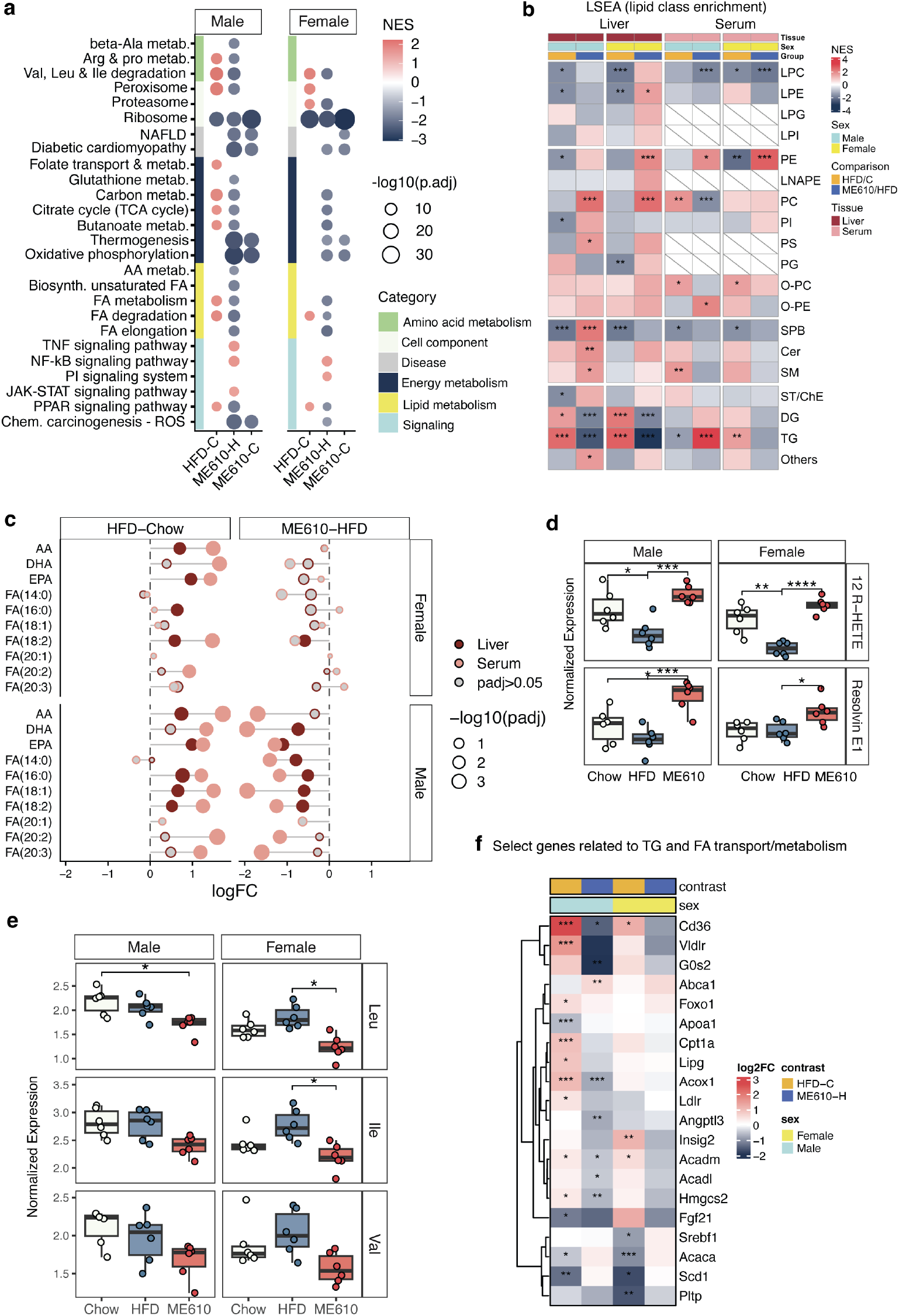
ME610 reverses HFD-induced metabolic pathways. **a.** Liver tissue from the DIO study was analyzed by RNA-seq (n = 5 males and n = 5 females from each group). Dot-plot from select, enriched pathways from KEGG analysis (p. adjust < 0.05) illustrating trends of ME610 reversing HFD-induced gene expression changes to amino acid, fatty acid, and energy metabolism. Dot color reflects NES, where red is positive enrichment and blue is negative. Dot size is inversely proportional to p.adjust. Comparisons include HFD-C (HFD group versus chow group); ME610-H (mice on HFD given ME610 versus HFD group); ME610-C (mice on HFD given ME610 versus chow group). Side annotation shows a manually assigned category. **b.** Heatmap showing LSEA enrichment from liver and serum lipidomics analysis of liver and serum from the DIO study. Comparisons include HFD-C and ME610-H in male and female mice (colors in top annotation). Heatmap is colored by NES. *, p.adjust < 0.05, **, p. adjust < 0.01, ***, p. adjust < 0.001. Cells with diagonal grey lines indicate categories with insufficient data to calculate enrichment in serum. **c.** Lollipop plot showing changes in individual fatty acids in females (*top*) and males (*bottom*). X-axis show the Log_2_ fold changes for comparisons of HFD-C (*left*) and ME610-H (*right*). Sizes of circles reflect significance. Circle color represents the tissue (dark color is liver, light is serum). Grey-filled circles did not meet the threshold for statistical significance (p. adjust < 0.05). **d.** Levels of fatty acid metabolites 12-R-HETE and resolvin E1 in the liver. Each point is a mouse in the study, boxes show the first through third quartiles with a line at the median, and whiskers show the range. *, p < 0.05; **, p < 0.01; ***, p < 0.001, ***, p < 0.0001; one way ANOVA. **e.** Levels of BCAA leucine (Leu), isoleucine (Ile), and valine (Val) in liver tissue from metabolomics data. Plotting and statistics as in *d.* **f.** Heatmap from RNA-seq showing changes to select genes related to VLDL, TG, and FA transport and metabolism. Colors represent log2 fold changes for comparisons of HFD-C and ME610-H in male and female mice. *, FDR < 0.2; **, FDR < 0.1; ***, FDR < 0.05.

Lipidomics and metabolomics data from the liver reflect pathways and HFD-reversal pattern from RNA-seq. Hepatic TGs and DGs are increased in HFD-C and reduced by ME610 (Fig. 7b; Extended Data Fig. 5h). Ratios of DG/TG and MG/DG are increased in ME610-H, suggestive of TG hydrolysis and consistent with reduced hepatic lipid droplets in ME610-treated mice (Extended Data Fig. 5i). Hepatic FFA species increase for HFD-C, including arachidonic (AA), eicosapentaenoic (EPA), palmitic (16:0), oleic (18:1), and linoleic (18:2) acids. ME610 reduces several of these, especially in males (Fig. 7c, dark circles). Concomitant increases of 12 R-HETE and Resolvin E1 suggests ME610 promotes conversion of AA and EPA to these bioactive metabolites (Fig. 7d). Branched chain amino acids (BCAA) are downregulated in ME610-H, reversing the tendency to increase in HFD-C (Fig. 7e). PCs increase for ME610-H, but the PC/PE ratio, associated with ER-stress related to cell membrane fluidity^46^, is unchanged (Extended Data Fig. 5i). In the metabolomics data, enriched pathways for ME610-H overlap substantially with GSEA/KEGG results from RNA-seq and include one-carbon, lipid, and amino acid metabolic processes (Extended Data Fig. 6).

Serum lipidomics reveals prominent effects on TGs and FA in response to diet and ME610, with some sex differences. For both sexes, TG species with total carbon chains of 48-56 and 0-4 double bonds tend to increase for ME610-H, with greater increases in males than females (Extended Data Fig. 7). Many longer-chained, more unsaturated TGs decrease for ME610-H, with stronger reductions in females than males. As a result, LSEA shows opposite enrichment patterns for circulating TGs in males and females (Fig. 7b). In males, there is negative enrichment for TGs in HFD-C and positive enrichment in ME610-H, while females show the opposite pattern, mirroring TG changes in the liver. Circulating FA increase for HFD-C to a similar degree for both sexes and are reduced by ME610, but males exhibit a stronger reduction in several FA species for ME610-H compared to females (Fig. 7c, pink circles).

The combination of lower liver TGs and higher serum TGs induced by ME610 in male mice suggests that it affects lipid transport between the liver and blood. Supporting this idea, *Cd3C* and *Vldlr*, importers of FA and VLDL, respectively, are upregulated in male mice for HFD-C and downregulated for ME610-H. (Fig. 7f). *Abca1,* involved in HDL formation and cholesterol efflux, is upregulated in ME610-H. Genes involved in VLDL transport and secretion from the liver are upregulated in HFD-C and remain upregulated in the presence of ME610 (*e.g., Erlin2*, *Rab1b,* and *Sar1b*). Additionally, ME610 downregulates *G0s2* and *Angptl3*, repressors of TG lipases ATGL and LPL, respectively^47, 48^. These ME610-induced changes trend in the same direction for female mice but often miss the threshold for statistical significance.

Collectively, the transcriptomics, lipidomics, and metabolomics data suggest ME610 reduces the burden of excess FA from HFD, easing constitutive activation of BCAA, TCA, OXPHOS, and FAO pathways. Reduction of hepatic TGs by ME610 may occur, in part, by reducing VLDL and FA import while promoting TG hydrolysis and lipid export. Reasons behind differing serum TG levels in males and females are unclear but may arise from less efficient handling of TGs in the periphery in males^49, 50^. Importantly, circulating TGs do not appear to be accumulating in WAT (Fig. 6).

### Peroxisomal lipid metabolism is associated with efficacy of ME610 in DIO

To identify the most important metabolic drivers of ME610 efficacy in the DIO study, we integrated the ‘omics datasets using two complementary approaches. First, we focused on pathways associated with the “HFD-reversal” signature from KEGG analysis. We selected genes from pathways enriched for both HFD-C and ME610-H with opposite directions in normalized enrichment scores (NES) (Fig. 7a; Table S16). Using xMWAS^51^, we performed network analysis with this subset of genes (n = 434) plus all detected lipids and metabolites. Notably, LRH-1 targets were overrepresented in the xMWAS input (47.9% *versus* 35.7% among all expressed genes, p = 3×10^-7^, Fisher’s exact test).

In the xMWAS networks (Extended Data Fig. 8a-b), genes and lipids related to peroxisome function and metabolism are prominent. Peroxisome-related genes comprise less than 10% of the input but 40-50% of the top RNA-seq nodes (Extended Data Fig. 8c). Hub genes control peroxisome biogenesis (*Pex1C*), peroxisomal FA β-oxidation (*Acox1, Decr2, Ech1, Hsd17b4*), ⍵-oxidation (*Cyp4a10*), and α-oxidation (*Ald3h2*)^52^. Other top gene nodes are related to FA transport (*Cd3C, Fabp2*) and ketone body biosynthesis (*Acss3, Hmgcl*) in males and to mitochondrial FAO (*Acadm, Eci1*) in females. Expression of these genes increase with HFD and revert to levels in the Chow group with ME610 (Fig. 8b). Among lipids and metabolites, the most central nodes consist of BCAA, tyrosine, MUFA-containing TG, and 22:6-containing PC (Extended Data Fig. 8a-b). Like the top gene nodes, these generally increase in HFD and decrease with ME610. Long-chain PUFA-containing lipids among the top nodes supports the importance of PxFAO in the HFD reversal signature.

**Figure 8.**
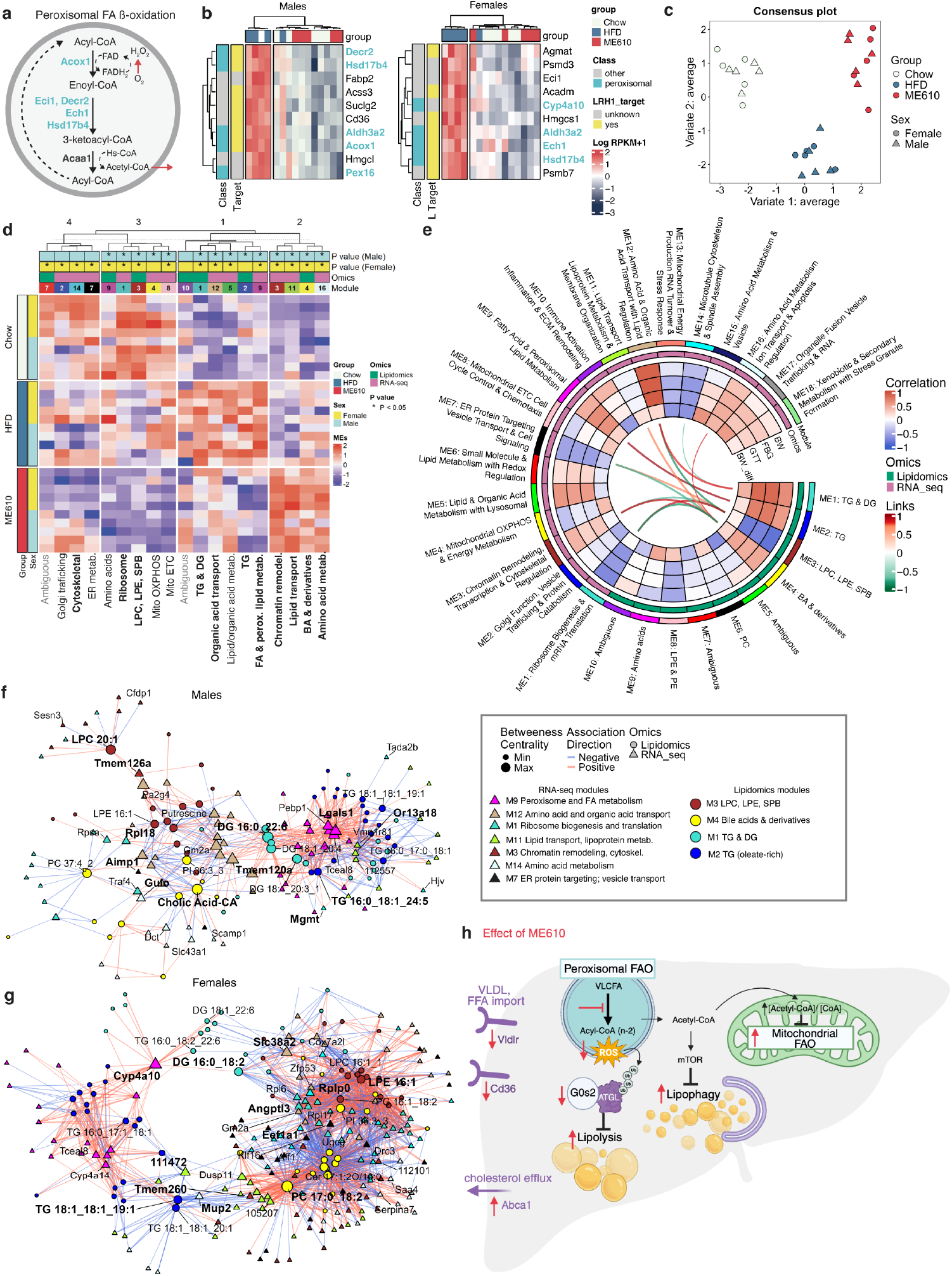
Coordinated lipid-gene expression networks underlying efficacy of ME610. **a.** Simplified diagram of the peroxisomal fatty acid pathway, with hub genes identified by xMWAS analysis highlighted in turquoise. **b.** Heatmaps of hub genes from xMWAS analysis showing a strong reversal of the HFD-induced upreguation by ME610. Yellow annotation shows genes identified as LRH-1 targets by published ChIP-seq datasets. Turquoise annotation shows peroxisome-related genes. See also Extended Data Fig. 8a-c. **c.** Consensus plot showing sample separation based on module eigengenes derived from WGCNA and integrated across datasets using the DIABLO framework in the mixOmics R package. Each point represents an individual sample, positioned according to the averaged latent variables (Variate 1 and Variate 2) from the multi-omics integration. Samples are colored by group (Chow, HFD, or 610ME) and shaped by sex (Male or Female). See also Extended Data Fig. 8d-e. **d.** Heatmap showing module eigenvalue (ME) expression patterns derived from WGCNA across samples from the Chow, HFD, and ME610 groups. Each column represents a module, and each row represents an individual sample. The color gradient ranges from red to blue, representing eigengene values from +2 (higher ME expression) to –2 (lower ME expression). Samples are annotated by group (Chow, HFD, ME610) and sex (Male, Female), while modules are annotated by their WGCNA-assigned colors. Omics layers (lipidomics vs. transcriptomics) are indicated along the top annotation bar. Hierarchical clustering was performed using Euclidean distance, allowing visualization of module-level co-expression patterns from both ‘omics datasets. GO annotations below summarize the main function(s) associated with each module (full GO results are provided in the source data). Top annotation indicates whether expression changes by group were statistically significant in males (*light blue*) or females (*yellow*) *, p < 0.05 by ANOVA. See Extended Data Fig. 8f-i for boxplots and p-values for all modules. **e.** Circular plot illustrating relationships among WGCNA MEs from lipidomics and transcriptomics, along with mouse phenotypes. The outer rings display annotated WGCNA modules, with RNA-seq modules labeled by enriched biological pathways and lipidomic modules by lipid class. Inner heatmap rings represent pairwise correlations between modules and phenotypic traits, with a color gradient from red (+1) to blue (–1) indicating correlation strength. Links in the center highlight strong cross-omics correlations between lipid and RNA-seq modules, color-coded by correlation magnitude (red for positive, green for negative) and displayed only for |r| > 0.75. Omics layers are distinguished by color-coded sidebars (lipidomics vs. RNA-seq). BW: body weight at sacrifice; BW_diff: body weight change from ME610 or vehicle treatment start until sacrifice; FBG: fasting blood glucose; GTT: Area under curve (AUC) from oral glucose tolerance test. **f-g.** Male (*f*) and female (*g*) gene–lipid correlation networks based on pairwise Pearson correlations (|r| ≥ 0.75, P < 0.01) among genes and lipid species from the top 20 ME values of high-influence RNA-seq and lipidomic modules from *e*. RNA-seq modules are shown as triangles and lipidomic modules as circles. Red and blue edges indicate positive and negative correlations, respectively. Node size reflects degree centrality, with the top three nodes highlighted. Color represents the module as indicated in the legend. **h.** Proposed model for ME610 efficacy in DIO. Suppressed peroxisomal FAO (PxFAO) reduces ROS generation, reducing oxidative stress and stabilizing ATGL by preventing ubiquitination and degradation. ATGL is further stabilized by ME610-induced suppression of *G0s2*, which binds and promotes ATGL degradation^47^. ATGL promotes lipolysis of hepatic lipid droplets. Reduced acetyl-CoA produced by PxFAO relieves inhibition of lipophagy^59^. Less acetyl-CoA imported into the mitochondria has been shown to stimulate mitochondrial FAO by reducing the acetyl-CoA/CoA ratio^61, 62^. We hypothesize that mitochondrial FAO is initially activated and then relaxes back to levels seen in chow-fed mice by the end of the study. Finally, ME610 reduces lipid burden on the liver by promoting lipid export (upregulation of Abca1 transporter) and attenuating import (downregulation of Vldlr and Cd36 transporters).

Complementing xMWAS, we performed module-based consensus network analysis across the liver ‘omics platforms using weighted gene co-expression network analysis (WGCNA). We performed GSEA on hub genes (module membership, MM > 0.7) and LSEA on hub lipids/ metabolites (MM > 0.6) to identify functions associated with modules. To visualize major expression patterns, we summarized each WGCNA module by its module eigengene (ME) and used these low-dimensional representations as input for partial least squares discriminant analysis (PLS-DA)^53^. This clearly separated treatment groups (Fig. 8c).

Group separation along variate 1 is driven by lipid M3 (enriched for LPC, LPE, and SPB), lipid M4 (bile acids and derivatives), RNA-seq M3 (chromatin remodeling, transcriptional, and cytoskeletal regulation), and RNA-seq M1 (ribosome biogenesis and mRNA translation) (Extended Data Fig. 8d-e). Variate 2 primarily separates the ME610 and Chow groups from the HFD group, resembling the HFD-reversal pattern seen across individual platforms. This variate is predominantly driven by lipid M1 (TG- and DG-enriched), lipid M2 (TG-enriched), and RNA-seq M9, associated with FA metabolism and peroxisomal lipid processing (Extended Data Fig. 8d-e). This expression pattern recapitulates xMWAS results identifying FA and peroxisomal lipid metabolism as central processes associated with reversing HFD-induced signatures.

Unbiased clustering of module eigengene (ME) values identifies four clusters with distinct expression patterns (Fig. 8e). Cluster 1 contains modules with higher ME values in the HFD group relative to Chow, with a partial reversal of the HFD effect by ME610 (includes variate 2 modules). MEs in clusters 2 and 3 include variate 1 modules and show directionally opposite responses by ME610 relative to Chow. Cluster 4 contains evidence for sex-specific differences in the ME610 group. Extended Data Fig. 8f-i compares ME values by group for all modules.

Several MEs correlate with disease outcomes in the DIO study, including post-treatment body weight (BW), FBG, and the area-under-the-curve for the GTT (Fig. 8e). RNA-seq M12 (nutrient transport and lipid regulatory signaling) shows particularly strong positive correlations with these outcomes (*r*_BW_ = 0.887, *r*_FBG_ = 0.912; and *r*_GTT_ = 0.930). Hierarchical clustering of hub genes in this module shows that diet is the major driver of expression changes in M12, since HFD and ME610 groups cluster away from the Chow group (Fig. S3). Lipid module M3 displays strong negative correlations with disease outcomes (*r*_BW_ = −0.855; *r*_FBG_ = −0.754; and *r*_GTT_ = −0.793, suggesting a protective lipid signature. This module is enriched for lysophospholipids and contains the lipid mediator oleoylethanolamide (OEA), a PPAR*a* agonist beneficial in rodent obesity models^54^. It is positively associated with RNA-seq modules related to ribosome biogenesis (M1, *r* = 0.900) and ER protein targeting/secretory functions (M7, *r* = 0.710) and negatively correlated with RNA-seq M3 (*r* = −0.723), a stress- and chromatin-associated module that contains Stat3 in its top 20 hub genes.

ME associations with treatment-related weight changes in individual mice (“BW_diff”) follow a distinct pattern relative to other disease metrics (Fig. 8e). Modules with the strongest positive associations with BW_diff include RNA-seq M10 (*r* = 0.62) and M17 (*r* = 0.57) and lipidomics module M9 (*r* = 0.49), all related to amino acid metabolism and vesicle trafficking. Modules negatively correlating with BW_diff include RNA-seq modules M14 (microtubule and cytoskeletal-related*, r =* -0.65) and M13 (mitochondrial energy production and stress response, *r* = - 0.62). The prominence of vesicle-related processes among these modules is striking.

In addition to following the HFD-reversal pattern, RNA-seq M9 (FA and peroxisome-related) positively correlates with phenotypic measurements (*r*_BW_ = 0.46, *r*_FBG_ = 0.66; and *r*_GTT_ = 0.62, *r_BW_diff_ =* 0.23) and to lipid module M2 (TG-enriched; *r =* 0.99, Fig. 8f). Hub TGs in lipid M2 contain mainly C16-C18 fatty acyl chains and 84% contain oleate, strongly reflecting the HFD diet composition (36% oleate). This relationship supports the idea that ME610 reduces the diet-induced burden on the FAO/peroxisomal machinery.

To characterize sex-specific gene-lipid interaction patterns in the DIO study, we constructed integrative molecular networks for female and male mice based on pairwise Pearson correlations between individual genes and lipids (Fig. 8f-g). Both networks exhibit a shared correlation architecture centered on the M3 and M4 lipid modules; however, denser correlations across modules in the female network indicates stronger cross-omics integration compared to the male network. Interestingly, the M9 RNA-seq module (peroxisome and FA-related) displays sex-dependent centrality. In females, key hubs include *Cyp4a14*, *Cyp4a10*, and *Vnn1*, related to long-chain FA and AA-related lipid metabolism. In males, *Acss3*, *Teal8*, and *Lgals1* dominate the central network, suggesting routing toward acetyl-CoA production, transcriptional regulation, and immune-associated metabolic control. Consistent with these transcriptional differences, the top 20 hub lipids connected to the M9 module in females are primarily long-chain FAs enriched in AA metabolism, whereas male hub lipids are biased toward ketone body metabolism. Therefore, the integrative analyses identify processes associated with adverse and favorable phenotypes in the DIO study, sex-specific responses to LRH-1 agonism, and suggest that peroxisomal and FA metabolism are important for efficacy of ME610 in obese mice.

## Discussion

The dietary phospholipid DLPC ameliorates liver fat and insulin resistance in DIO models in an LRH-1-dependent manner^10^, demonstrating an important role of LRH-1 ligands in liver health. We have developed a potent, selective LRH-1 agonist that makes DLPC-like contacts near the mouth of the binding pocket (Fig. 1)^32^. The ether in ME610 increases solubility without affecting ligand binding mode or response in cellular reporter assays versus 6N-10CA, the current most potent LRH-1 agonist (Fig. 1). Therefore, ME610 is a valuable tool for in vivo studies with potential clinical utility.

The *Nr5a2^humLBD^* mouse provides an excellent model for studying ligands tailored to the human binding pocket, addressing limitations related to sequence differences in rodent Lrh-1^36, 55, 56^. Previous strategies to produce a humanlike ligand response in mice have involved overexpression of the human receptor, sometimes in the context of mLRH-1 KO^22, 27, 28, 57^. In *Nr5a2^humLBD^* mice, expression of the humanized receptor is controlled by the endogenous promoter, preserving natural expression patterns and genomic context. Minimal editing of the binding pocket likely preserves LRH-1’s DNA-protein and protein-protein interactions important for transcriptional activity.

Efficacy of ME610 in the DIO model is associated with broad effects on metabolism, including gene expression changes in lipid, one-carbon, amino acid, and energy metabolic pathways. This response is accompanied by reduced hepatic TGs, DGs, and FA, and signatures of TG hydrolysis and lipid export. Reduced or stable white adipose depot mass suggests lipids are not being redistributed to WAT. Integrative analyses identify peroxisomal and FA metabolic genes and related lipid modules as central features associated with treatment response. This supports a model in which ligand-driven LRH-1 activation eases metabolic stress in obese mice by relieving excess FA load (Fig. 8h).

The centrality of genes related to PxFAO is intriguing, as this process has not been previously associated with LRH-1. PxFAO has been proposed as an “overflow” system to compensate for increased demand on mitochondrial FAO in MASLD^58^; however, it also contributes to hepatic steatosis by inhibiting lipolysis, lipophagy^59^, and mitochondrial FAO (reviewed recently)^60^. Inhibiting Acox1, the rate-limiting enzyme in PxFAO, reduces hepatic lipid droplets, improves insulin sensitivity, and reduces weight gain in rats on HFD^61^. These effects are associated with enhanced ROS metabolism and mitochondrial FAO^61^. In ob/ob mice, inhibition of Acox1 stimulates mitochondrial FAO and attenuates steatosis and insulin resistance^62^. The prominence of PxFAO in the module associated with reversal of HFD-induced genes, TG reduction, and improved health suggests that this process is particularly important for efficacy. Alternatively (or additionally), reduced expression of PxFAO genes could be a compensatory response to lower lipid levels.

Together, these data extend the mechanistic understanding of LRH-1 function in obesity and demonstrate the potential of synthetic LRH-1 agonists in MASLD. Strengths of our approach include the multi-omics analyses and the inclusion of both males and females: sex differences were prominent, and females are understudied in LRH-1 biology. A limitation of this work is that we have not characterized off-target effects of ME610 using a knockout mouse. Previous studies have shown hepatic expression of LRH-1 is required for agonist-induced efficacy in vivo^10^, and ME610 is highly selective for LRH-1 over other receptors in vitro (Fig. 1). However, off-target effects may contribute to metabolic changes. For example, activation of the closely-related nuclear receptor SF-1 in pancreas may cause off-target beneficial effects^63^. An additional limitation is that the multiweek treatment in the DIO study produced a mix of direct and compensatory responses that makes it difficult to discern direct LRH-1 targets responsible for efficacy. Comparisons to published LRH-1 ChIP-seq show enrichment in direct LRH-1 targets (Fig. 3; 8), and network analyses identify key phenotypic drivers. Observations from these analyses will form the basis for future mechanistic studies.

## Methods

### Chemical synthesis

Detailed synthetic methods and characterization of intermediates and final products are provided in the Supplementary materials.

### Protein purification

The LRH-1 LBD was expressed and purified as described previously^31^. Briefly, BL21 (DE3) *Escherichia coli* cells were transformed with the human LRH-1 LBD (residues 299-541) with an N-terminal His_6_ tag in a pET32a vector. Cells were grown at 37 °C in lysogeny broth to an OD_600_ of 0.6. Protein expression was induced with 1 mM isopropyl β-D-1-thiogalactopyranoside for 4 hours at 30 °C. Cells were centrifuged and stored at -80 °C. The cell pellet was resuspended in lysis buffer [20 mM Tris-HCl, 150 mM NaCl, 5% glycerol, 25 mM imidazole, 0.2 mM phenylmethylsulfonyl fluoride, DNase, and lysozyme (pH 7.4)] and lysed via sonication. Protein was isolated with Ni^2+^ affinity chromatography. The human LRH-1 LBD purifies bound to bacterial phospholipids when expressed in *Escherichia coli* ^37^. We removed co-purified bacterial lipids for the LRH-1 LBD used in fluorescence polarization (FP) binding assays by incubating the protein with 4-fold molar excess of DLPC overnight at 4 °C. The LBD was then purified with size-exclusion chromatography (SEC) into assay buffer [150 mM NaCl, 20 mM Tris-HCl, and 5 % glycerol (pH 7.4)]. The LRH-1 LBD used for thermal stability assays was purified in a similar manner but was not complexed with DLPC prior to SEC purification.

LRH-1 LBD used for crystallography was incubated with TEV protease to remove the His_6_ tag and subjected to a second round of Ni^2+^affinity chromatography to separate the tag from the protein. The cleaved protein was then complexed with ME610 as described below. All purified protein was flash frozen in liquid nitrogen in aliquots and stored at -80 °C until use.

### LRH-1 binding assays

Fluorescence polarization (FP) competition binding assays were conducted as described previously^31, 64^. Briefly, experiments were conducted in 30 µL of assay buffer [150 mM NaCl, 20 mM Tris-HCl, and 5% glycerol (pH 7.4)]. 6N-FAM (10 nM/well) was incubated with LRH-1 LBD (5 nM/well). Unlabeled compounds were added at concentrations indicated in figures, with DMSO in each well held constant at 6.7% v/v. Each experiment was performed two times with four technical replicates each. Data from each independent experiment were first normalized so that the highest and lowest values corresponded to 100 and zero, respectively. Data were then fit to a one-site, fit *K*_i_ curve in Prism, assuming a probe concentration of 10 nM and probe affinity (*K*_D_) of 8.1 nM (the latter measured in forward binding experiments with the same protein batch used for the competition assays).

### Thermal shift assays

Thermal stability of LRH-1-ligand complexes was quantified using a Tycho NT.6 Nanotemper. Purified LRH-1 LBD was incubated with 4-fold molar excess of ligand or DMSO overnight at 4 °C in assay buffer [150 mM NaCl, 20 mM Tris-HCl, and 5% glycerol (pH 7.4)]. DMSO was 4.2% v/v in all samples. Complexes were centrifuged at high speed for five minutes and then loaded into capillaries. Tryptophan/tyrosine fluorescence was monitored at wavelengths 330 and 350 nm over a range of temperatures (35 °C – 95 °C). The inflection point was determined with resident analysis software. Two separate experiments were conducted in triplicate. Results are reported as differences in inflection point for each ligand-bound complex relative to the DMSO control condition.

### Cell culture

HeLa cells were purchased from Atlantic Type Culture Collection (ATCC). Cells were cultured in humidified incubators at 37 °C and 5% CO_2_, in phenol red-free MEMα + 10% charcoal-dextran treated fetal bovine serum (FBS). Cells were authenticated by ATCC and routinely tested negative for mycoplasma.

### Reporter assay

HeLa cells were seeded at ∼7500 to 10,000 cells per well in 96-well plates (white-walled, clear bottom) in MEMα + 10% charcoal-stripped FBS. Once cells reached 80 to 90% confluence, they were transfected with LRH-1 (in pCI vector, 5 ng/well), a reporter plasmid with an NR5A response element derived from the SHP promoter cloned upstream of Firefly Luciferase (in pGL3-Basic vector, 50 ng/ well), and a plasmid expressing Renilla Luciferase under the control of the CMV promoter, used for normalization (1 ng/well). Cells were transfected with FuGENE at a ratio of 3:1 (FuGENE:DNA). Twenty-four hours after transfection, compounds (as 10 mM DMSO stocks) were diluted in Opti-MEM and added to wells to achieve final concentrations indicated in figures (final DMSO concentration was 0.37%). Luciferase signal was measured after 24 hours using the DualGlo kit (Promega) with a BioTek Neo plate reader. Firefly Luciferase signal for each well was divided by the well’s Renilla Luciferase signal intensity and then normalized relative to the DMSO control. Each experiment was conducted with three biological replicates, each with three technical replicates averaged prior to data analysis. Compiled data were analyzed with GraphPad Prism (version 10), using a three-parameter logistic function (Hill slope = 1) to calculate EC_50_.

### Solubility, metabolic stability, and pharmacokinetics

Studies were conducted by Pharmaron, Inc, using protocols detailed in the Supplementary materials.

### X-ray crystallography and structure determination

Purified LRH-1 LBD (His_6_ tag removed) was incubated with ME610 at 4-fold molar excess overnight at 4 °C. The complex was then purified via SEC into crystallization buffer (150 mM NaCl, 100 mM ammonium acetate, 1 mM EDTA, 2 mM CHAPS, and 1 mM DTT (pH 7.4)) and incubated with a peptide corresponding to human TIF2 NR box 3 (^+^H_3_N-KENALLRYLLDKDD-CO_2-_) at 4-fold molar excess, along with an additional 2-fold molar excess of ME610, for two hours at room temperature. The complex was then concentrated to ∼6 mg/mL and crystals were generated via hanging drop vapor diffusion in crystallant containing 0.1 M tri-sodium citrate (pH 4.6), 14% tert-butanol, and 5% glycerol at 4 °C. Data were collected remotely from the Southeast Regional Collaborative Access Team (SER-CAT) at the Advanced Photon Source (Argonne National Laboratories, Chicago, IL). Data were processed using HKL2000 and phased with molecular replacement, using PDB entry 7JYD as the search model^65^. Structure refinement was performed with Phenix^66, 67^ and Coot^68^. Additional refinement was performed with PDB-REDO^69^. Final figures were constructed with PyMOL (Schrödinger, LLC), which was also used to identify hydrogen bonds between the ligand and LRH-1^70^.

### Molecular dynamics simulations

*Model construction.* A total of six models were used in MDS, modeling three ligand bound states in the presence or absence of the Tif2 coregulator. Starting models comprised residues 300-538 of human LRH-1 and 742-751 of Tif2 from PDB 7TT8^31^. Because a portion of the H2-H3 loop (residues 333-337) was not resolved in the structure, we rebuilt it manually using a combination of pymol^71^ and Coot^72^. Residue K338 was deleted prior to rebuilding the loop because the sidechain occupied backbone density. To guide loop building in absence of strong electron density, we referred to a published structure with better density in this region (PDB 5L11)^26^. The structure was then refined against experimental density in Phenix using phenix.refine^66^. This input model was imported into Maestro (Schrödinger), which was used to edit the ligand to contain one or two ethers (as well as the unedited ligand 6N-10CA). The protein preparation tool in Maestro was used to cap N- and C-termini, optimize hydrogen bond assignments using PROPKA, and minimize hydrogens using the OPLS4 force field. Waters within 5Å of the ligand were retained. Identical models lacking the Tif2 peptide were also prepared by deleting the Tif2 residues from the prepared PDB files. *Simulations.* All complexes were solvated in an octahedral box of TIP3P water with a 10 Å buffer around the complex. Na+ and Cl-ions were added to neutralize the complex and achieve physiological conditions (150 mM sodium chloride). All systems were generated with the Tleap module of AmberTools24^73^, using the ff14SB force field^74^. Parameters for all ligands were obtained using the Antechamber module^75^ of AmberTools. All minimizations and simulations were performed with Amber24^76^.

Equilibration and production runs were conducted in quadruplicate for each complex. Using a minimization protocol of 5000 steps of steepest descent followed by 5000 steps of conjugate gradient, systems were minimized in an iterative manner: i) with 500 kcal/mol.Å^2^ restraints on all atoms; ii) with 500 kcal/ mol. Å^2^ restraints on restraints retained on ligand and peptide atoms; iii) with restraints removed from all atoms. Following minimization, systems were heated from 0 to 300 K using a 100-ps run with constant volume periodic boundaries and 5 kcal/mol. Å^2^ restraints on all protein and ligand atoms. Equilibration was performed using a 12-ns MD run with 10 kcal/mol. Å^2^ restraints on protein and ligand atoms using the NPT ensemble. Restraints were reduced to 1 kcal/mol. Å^2^ for an additional 10 ns of equilibration. All restraints were removed, and production runs were performed for 300 ns in the NPT ensemble. A 2-fs time step was used for all simulations. All bonds between heavy atoms and hydrogens were fixed with the

SHAKE algorithm^77^. A 10 Å cutoff was used for nonbonded interactions. Long range electrostatics were evaluated using Particle Mesh Ewald (PME). For analysis, frames were obtained at 10 ps intervals from each simulation. The CPPTRAJ module^78^ of AmberTools was used for structural averaging of MD trajectories and distance calculations. The stability of hydrogen bonds between ligands and protein was calculated with the hbond function in Amber, using a distance cutoff of 3.5Å and angle cutoff of 135 degrees. Pi-stacking was quantified by custom CPPTRAJ and python scripts. Betweenness measurements for each complex were calculated using correlationplus python package^79^. Scaled betweenness values were visualized as a heatmap and mapped onto structures for 3D visualization by substituting them into the b factor column of .pdb files for each complex.

### Animal studies

All experimental procedures were approved by the Institute for Animal Care and Use Committee at Emory University School of Medicine and conformed to the *Guide for the Care and Use of Laboratory Animals*, published by the National Institutes of Health (NIH Publication No. 86-23, Revised 1996) and with federal and state regulations. The mice were group housed by gender, allowed access to water and food *ad libitum,* and maintained under a 12-hour light–dark cycle. All mice were randomly assigned to the treatment groups.

### Generation of *Nr5a2*^humLBD^ mice

We used a CRISPR-Cas9 strategy to develop transgenic mice expressing a humanized form of LRH-1 in collaboration with the Emory Mouse Transgenic and Gene Targeting Core at Emory University. The CRISPR construct comprises a 180-bp region containing an 18-bp mutation region encoding six amino acids in the human LRH-1 ligand binding pocket (LBP) that create significant structural differences from the mouse LBP. In addition to these 18 base pairs, the construct includes two silent mutations to create the protospacer adjacent motif (PAM) sites required by Cas nuclease. The construct sequence starts in intron 5 of the mouse Nr5a2 gene sequence and continues through most of exon 6. The CRISPR donor sequence was as follows:

GGGGAACTTGGTGCTGCTGGTTCTGTTGGTGGTCACAGCCCTGATTCTGTGTTGCAGGTGGAC TACTCCACAATCATCTCACAGGCGGGCGCGACCCTGAACAATCTCCTGAGTCTCGCACAGGAGCTGG TGGTGAGGCTCCGTTCCCTTCAGTTCGATCAGCGGGAGTTTGTATGTCTC

To develop transgenic humanized mice, C57BL/6 mouse embryos were injected with Cas9 nuclease, a gRNA guide sequence, and the cloned 180-bp DNA repair template to induce cleavage and homologous DNA repair. Embryos were then implanted into pseudo-pregnant host females. Genotyping primers were designed to flank the 180-bp construct region. Mice bearing the humanized insert were identified by utilizing a novel AciI restriction cut site contained only in the insert but not the wild-type sequence. A secondary genotyping strategy involved using a third primer in the mutation site itself, thus producing a second amplicon specific to the transgenic allele. The initial CRISPR transgenic mice were effectively heterozygous, expressing both transgenic and wild-type alleles.

Each mouse was genotyped, then sequenced to ensure presence of the humanized insert in the correct genomic location. PCR products were cloned using a TOPO-TA vector and expressed in chemically competent *E. coli*. DNA from single bacterial colonies was sequenced and integrity of the humanized insert sequence was confirmed for every mouse in the first 2 generations of breeding. Some routine genotyping was outsourced to TransnetYX, Inc.

To minimize the presence and propagation of any off-target effects of CRISPR, we backcrossed 4 times to WT C57BL/6 mice. We bred the original CRISPR-injected mouse (effective heterozygote) to a WT mouse, then bred heterozygous mice from the subsequent N1, N2, and N3 generations to WT. N4 heterozygotes were then intercrossed to produce homozygous mice bearing two copies of the humanized insert. All N1 and N2 mice were sequenced as described above.

### ME610 treatment in naïve *Nr5a2^humLBD^* mice

Male *Nr5a2^humLBD^* mice (n = 6) were administered five i.p. injections of vehicle (5% DMSO in saline) or ME610, twice daily, over a span of three days. Doses given were 0.003, 0.03, 0.3, 3, and 30 mg/kg. Four hours after the last treatment, liver samples were harvested for analysis. RNA extracted from liver tissue from four mice per group in vehicle, 3 mg/kg, and 30 mg/kg groups was used for RNA-sequencing (n = 4). Lipidomics analysis was conducted using liver samples from all treatment groups (n=6).

### ME610 treatment in a DIO model

Wild-type (WT) and *Nr5a2^humLBD^* mice (male and female) were administered one of the following diets (Research Diets, Inc) starting at 8 weeks of age for 16 weeks: (1) Control Diet (10% fat diet, D12450Ji) or (2) High Fat Diet (HFD, 60% fat; D12492i). Blood glucose levels (overnight fasting) and body weights (non-fasting) were obtained prior to the start of the HFD feeding and then body weights were measured every 4 weeks until the conclusion of the feeding period. Blood obtained via a tail snip was screened using an Xtra glucose-monitoring system (Precision). At 11 weeks of HFD feeding, mice were subjected to a blood glucose test (BGT; overnight fasting). These initial readings confirmed the establishment of DIO and hyperglycemia. Mice not displaying changes in body weight (35% increase over initial weight) or hyperglycemia (i.e. 40% increase in blood glucose levels *versus* chow fed mice) were removed from the study. The mice meeting these criteria were randomly assigned to receive ME610 or vehicle (5% DMSO in saline) via a daily intraperitoneal injection starting at 12 weeks of HFD feeding and continuing for 4 weeks. At 16 weeks of HFD feeding (4 weeks of treatment), different cohorts of mice were subjected to a glucose tolerance test (GTT) or an insulin tolerance test (ITT).

For the GTT, a glucose solution (2.0 g/kg body weight in a 10% solution) was given intraperitoneally to mice that were fasted overnight. Both before and after injection (at 0,15, 30, 45, 60, 90, and 120 minutes), blood samples were obtained by tail vein clipping to determine blood glucose using an Xtra glucose-monitoring system (Precision). For the ITT, insulin (2 U/kg) was given intraperitoneally to *ad libitum* fed mice. Both before and after injection (at 0,15, 30, 45, 60, 90, and 120 minutes), blood samples were obtained by tail vein clipping to determine blood glucose using an Xtra glucose-monitoring system (Precision). Two days after the GTT or the ITT, the mice were euthanized to collect blood and tissue.

### Lipid droplet staining

Liver samples were fixed in 10% buffered formalin for 1 hour. The samples were then placed in a solution of 15% sucrose in PBS until they sank and then transferred to 30% sucrose in PBS until they sank again. Samples were then frozen in Optimal Cutting Temperature medium. Serial 5-μm liver sections from each group were stained with Oil-Red-O. Digital images of the slides were captured with a NanoZoomer (Hamamatsu) and then analyzed using ImageJ to assess the number and average size of the lipid droplets per field of view. Percent area of lipid droplet staining per field of view was also assessed.

### Western Blot Analysis

Nuclear fractions were obtained using the Nuclear Extract Kit (Abcam) according to manufacturer’s instructions. Protein concentrations were measured with the Pierce BCA protein assay (Thermo Fisher Scientific). Western blot was performed as previously described^80^. Briefly, equal amounts of protein were loaded into lanes of Criterion TGX (Tris-Glycine eXtended) Stain-Free PAGE gels (BioRad). The gels were electrophoresed and activated using a ChemiDoc MP Visualization System (BioRad). The protein was then transferred to a polyvinylidene difluoride membrane. The membranes were then imaged using a ChemiDoc MP Visualization System to obtain an assessment of proper transfer. The membranes were then blocked (3% BSA) for 1 hour at room temperature. The membrane was cut into two sections and probed with a primary antibody to LRH-1 (1:1000; ABE2867, Millipore Sigma) or fibrillarin (1:1000; 2639, Cell Signaling) overnight at 4 °C. Immunoblots were then incubated with an anti-rabbit secondary antibody (1:1000; 7074, Cell Signaling) for 1 hour at room temperature. Immunoblots were probed with a Super Signal West Dura kit (Thermo Fisher Scientific) to visualize signal, followed by visualization using a ChemiDoc MP Visualization System (BioRad). Data were analyzed using Image Lab (BioRad). The intensity of LRH-1 was normalized to the intensity of fibrillarin.

### RNA isolation

Liver tissue was snap-frozen in liquid nitrogen and stored at -80 °C prior to use. Tissue was homogenized in buffer RLT (Ǫiagen) using bead-based homogenization. RNA was purified using the RNeasy kit (Ǫiagen) with on-column DNase treatment. Purified RNA was stored at -80 °C.

### RNA sequencing

#### Dose titration study with MEC10 in Nr5a2^humLBD^ mice

Library preparation and RNA-seq for the dose-response experiment in *Nr5a2^humLBD^* mice (Figure 3) was conducted at Emory Primate Center Genomics Core. RNA quality was assessed using a TapeStation 4200 (Agilent), and then 10 ng of total RNA was used as input for cDNA synthesis using the Clontech SMART-Seq v4 Ultra Low Input RNA kit (Takara Bio), according to the manufacturer’s instructions. Amplified cDNA was fragmented and appended with dual-indexed bar codes using the NexteraXT DNA Library Preparation kit (Illumina). Libraries were validated by capillary electrophoresis on a TapeStation 4200 (Agilent), pooled at equimolar concentrations, and sequenced on an Illumina NovaSeq6000 instrument at 100SR, yielding ∼35 million reads per sample. Reads were 150 bp long, and an average of 40 million single-end reads were obtained per sample.

#### Nr5a2^humLBD^ mice vs WT and DIO study

All other RNA-seq studies were conducted by Novogene Biology Information Technology Co., Ltd. (Beijing, China). mRNA was purified from total RNA using poly-T oligo-attached magnetic beads. After fragmentation, the first strand cDNA was synthesized using random hexamer primers, followed by second strand cDNA synthesis. The library was ready after end repair, A-tailing, adapter ligation, size selection, amplification, and purification. Ǫuality check of the resulting library was performed with Ǫubit and real-time PCR (quantification) and bioanalyzer (size distribution detection). Libraries were pooled and sequenced on an Illumina Novaseq platform. Reads were 150 bp long, and an average of 20 million paired-end reads were obtained per sample. Reads were removed if they contained an adaptor, more than 10% of the bases could not be determined, or if they were under the quality threshold.

### Transcriptomics analysis

RNA-seq reads were mapped to a cDNA index constructed from the *Mus musculus* genome (assembly GRCm39) with Salmon (version 1.10.1)^81^. For the experiment with single-end reads (dose response study), options “seqBias” and “validateMappings” were selected. For studies with paired-end reads, we also used option “gcBias,” as recommended in Salmon documentation. Changes in gene expression were quantified using the *EdgeR* package (version 4.2.0) in R (version 4.4.0)^82^. Lowly expressed reads were filtered from the data using custom thresholds. The *sva* package was used to assess and model batch effects in the data^83^. In the DIO experiment, there was a significant effect of cohort (mice from all groups staggered through the study on different dates), and this was used as a batch factor instead of surrogate variables identified by sva. Significance of gene expression changes was determined by likelihood ratio tests in EdgeR, using the glmLRT function, incorporating batch effects as coefficients in the linear model. Batch-corrected log_2_ RPKM +1 or log_2_ CPM +1 were used for data visualization and clustering. Assessment of statistical significance used FDR values calculated in the linear model, including in boxplots or barplots involving RPKM or CPM values.

The published dataset involving knockout of LRH-1 in hepatocytes of adult, male mice^22^ (used for comparisons in Figure 3) was re-analyzed after downloading the fastq files from the SRA database (GSE109431). Reads were mapped to the mouse genome GRCm38 using STAR (v.2.7.10b)^84^ and analyzed with EdgeR as described above. For comparisons shown in Figure 3, the log_2_ counts per million (CPM) values for each dataset were centered around 0 for the controls (WT mice in the KO dataset and vehicle-treated mice in the ME610-titration dataset.

### Enrichment analysis for transcriptomics

#### Gene ontology (GO) enrichment analysis

The input for GO analysis was the set of genes for each comparison with FDR < 0.1. Enrichment was assessed against a background universe comprised of all expressed genes in the dataset(s). The function enrichGO in the R package *ClusterProfiler* was used for analysis^85^, with p-value cutoffs of 0.05. The Benjamin Hochberg method was used to calculate adjusted p-values. *Gene set enrichment analysis (GSEA) and Kyoto Encyclopedia of Genes and Genomes (KEGG) analysis*.

For each comparison, all expressed genes were ranked by score, calculated with the formula: sign(log_2_ fold change) * -log_10_(p-value). Ranked gene lists were evaluated for ontology enrichment using the function enrichGSE (GSEA) and gseKEGG (KEGG) using the ClusterProfiler R package^85^. The Benjamin Hochberg method was used to calculate adjusted p-values (p.adj), with a p.adj < 0.05 being considered statistically significant.

### Comparisons to LRH-1 target genes from ChIP-seq

Analyzed LRH-1 ChIP-seq data from human A549 cells CRISPR-edited to express GFP-tagged LRH-1 was extracted from the ENCODE project consortium^86, 87^ (contributed by Michael Snyder, Stanford). We also extracted published LRH-1 ChIP-seq data from mouse pancreas^40^ and HepG2 hepatocarcinoma cells overexpressing flag-tagged human LRH-1^39^. Genes with promoters bound by LRH-1 in the supplementary data files from these studies were defined as LRH-1 target genes. Enrichment of LRH-1 targets in the changing genes from the dose titration study with ME610 was determined versus a background set of all expressed genes using Fisher’s exact test, followed by the Benjamin-Hochberg correction for multiple comparisons. Adjusted p-values < 0.05 were considered statistically significant.

### Weighted gene co-expression network analysis (DIO experiment RNA-seq)

Weighted gene co-expression networks were constructed using the WGCNA package to identify consensus gene modules while accounting for sex-stratified structure, following an analogous framework to the lipid network analysis. A soft-thresholding power of 12 (scale-free fit, R² = 0.901) was selected, and a signed consensus network was generated by computing component-wise consensus topological overlap matrices (TOMs) across the female and male datasets. Genes were clustered using average-linkage hierarchical clustering on the consensus dissimilarity (1 − TOM). Modules were defined using Dynamic Tree Cut (deepSplit = 3, cutHeight = 0.998, minimum module size = 60, pamRespectsDendro = FALSE), and closely related modules were merged at a cut height of 0.20 (eigengene correlation ≥ 0.80). Consensus module eigengenes (ConMEs) were then computed to summarize gene expression patterns within each module. In total, 18 consensus gene modules were identified. Module eigengenes were used for downstream association analyses with phenotypes via Pearson correlation.

### Lipidomics: dose titration study

#### Ǫuality Control and Internal Standards

Pooled quality control samples were prepared by aliquoting 120 µL of extracted liver samples into a single vial. This sample was spiked with 10 µL internal standard (Splash Lipidomix, Avanti Polar, Birmingham, AL) and extracted in the same manner as patient samples. This analytical grade standard contains odd-chain, deuterated lipids in lipid classes and ratios present in human plasma. Pooled ǪC samples were run after every 10 samples as well as at the beginning and end of the analytical run. Ǫuality control samples were used to ensure the stability of the instrument during analysis. The coefficient of variation for each identified lipid within the pooled ǪC was then calculated using a cutoff of <40%. The internal standard was used to optimize instrumental parameters, such as electrospray voltage, collision energy, and others; these parameters were held consistently over the course of analysis. Lipids in the internal standard were also used to monitor injection consistency from sample to sample.

#### Lipid extractions

Lipids were extracted from plasma using a high throughput, monophasic, methyl t-butyl ether (MtBE) based method. Using an automated pipetting and sample preparation system (Biotage Extrahera, Uppsala, Sweden), 50 µL of homogenized liver samples was loaded into preconditioned wells containing 10 µL methanol and 10 µL of internal standard, Splash Lipidomix (Avanti Polar, Birmingham, AL). To each well, 200 µL methanol containing 50 µg/mL BHT was then added and the sample was mixed by 3 up and down passes of the automated sample handling pipette. The samples were then centrifuged at 4000 rpm for 5 minutes to pellet precipitated protein. The supernatant was recovered and transferred to a separate deep well 96-well plate for extraction. To extract lipids from the supernatant, 250 µL MtBE:methanol (3:1 v/v) was added to all wells and mixed with 3 up and down passes of the automated sample handling pipette. The sample plate was then centrifuged at 1000 g for 3 minutes and the supernatant filtered through a 0.25mm polytetrafluoroethylene (PFTE) filter plate (Biotage, ISOLUTE® FILTER+, Uppsala, Sweden) The recovered extract was then dried under nitrogen gas and subsequently reconstituted to 200 µL in acetonitrile:isopropanol (1:1 v/v) methanol for LC/MS analysis.

#### Chemicals and lipid standards

Chemical reagents and HPLC grade organic solvents were purchased from Sigma Aldrich (St. Louis, MO, USA). These include HPLC grade Water, Methanol, Acetonitrile (ACN), Isopropanol (IPA), Methyl tert-Butyl Ether (MtBE), Methyl Formate, Hexane, Ethyl Acetate, Butylated Hydroxytoluene, Ammonium Acetate, Ammonium Formate, and Formic Acid. Analytical grade, synthetic lipid standard, SPLASH Lipidomix, was purchased from Avanti Polar Lipids (Alabaster, AL, USA). PGE2 Ethanolamide (PGE2-EA), Oleoyl Ethanolamide (OEA), Palmitoyl Ethanolamide (PEA), Anandamide (AEA), Docosahexaenoyl Ethanolamide (DHEA), Linoleoyl Ethanolamide (LEA), Stearoyl Ethanolamide, 2-Arachidonoyl Glycerol (2AG), Docosatetraenoyl Ethanolamide (DEA), alpha-linolenoyl ethanolamide (ALEA), 9Z-octadecenamide (oleamide), dihomo-gamma-linolenoyl ethanolamide, docosanoyl ethanolamide, (±)9,10-dihydroxy-12Z-octadecenoic acid (9,10 DIHOME), prostaglandin E2 (PGE E2), 20-hydroxy-5Z,8Z,11Z,14Z-eicosatetraenoic acid (20-HETE), (±)-9-hydroxy-5Z,7E,11Z,14Z-eicosatetraenoic acid (9-HETE), (±)14,15-dihydroxy-5Z,8Z,11Z-eicosatrienoic acid (14,15 DHET), (±)5-hydroxy-6E,8Z,11Z,14Z-eicosatetraenoic acid (5 HETE), 12R-hydroxy-5Z,8Z,10E,14Z-eicosatetraenoic acid (12 R-HETE), (±)11,12-dihydroxy-5Z,8Z,14Z-eicosatrienoic acid (11,12-DHET), (±)8,9-dihydroxy-5Z,11Z,14Z-eicosatrienoic acid (8,9-DHET), (±)5,6-epoxy-8Z,11Z,14Z-eicosatrienoic acid (5,6 EET), (±)5,6-dihydroxy-8Z,11Z,14Z-eicosatrienoic acid (5,6-DHET), Thromboxane (TXB2), (±)12(13)epoxy-9Z-octadecenoic acid (12(13)-EPOME), (±)-13-hydroxy-9Z,11E-octadecadienoic acid (13 HODE), prostaglandin F2a (PGF2A), (±)14(15)-epoxy-5Z,8Z,11Z-eicosatrienoic acid (14(15)-EET), (±)15-hydroxy-5Z,8Z,11Z,13E-eicosatetraenoic acid (15-HETE), Leukotriene B4 (LTB4), (±)8,9-epoxy-5Z,11Z,14Z-eicosatrienoic acid (8(9)-EET), and (±)11,(12)-epoxy-5Z,8Z,14Z-eicosatrienoic acid (11(12)-EET) were purchased from Cayman Chemical (Ann Arbor, MI, USA).

### Lipidomics: DIO study

#### Ǫuality Control and Internal Standards

Pooled quality control (ǪC) samples were prepared by aliquoting 10 µL of liver homogenate from each sample into separate vials. Each ǪC sample (50 µL) was spiked with 10 µL internal standard (Splash Lipidomix, Avanti Polar, Birmingham, AL) and extracted in the same manner as plasma samples. This analytical grade standard contains odd-chain, deuterated lipids in lipid classes and ratios present in human plasma. Pooled ǪC samples were run after every 10 samples as well as at the beginning and end of the analytical run. ǪC samples were used to ensure the stability of the instrument during analysis. The coefficient of variation for each identified lipid within the ǪC was then calculated using a cutoff of <50%. The internal standard was used to optimize instrumental parameters, such as electrospray voltage, collision energy, and others; these parameters were held consistently over the course of analysis. The internal standards were also used to monitor injection consistency from sample to sample.

#### Lipid extraction used for untargeted lipidomics

Lipids were extracted from liver samples using a high throughput, monophasic, methyl t-butyl ether (MtBE) based method. Using an automated pipetting and sample preparation system (Biotage Extrahera, Uppsala, Sweden), 50 µL of homogenized liver samples was loaded into a 2 mL 96-deep well plate. To precipitate proteins, 10 µL of internal standard (Splash Lipidomix, Avanti Polar, Birmingham, AL) and 200 µL methanol with 50 µg/mL BHT were added into each well, mixed thoroughly, and centrifuged at 4000 rpm for 3 min. The supernatant was transferred into a 0.25 µm polytetrafluoroethylene (PFTE) filter plate (Biotage, ISOLUTE® FILTER+, Uppsala, Sweden). Further elution was performed by sequentially adding 800 µL MtBE:methanol (3:1 v/v) and 800 µL of 100% MtBE, with centrifugation steps (4000 rpm for 3 min) in between. The recovered extract was collected and then dried under liquid nitrogen and reconstituted to 200 µL in acetonitrile:isopropanol (1:1 v/v) methanol for LC/MS analysis.

#### Untargeted Lipidomic Analysis

Extracted lipids were resolved using a 1290 Infinity II LC system (Agilent San Jose, CA, USA). To separate lipid classes, a 5 µL sample was injected onto an Accucore C18 column (100 mm x 4.6 mm, Thermo, Waltham, MA) and resolved over a 25-minute gradient using Solvent A (10 mM ammonium formate in 60% acetonitrile with 0.1% formic acid) and Solvent B (10 mM ammonium formate in 10% acetonitrile and 90% isopropanol with 0.1% formic acid). The column was heated to 55°C in a temperature-controlled column chamber, and a 0.5 mL/min flow rate was used for analysis. Eluted lipids were analyzed by a hybrid trapped ion mobility-quadrupole time-of-flight mass spectrometer (TIMS TOF Pro 2, Bruker Daltonics, Bremen, Germany) with Vacuum Insulated Probe Heated ElectroSpray Ionization source (VIP-HESI) in positive and negative ionization modes. The instrumental parameters were optimized using analytical-grade internal standards in ǪC samples, as detailed in the chemicals and lipid standards section, and were maintained consistently throughout the analysis. The untargeted lipidomics was performed using data-dependent acquisition with parallel accumulation-serial fragmentation (DDA-PASEF) based method. The LC gradient used for analysis and MS Instrumental parameters are listed in Tables S3, and S4, respectively, located in the Supplementary material. The raw data were processed using MetaboScape 2024b (Bruker Daltonics) for peak detection, lipid identification, and sample alignment. Lipids were annotated using the T-Rex 4D algorithm based on m/z, MS/MS fragmentation patterns, and collisional cross-section (CCS) values. To ensure the highest data quality, all annotated lipids were manually curated following the automated database annotation and the criteria for lipid annotation are listed in Table S5 (Supplementary material). Annotated lipids with a coefficient of variation (CV) of <50% in ǪC samples were retained for further analysis. All annotated lipids were normalized to their corresponding internal standards within each lipid class. Principal component analysis (PCA) was performed on all samples to assess data consistency and identify outliers.

### Weighted lipid co-expression network analysis

A weighted lipid co-expression network was built with R package WGCNA, and we performed consensus network analysis to identify lipid co-expression modules while accounting for sex-specific network structure. Briefly, A thresholding power of 7 was chosen (scale-free fit index, R2 = 0.899), and the signed network was generated by the component-wise average values for consensus topologic overlap matrices (TOM) across sex-stratified datasets. Hierarchical clustering was performed on the consensus dissimilarity (1−TOM) using average linkage. Modules were identified using the Dynamic Tree Cut algorithm (deepSplit = 3, cutHeight = 0.995, minimum module size = 25, pamRespectsDendro = FALSE), and highly similar modules were merged based on eigengene similarity (merge cut height = 0.25, corresponding to eigengene correlation ≥ 0.75). Consensus module eigengenes (ConMEs) were computed, ensuring consistent eigengene estimation across both sexes. After network analysis, 10 consensus modules were found. The module eigenlipid values (MEs, the first principal component of a given module) available for the corresponding sample for calculating associations between modules and phenotypes using the Pearson correlation method.

### Module-based multi-omics integration

Module-level multi-omics integration was performed using DIABLO (Data Integration Analysis for Biomarker Discovery using Latent cOmponents) implemented in the *mixOmics* R package. Consensus module eigengenes/eigenlipids (ConMEs) derived from the WGCNA analyses were used as reduced-dimensionality features to integrate coordinated lipidomic and RNA-seq expressions. A multiblock discriminant model was constructed to identify latent components that maximize inter-omic covariance while discriminating phenotypic groups. Model tuning was conducted using repeated M-fold cross-validation (validation = “Mfold”, folds = 2, nrepeat = 1), and optimal feature selection was determined by the lowest balanced error rate (BER) using the *mahalanobis.dist* metric with the lowest classification error rate (0.14 in lipidomics; 0.02 in RNA-seq for two components). Candidate feature sets were evaluated using seq(2, 8, 2) for lipidomics modules and c(1:8, seq(10, 18, 2)) for RNA-seq modules as input to test.keepX. The final DIABLO model incorporated three latent components, including 6 lipidomic modules and 11 RNA-seq modules selected on the basis of optimal performance. These components captured shared cross-omics structure across the sex-stratified datasets and enabled identification of coordinated lipid-gene module relationships. Component loadings, module network, and sample projections (PLSDA) were used for interpretation and for evaluating integrated associations with phenotypic outcomes.

### Module enrichment analysis

To determine which lipid classes were enriched within each WGCNA module, we performed over-representation analysis (ORA) using one-sided Fisher’s exact tests, with all annotated lipids serving as the background universe. Hub lipids were defined as those with high module membership (MM > 0.6 and p < 0.05). For each lipid class, the number of lipids present in the hub-lipid set (query) and the full lipid background (universe) was tabulated, and enrichment was assessed using one-sided Fisher’s exact tests. For each class, the proportion of lipids in the query and background was calculated, and fold enrichment was defined as the ratio of % query to % background. Raw p-values were adjusted using the q-value method (False discovery rate, FDR), and lipid classes with FDR q-values < 0.05 were considered significantly enriched.

To identify Gene Ontology (GO) pathways enriched among hub genes in each RNA-seq co-expression module, a separate analysis was performed using the C5 GO gene-set collections (Biological Process, BP; Cellular Component, CC; and Molecular Function, MF) from MSigDB R package. Hub genes were defined as those with high module membership (MM > 0.7 and p < 0.05). Enrichment testing was carried out using the fora() function in the fgsea R package, with each module’s hub-gene list as the query and all detected Entrez gene IDs as the background universe. For each module, raw p-values were computed and adjusted for multiple testing using the Benjamini-Hochberg method. All enrichment analyses were performed separately for female and male networks.

### Biocrates Metabolomics Analysis

#### Overall workflow

The MxP^®^ Ǫuant 500 kit (biocrates life sciences ag, Innsbruck, Austria) was utilized for targeted metabolite quantification in 10 μl serum and 20-30 mg liver samples. The principal workflow is based on a patented 96-well plate and consists of a derivatization step, the extraction of the analytes, and the LC MS/MS carried out by ExionLC/Sciex Ǫtrap 5500 (SCIEX, Waltham, MA) LC-MS system. Sample preparation, instrument set-up, MS measurement (including system suitability testing, SST), and data processing were carried out following standard operating procedure (SOP) provided by manufacturer. Analyte-specific multiple reaction monitoring (MRM) was used for detection and quantification of 106 metabolites in the classes of alkaloids, amine oxides, amino acids, amino acids related, bile acids, biogenic amines, carboxylic acids, cresols, fatty acids, hormones and related, indoles and derivatives, nucleobases and related, vitamins and cofactors.

#### Sample preparation

After SST approval, sample preparation was carried out following the SOP. Briefly, calibrators and ǪC samples were prepared according to protocol, and 10 μl of all samples (including ǪC samples, calibrators, zero samples, and serum and homogenized liver samples) were pipetted onto the filter inserts of the 96-well plate following the predefined plate layout (1 blank, 3 zero samples (only internal standards), a set of 7 calibrators, a set of ǪCs (ǪC1, ǪC3, 5 ǪC2, and pooled sample ǪCs), and samples). The samples were dried under nitrogen for 30 min and subsequently derivatized with 50 μl phenylisothiocyanate (PITC) solution at room temperature for 60 min. Then, the derivatized samples were dried under nitrogen. Metabolites were extracted by applying 300 μl of a solution of 5 mM ammonium acetate in methanol to each well and shaking the plate for 30 min at 450 rpm. The plate was centrifuged for 2 min at 500 x g to collect the extracts through the filter inserts. Then, 150 μl extracts were transferred from the capture plate to an empty 96 deep well plate and diluted with water (1:1) for LC-MS/MS analysis.

#### LC-MS/MS

UHPLC-separations were performed on a kit-specific UHPLC column applying gradients from solvent A: 100 % water with 0.2 % formic acid to solvent B: 100 % acetonitrile with 0.2 % formic acid at 0.5 ml/min and 50 °C. The injection volume was 5 μl, the autosampler temperature was 10 °C and the total LC-MS/MS analysis time was approximately 6 min per injection.

#### Raw data processing and quality check (ǪC)

Raw data files were imported into the kit software MetIDǪ for initial data processing including retention time adjustments and peak integration review. Exported peak area data were subsequently analyzed using an in-house pipeline. Briefly, coefficient of variation (CV) of internal standards of each well, and CV of all ǪC2 were examined. All metabolites passed the CV criteria <60%. The mean value of the blank and zero samples was calculated for each metabolite as an approximation of the background noise. The limit of detection (LOD) was defined as three times this mean for each analyte. Seventy-three and 83 out of 106 metabolites were above LOD and were retained for further analysis for the liver and serum samples, respectively.

### Oxylipin Analysis

#### Sample preparation

Approximately 50 mg liver samples were homogenized in ice cold 250µL PBS using a glass bead homogenizer. 750 µL of 20% methanol was added to the homogenized samples, followed by spike BHT solution as 0.1% as the final concentration, and approximately 80 µL acetic acid to ensure the pH of the sample is around 3. The mixed samples were centrifuged at 14000rpm for 10 minutes to pellet any tissue debris. The supernatant which contains oxylipins were extracted using an automated C18 Solid Phase Extraction (SPE) manifold (Biotage Extrahera, Uppsala, Sweden). Samples were prepared by depositing homogenized samples on the SPE plate, the sample was rinsed with 800 µL water, followed by 800 µL hexane. The oxylipins were then eluted off from the SPE column with 400 µL methyl formate. The recovered oxylipin fraction was then dried under nitrogen and subsequently reconstituted in acetonitrile to be analyzed by LC/MS. Serial dilutions analytical grade standards in the linear range of 0.1 nM-10 nM were created for each standard to determine the lower limits of detection and upper limits of detection.

#### LC-MS/MS

Extracted oxylipins were resolved using an Agilent Infinity II/6495c LC-MS/MS system. 6 µL sample was injected onto an Accucore C18 column (100 x 4.6, ThermoFisher, Waltham, MA) and resolved on a 23-minute gradient using water as solvent A and acetonitrile as solvent B, both containing 10mM ammonium acetate. The column was heated to 50°C in a temperature-controlled column chamber and 0.4 mL/min flow rate was used for analysis. Eluted oxylipins were analyzed by Agilent 6495c triple quadrupole mass spectrometer in the negative and positive ion mode using an MRM based method. Instrument parameters were optimized using external analytical grade standards. See also Tables S6-S8 in the Supplementary materials for information about oxylipin quantification.

### Bile Acid Analysis

#### Sample preparation

Approximately 50 mg liver samples were homogenized in a 1:4 v/v water:methanol solution followed by shaking for 10 min at 300 rpm at 4°C and centrifuging at 10000 x g for 5 min at 4°C. The supernatants were transferred to a 96-well plate. Then, 1000uL of methanol was added to the samples prior to homogenized again and centrifuged with same condition, the supernatants were transferred to the same 96-well plate. Finally, another 500 µL of methanol was added to the previous sample, homogenized again and centrifuged with the same condition and the supernatants were combined with previous supernatant. The collected eluant was then dried with nitrogen gas and constituted in 200uL of 1:1 v/v acetonitrile:methanol and stored at -80°C until analysis. Serial dilutions analytical grade standards in the linear range of 0.1 nM-10 nM were created for each standard to determine the lower limits of detection and upper limits of detection.

#### LC-MS/MS

Chromatography was run on Agilent 6495C with 1290 Infinity II on Agilent Zorbax Eclipse Plus C18 (2.1 x 100mm, 1.8µm) column at flow rate of 0.5 ml/min at 50°C during a 23-minute gradient. The mobile phase of UPLC grade solvents consisted of solvent A: 0.1% formic acid in water and solvent B: 0.1% formic acid in acetonitrile. The injection volume of sample was 5µL. Instrument parameters were optimized using an external analytical grade standard solution: nozzle voltage at -2000V, Sheath Gas Flow at 11L/min and Sheath Gas Temp at 400°C. A total of 23 bile acids was analyzed in the negative ion mode using an MRM based method. The external standard solution was also used to optimize instrumental parameters, establish limits of detection and calibration curves.

### Targeted assay data processing and analysis

#### Data processing

Oxylipins and bile acids were quantified using Skyline software, where the area under the curve of each identified feature is calibrated against an external calibration curve. Twenty-two bile acids and 24 oxylipins with their ǪC CV <80%, median ǪC area above blank, and max sample area above 5x blank were retained for further analysis. Bile acid concentration data and biocrates data were normalized by log2 transformation followed by sample centering. Oxylipin peak area data were normalized by log2 transformation followed by feature unit variance standardization and sample centering. removeBatchEffect function from Limma R package was applied to remove the batch effect originated from different mice sac day, while preserving sex and diet effects.

#### Differential analysis

Differential analyses were performed using the normalized data separately for males and females, and for each targeted assay datasets. We ran a linear means model without an intercept (model.matrix(∼0+tr)) and performed t-tests between each contrast by running limma::contrasts.fit, and empirical Bayes variance shrinkage (limma::eBayes). P-values were adjusted using the Benjamini-Hochberg procedure to account for false discovery rate (FDR).

#### Enrichment analysis of metabolomics data

To identify biologically relevant pathways altered, pathway enrichment analysis was performed using RaMP-DB^88^. Statistically significant metabolic features (adjusted p-values < 0.05), determined via limma differential analysis, were mapped to their corresponding Human Metabolome Database (HMDB) identifiers. For each experimental contrast (stratified by sex), the list of significant analytes was subjected to over-representation analysis using the runEnrichPathways function. Pathways were considered significantly enriched if they passed an FDR-adjusted p-value threshold of 0.05. For plotting, we selected pathways containing greater than three metabolites and showing some specificity for treatment/sex combinations (*e.g.,* not significant for all 6 contrasts (group x sex)).

### xMWAS integration

The R package xMWAS (https://github.com/kuppal2/xMWAS) was used to integrate metabolomics, lipidomics, and transcriptomics data. Input data include all liver metabolites, all bile acids, all metabolites, and a subset of genes from significant KEGG pathways that exhibited inverse HFD vs ME610 effect (i.e., opposite NES in HFD_F vs ME610_F or HFD_M vs ME610_M) in either sex (n=17 pathways, see supplementary information for details). Multivariate statistical method partial least squares regression (PLS) was used where correlation threshold is 0.52 and p-value threshold is 0.05. The integration network was evaluated by eigenvector centrality (importance) and nodes with top centrality (>0.7 for females, >0.5 for males) and their edges were selected.

**Extended Data Fig. 1.**
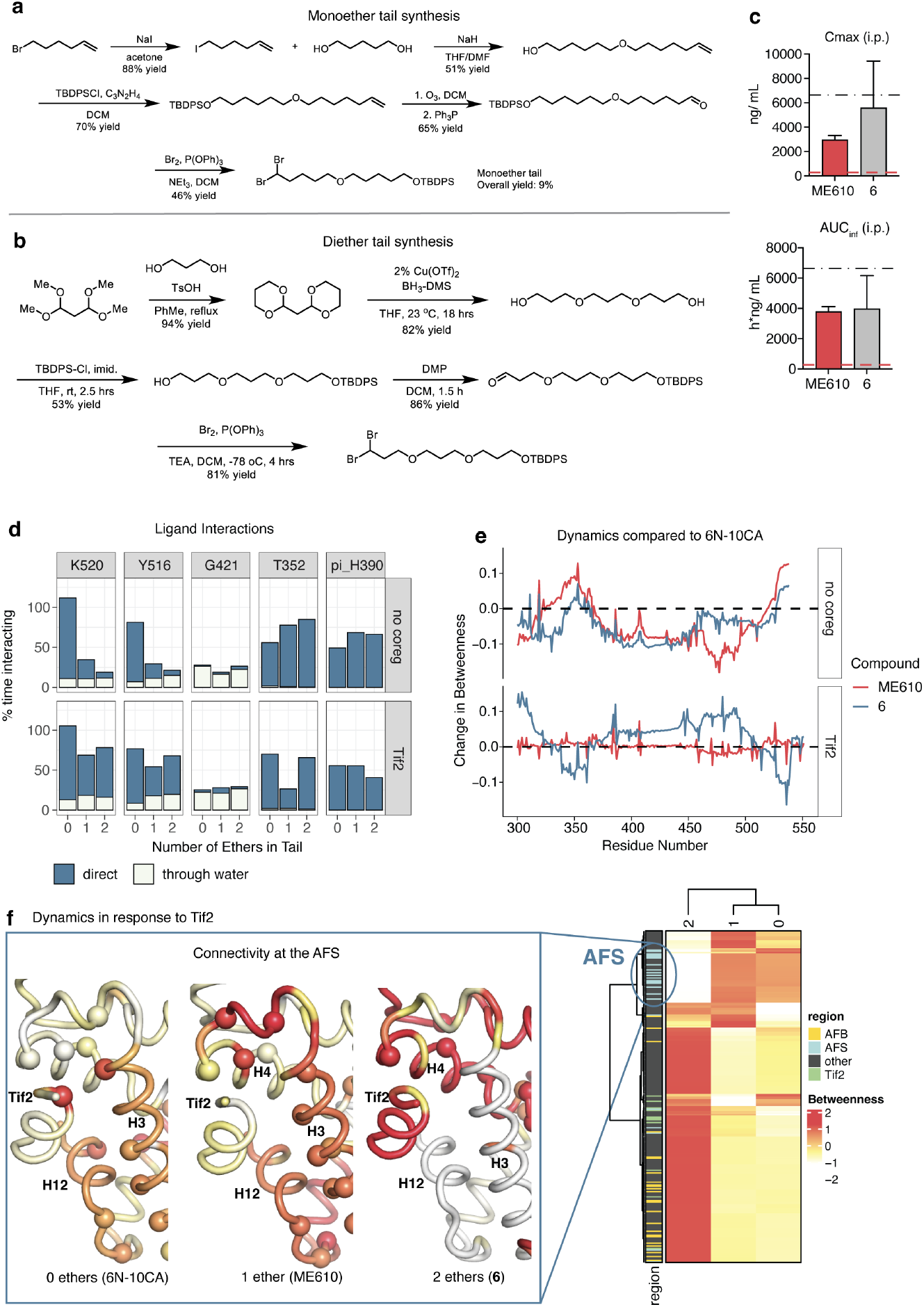
ME610 is a promising LRH-1 agonist. **a-b.** Synthetic routes for the monoether (*a*) and diether (*b*) tails used to construct compounds in this manuscript. Full synthesis and characterization are described in the supplemental methods section. **c.** Comparison of C_max_ (*top*) and AUC_inf_ (*bottom*) values in rats following a single i.p. dose of ME610 or the corresponding diether, **6** (20 mg/kg). Black dashed line shows the EC_90_ for **6** in LRH-1 luciferase reporter assays (converted to ng/mL), and red dashed line shows the ME610 EC_90_. Full PK results are presented in the supplementary materials (Table S1). **d.** Mean interaction time with key LRH-1 residues in 300 ns molecular dynamics simulations (MDS) conducted in quadruplicate. Starting models were bound to either ME610 or 6 and were run +/- the Tif2 coregulator peptide at the AFS. Blue bars represent direct interactions, and white bars water-mediated interactions. **e.** Correlation current flow betweenness values for each residue was calculated using network analysis with the correlationplus python package. Each line represents the difference in betweenness for each residue in complexes bound by ME610 (red line) or **6** (blue line) minus 6N-10CA-bound protein. When Tif2 is bound (bottom panel), the red line is level, showing equivalent betweenness to 6N-10CA-bound protein. **f.** Heatmap shows scaled per-residue betweenness values for different LRH-1-ligand complexes. Regulatory surfaces AFS (blue) and AFB (yellow) are shown in the heatmap annotation. *Inset,* close view of the AFS showing reduced connectivity in the LRH-1-Tif2-**6** complex. Scaled betweenness values were mapped onto the structure in pymol, and central nodes are shown as spheres.

**Extended Data Fig. 2.**
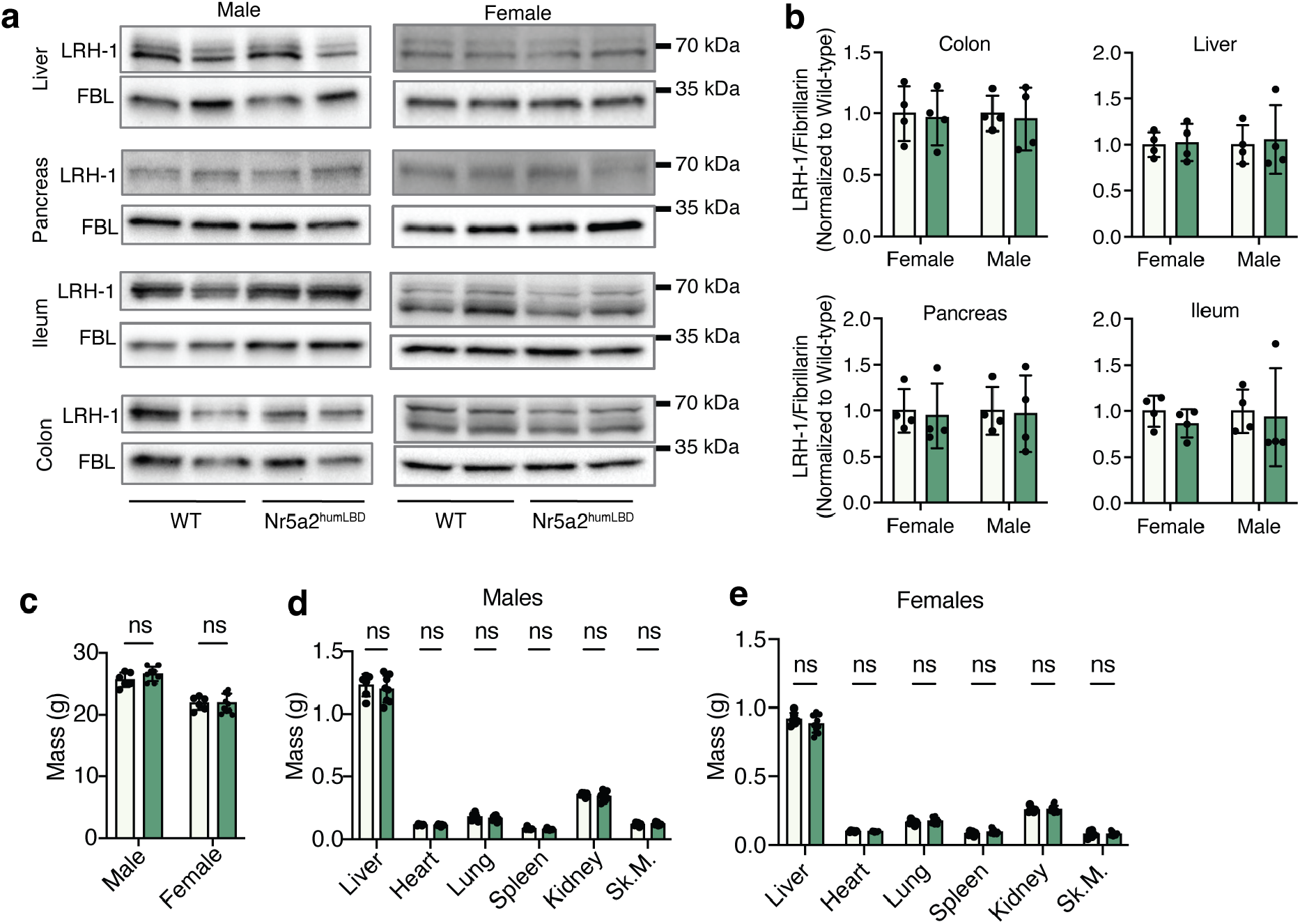
Characterization of humanized LRH-1 mice. **a.** Western Blots showing that humanizing the Lrh-1 ligand binding domain does not affect Lrh-1 protein levels in the liver, pancreas, ileum, or colon. Representative bands are shown from two male and female mice from each genotype. **b.** Ǫuantification of blots from four male and four female mice from each genotype. Lrh-1 expression was normalized to FBN. Bars represent the mean, and error bars are SD. **c.** Body weights do not differ by genotype in male or female Nr5a2^humLBD^ mice versus WT mice. Weights were measured for 10-week-old mice on a standard chow diet (n = 6-11). **d.** Organ weights do not differ by genotype in males or females.

**Extended Data Fig. 3.**
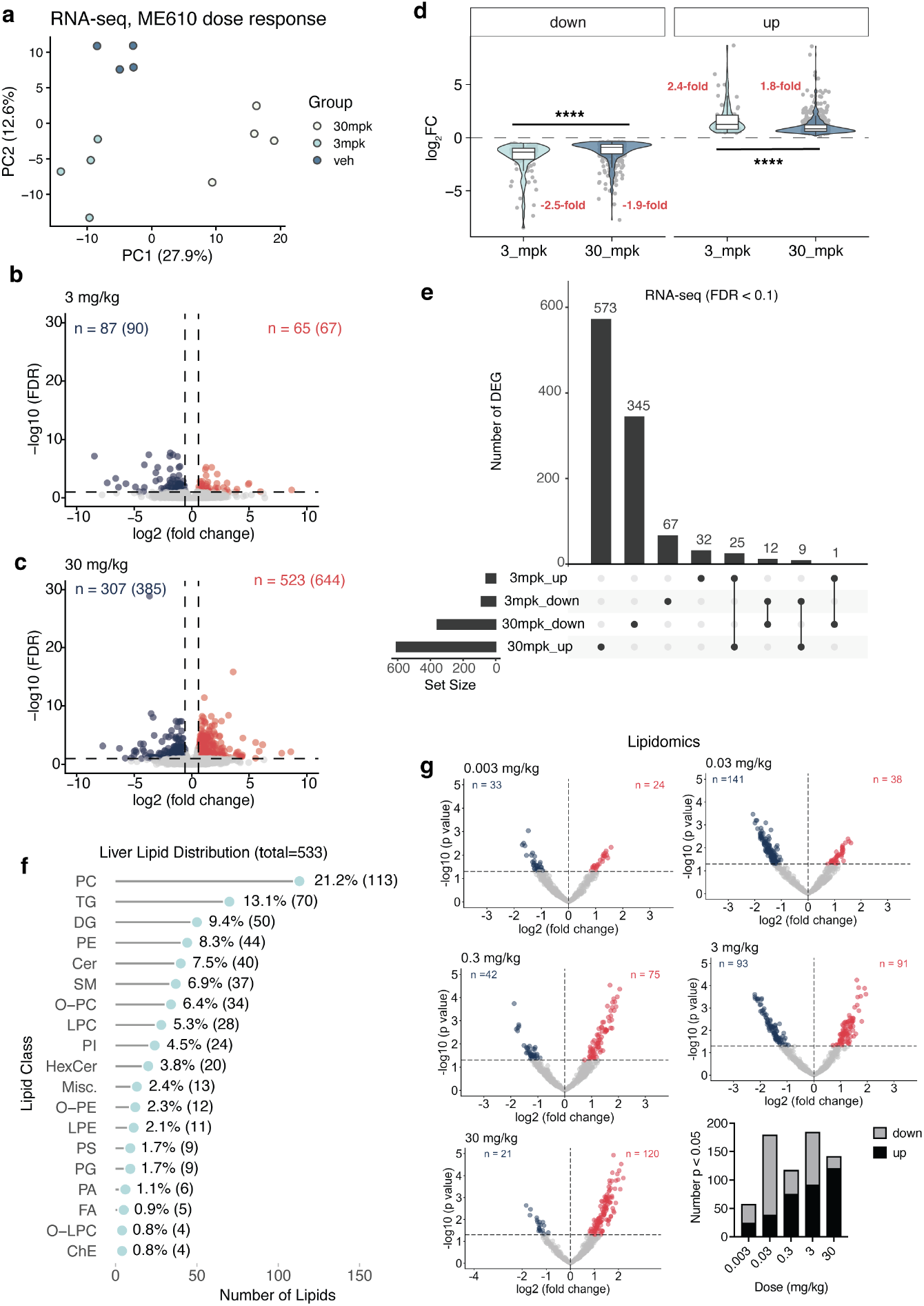
Dose-dependent effects on expression of genes and lipids by ME610 in male *Nr5a2^humLBD^* mouse liver. **a.** PCA plot showing separation by group for the top 500 most variable genes. **b-c.** Volcano plots showing distributions of differentially expressed genes (DEGs, FDR < 0.1) for 3 mg/ kg (*b*) or 30 mg/kg (c) versus veh groups. *Inset* shows numbers of DEGs. **e.** Upset plot showing overlap in DEGs for each comparison. **f.** Counts and percentages of each lipid class detected in our data. **g.** Volcano plots showing numbers and distributions of changing lipids at each dose level.

**Extended Data Fig. 4.**
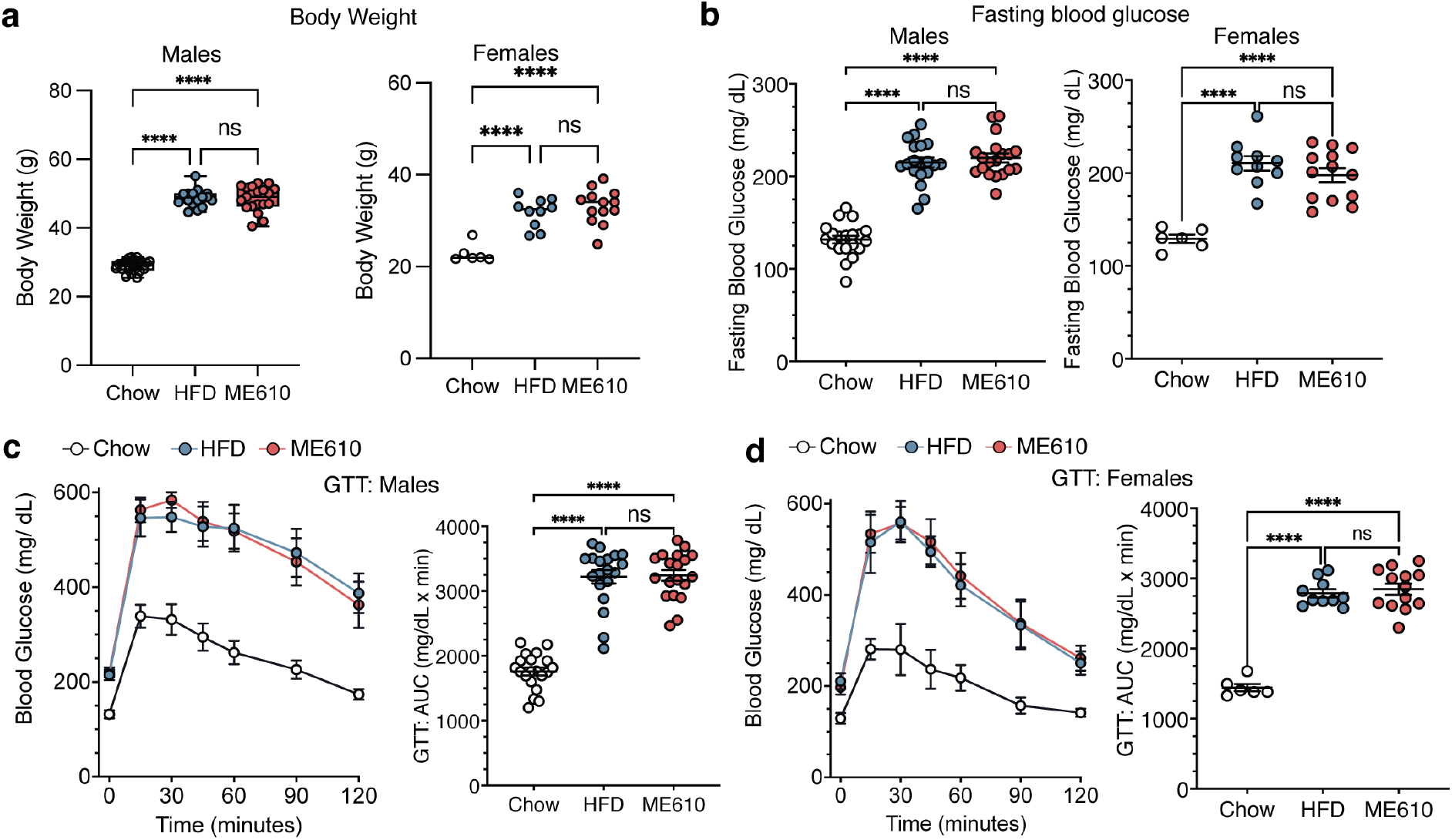
Mice partitioned into HFD and ME610 groups are phenotypically indistinguishable prior to treatment. Mice were given chow or HFD for 12 weeks; then were split into the groups shown in the x-axes labels. **a-d.** At the start of treatment with vehicle or ME610, male and female mice did not differ in body weight (*a*), fasting blood glucose (*b*), or performance in GTT (*c-d*). Values are plotted as in Fig. 5. *, p < 0.05; ****, p < 0.001 by one-way ANOVA, followed by Tukey’s test for multiple comparisons.

**Extended Data Fig. 5.**
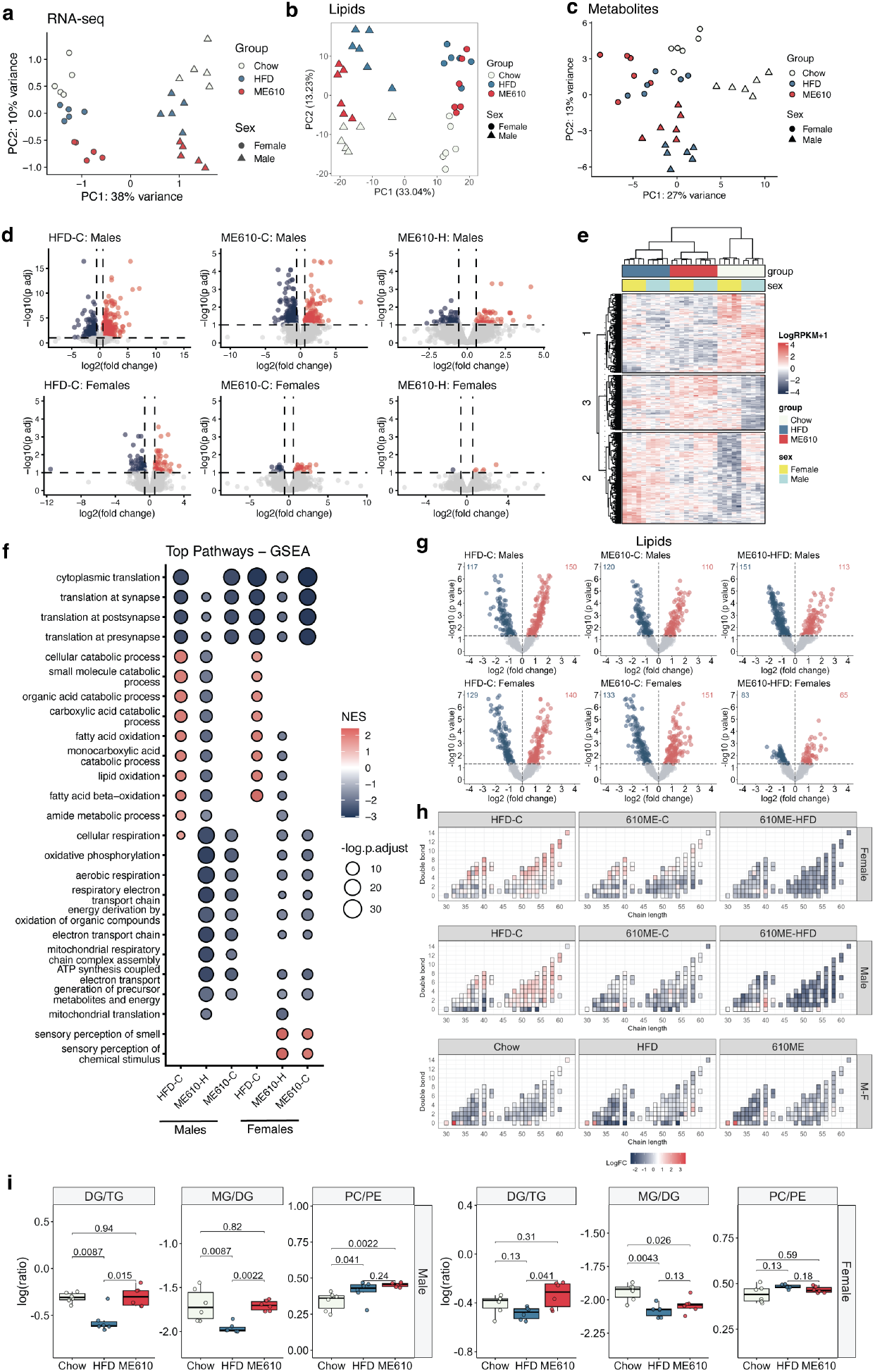
ME610 reverses metabolic pathways affected by HFD in the liver. **a-b.** PCA plots from the DIO study in RNA-seq data (*a*), lipidomics (*b*), and metabolomics (*c*) show separation by sex and treatment group across platforms. **d.** Volcano plots showing distribution of changing genes (FDR < 0.1; |fold change| >1.5) for each comparison in RNA-seq. **e.** Heatmap from the top 250 most changing genes in males and females cluster by group and show sex-specific changes, especially in the chow group. **f.** Top pathways identified by GSEA analysis from a ranked gene list from the RNA-seq study. Genes were ranked by a composite score based on p-value and direction of change (see Materials and Methods). **g.** Volcano plots showing expression changes in the lipidomics data. **h.** Distribution in hepatic TG and DG species (total chain length *versus* number of double bonds) for group comparisons, shown as a heatmap. Red indicates increased expression, and blue is decreased expression. TG (greater total chain lengths than DG) appear in the right cluster of each subpanel, and DG are in the left cluster. **i.** Ratios of related lipid species. Each point represents ratio of the lipids indicated in each subpanel for individual mice. Boxes show the interquartile range, whiskers are the range. Significance of differences was determined by one-way ANOVA. *Insets,* p-values for indicated pairwise comparisons.

**Extended Data Fig. 6.**
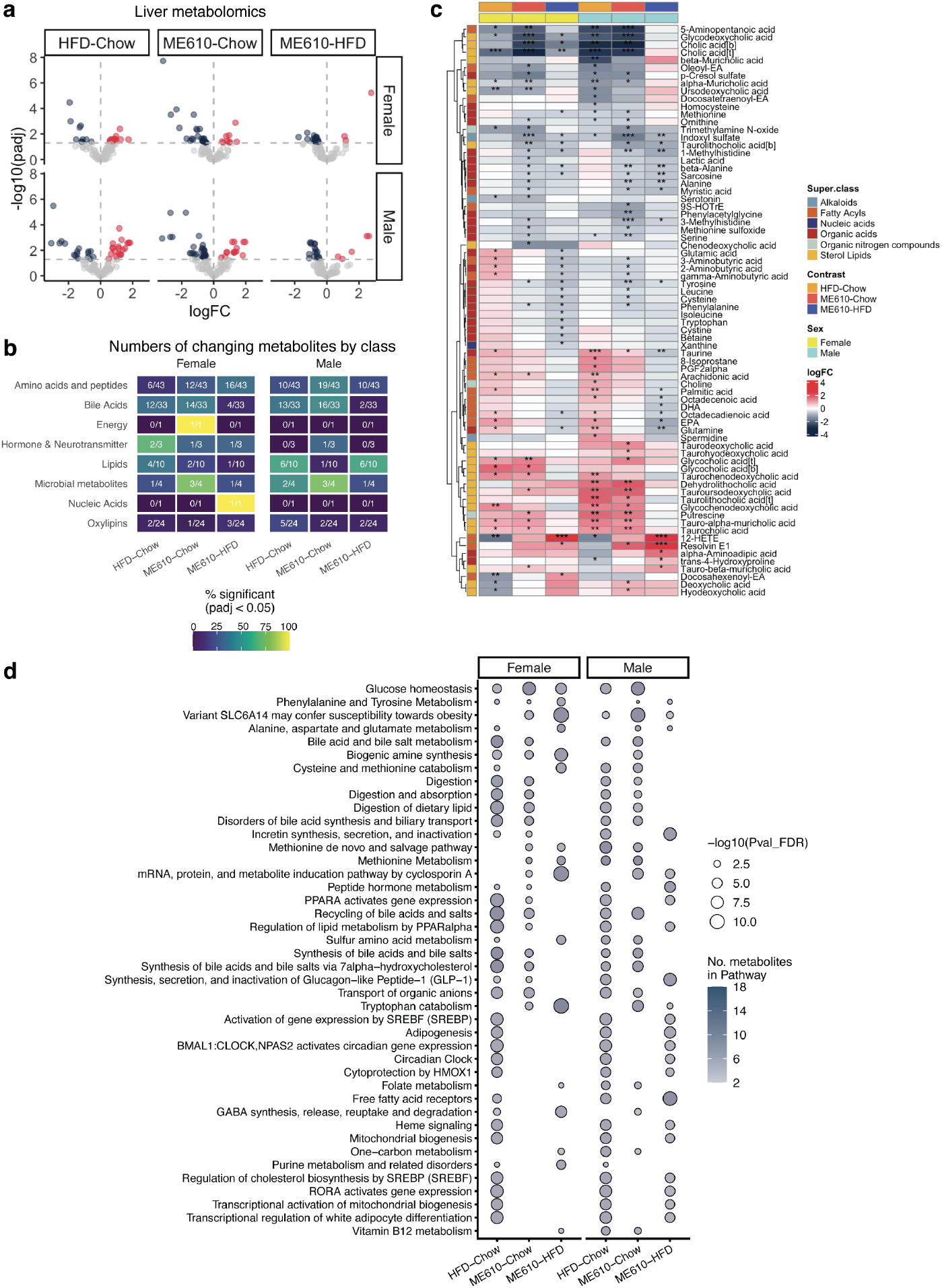
Liver metabolomics data. **a.** Volcano plots from pairwise group comparisons. **b.** Heatmap showing metabolites with statistically significant changes in at least one group. *, p < 0.05; **, p < 0.01; ***, p < 0.001 by ANOVA. **c.** Pathway enrichment among liver metabolites shows similarities to KEGG and GSEA from RNA-seq data. See Supplementary information for plots of all metabolites.

**Extended Data Fig. 7.**
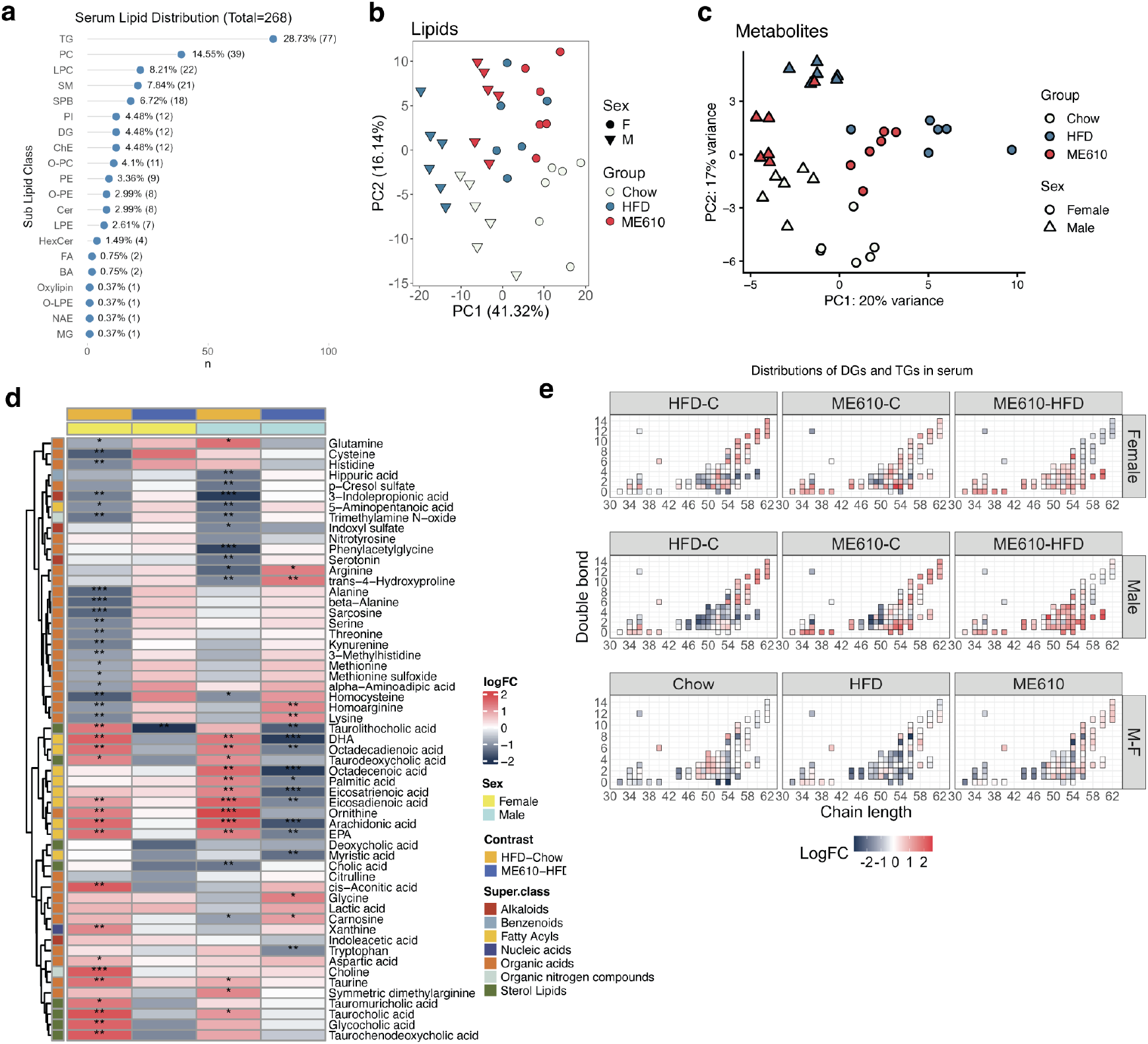
ME610 increases TGs and decreases FAs in the serum in the DIO model. **a.** Numbers of detected lipids in the serum by class. **b-c.** PCA plots from serum untargeted lipidomics (*b*) and targeted metabolomics (*c*) in the DIO experiment. **d.** Heatmap showing log2 fold changes for individual serum metabolites for each comparison. *, p < 0.05, **, p < 0.01, ***, p < 0.001 by ANOVA. Side annotation indicates the metabolite class. **e.** Heatmap of log2 fold changes of individual TGs in the DIO study. See associated source data for statistics tables.

**Extended Data Fig. 8.**
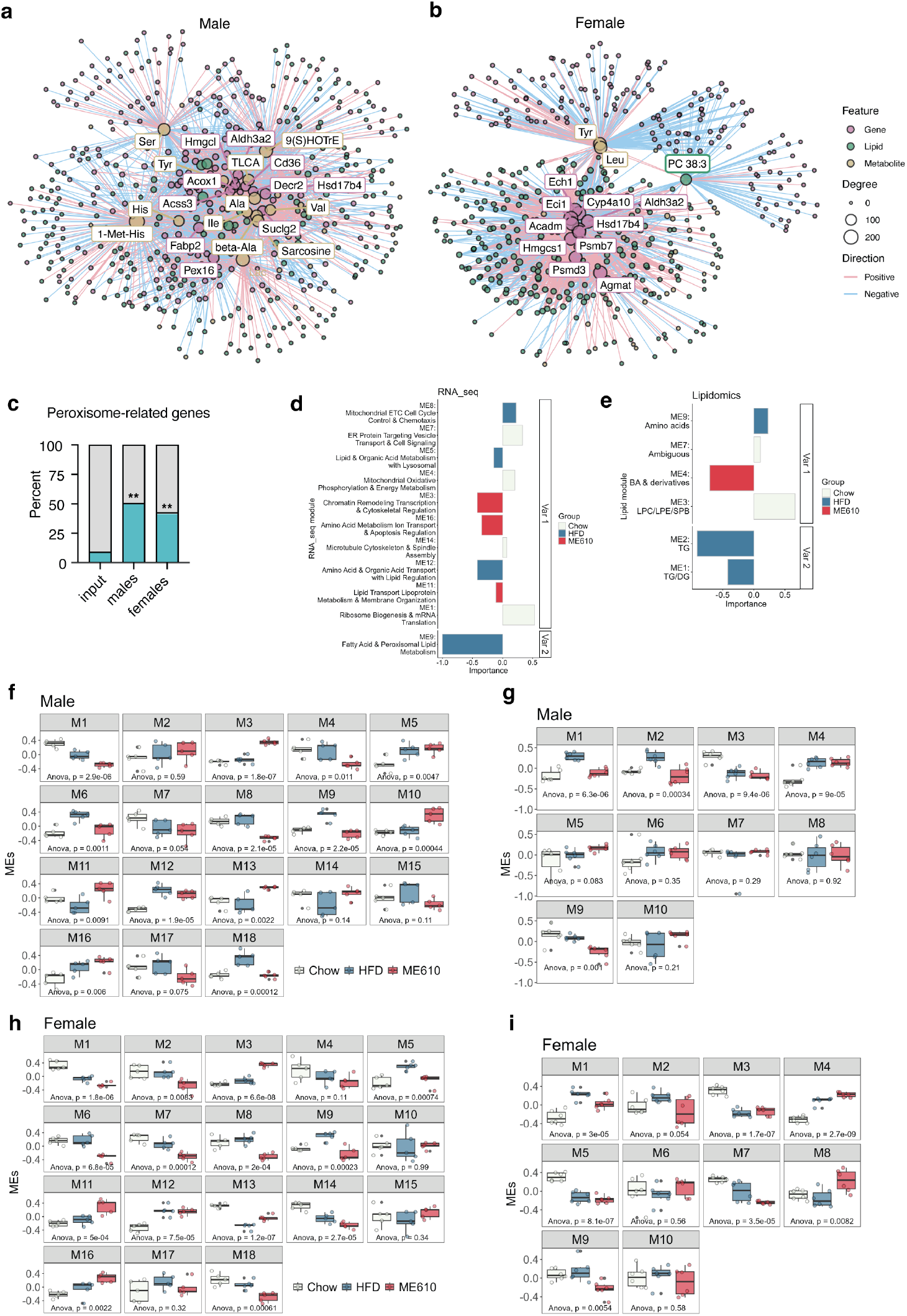
Coordinated lipid-gene expression networks underlying efficacy of ME610. **a-b.** xMWAS networks for male (*a*) and female (*b*) *NR5A2^humLBD^* mice in the DIO study. Node colors indicate the omics type and sizes indicate the degree. Edge colors represent positive or negative correlations between nodes. Top nodes are highlighted with text boxes. c. Enrichment of peroxisome-related genes among top xMWAS nodes, compared to a background set of total genes input into the analysis (input, see source data for full list of genes). Significance was determined by Fisher’s exact test, followed by the Benjamin-Hochberg correction for multiple comparisons. Adjusted p-values < 0.05 were considered statistically significant. For males, p. adjust = 0.0003; for females, p. adjust = 0.02, both versus input. **d-e.** Barplots illustrating the importance of each module in the consensus plots shown in Figure 8c. **f-i.** Plots of module eigengene values by group for RNA-seq, males (*f*), lipidomics, males (*g*), RNA-seq, females (*h*), and lipidomics, females (*i*). Each point depicts the scaled ME value for an individual animal in the DIO study. Boxes indicate the medians and interquartile ranges. Statistical significance was determined by one-way ANOVA, and p-values are displayed in each panel as insets.

## Data availability

Diffraction data from the LRH-1-ME610 crystal structure have been deposited in the Protein Databank (PDB code 36FH). RNA-seq data has been uploaded to the SRA database under the following project identifiers: PRJNA1538216 (Nr5a2^humLBD^ vs WT); PRJNA1538299 (ME610 dose response); PRJNA1538337 (DIO study). We have also provided source data for each figure. Code necessary to reproduce the figures will be shared on the Ortlund lab Github upon publication.

## Acknowledgements

The authors thank the beamline staff at Argonne National Laboratory, South East Regional Collaborative Team, for support during remote x-ray data collection.

## Ethics declarations

EAO and JWC have co-founded Allonix Therapeutics, which has licensed the molecules detailed in this manuscript. SGM, ARF, EAO, MLC, RMS, HZ, and NTJ hold patents related to LRH-1 modulators.

## Author Contributions

Conceptualization SGM, ARC, MLC, NTJ, EAO, JWC. Investigation: SGM, ARC, RMS, RP, RS, MLC, EJM, DB, HZ, ARF, SMA, AMJ. Formal analysis; SGM, ARC, MLC, CYC, ZH. Writing–original draft; SGM, ARC, CYC, ZH. All authors approve the final version of the manuscript.

