## Supplemental methods, tables, and figures for "An LRH-1 agonist reverses hepatic metabolic dysfunction in obesity"

**Table of Contents**

I. Supplementary methods
   a. Chemical Synthesis
      i. Overview and synthetic schemes
      ii. General information
      iii. General procedures
      iv. Synthesis of individual compounds
   b. Solubility and permeability assays
   c. Pharmacokinetics

II. Supplementary Tables
   a. Table S1. PK data summary
   b. Table S2. X-ray statistics for LRH-1 LBD-ME610 crystal structure
   c. Table S3. Chromatography parameters from lipidomics: Gradient time and the percentage of solvent A and B in mobile phase
   d. Table S4. Mass spectrometry instrumental parameters for untargeted lipidomic assay
   e. Table S5. The tolerances and scoring of the lipid annotation
   f. Table S6. Oxylipin chromatography gradient
   g. Table S7. Mass spectrometry instrumental parameters for targeted oxylipin assay
   h. Table S8. Oxylipins transition list

III. Supplementary Figures
   a. Figure S1. Characterization of ether-containing compounds
   b. Figure S2. *Nr5a2*$^{humLBD}$ and WT mice respond similarly to HFD
   c. Figure S3. Heatmaps from RNA-seq WGCNA modules

IV. Supplementary References

**I. Supplementary methods**

**Chemical Synthesis**

**Synthesis of monoether and diether "tails"**

The synthetic strategy involved synthesizing the fully elaborated ether-containing tails and then cyclizing each in turn. As illustrated in Extended Data Fig. 1a, the monoether tail synthesis was achieved by monoprotection of pentane diol, followed by alkylation with 6-iodo-hex-1-ene (prepared in one step from the bromide via Finkelstein reaction). The long-chain alkene was subjected to ozone and the resulting ozonide reductively opened with triphenylphosphine to furnish the aldehyde in 76% yield. Dibromination following the typical conditions was then employed; however, the purification was improved. Instead of 100% hexanes, the addition of 5% methyl *tert*-butyl ether more readily separated the desired product from triphenyl phosphate and allowed the average yield of that reaction to double (ca. 80%).

To synthesize the diether tail (Extended Data Fig. 1b), we first generated a key starting material, tripropylene glycol. While ethylene glycols of various lengths are common and commercially available, there is a dearth of literature surrounding the three carbon analogs. One report from over 50 years ago synthesized the compound by reductively cleaving a cyclic acetal, di(1,3-dioxan-2-yl)methane, with 20 equivalents of borane in refluxing THF for 5 days. While we used this method early to establish proof of concept, some optimization had to be done. After screening a series of Lewis acid catalysts, copper triflate was selected as it provided high yield (82% vs 12% for the literature conditions), overnight reaction time, and room temperature reaction. With ready access to tripropylene glycol, we then protected one of the alcohol groups (53% yield), oxidized it with DMP (86% yield), and dibrominated (78% yield) as was done with previous compounds.

The dibrominated monoether tail was subjected to cyclization and deprotection to provide the fully assembled monoether core in 72% yield (Scheme 1). Similar to the reported method for 6N-10CA[3], the diol was oxidized to the ketone-acid (58%) and then the carboxylic acid protected as the methyl ester (99%). Next, the ketone was reductively aminated utilizing ammonia and titanium[IV] isopropoxide, furnishing the primary amine in 77% yield. Following amination, the amine was stirred with sulfamide in alcoholic solvent to install the sulfamide group (74%). For compound **6,** the diether tail was then cyclized and carried through to the final sulfamide compound (14% over 6 steps, analogous to the synthetic steps for ME610 in Scheme S1).

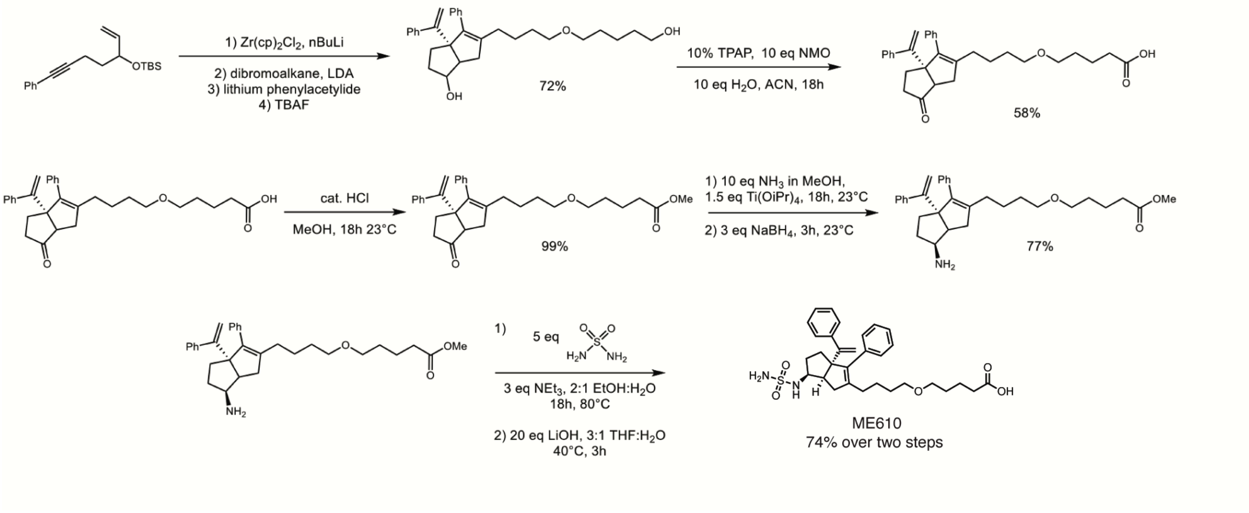

Scheme S1. Elaboration of monoether dibromo tail to ME610

**Synthesis of 3 and 5 (10CA derivatives)**

We synthesized derivatives of the cyclopentanol compound 10CA using the same ethereal dibromoalkanes, with a representative synthesis shown in Scheme S2. Cyclization of a methoxymethyl protected enyne facilitated selective deprotection of the diol. Oxidation of the free terminal alcohol to the carboxylic acid under Ley-Griffith conditions furnished the acid in moderate yield. Acidic deprotection of the methoxymethyl protecting group revealed the secondary alcohol.

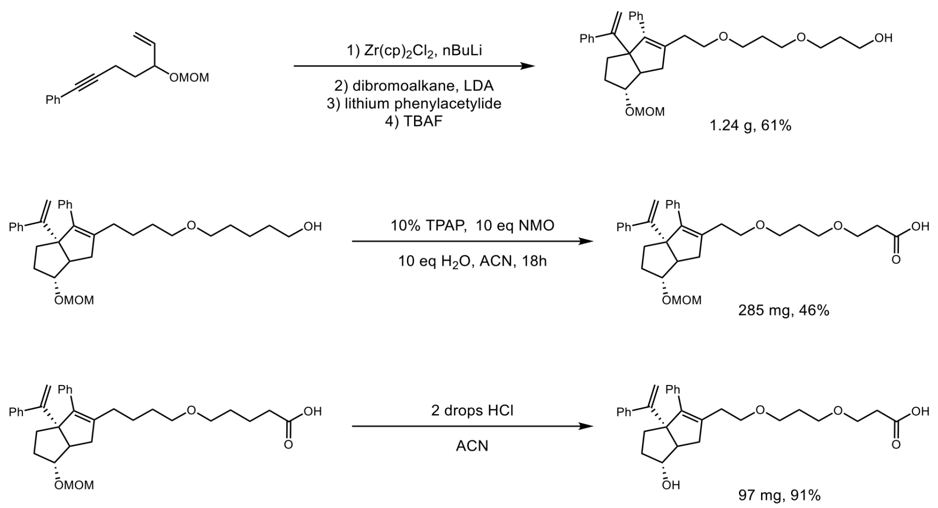

Scheme S2. Representative synthesis of 10CA derivatives with ether tails (compound **5** is shown). Syntheses of 10CA (**1**) [2] and 6N-10CA (**2**)[3] were carried out as previously described.

**GENERAL INFORMATION:**

All reactions were carried out in oven-dried glassware, equipped with a stir bar and under a nitrogen atmosphere with dry solvents under anhydrous conditions, unless otherwise noted. Solvents used in anhydrous reactions were purified by passing over activated alumina and storing under argon. Yields refer to chromatographically and spectroscopically ($^1$H NMR) homogenous materials, unless otherwise stated. Reagents were purchased at the highest commercial quality and used without further purification, unless otherwise stated. n-Butyllithium (n-BuLi) was used as a 2.5 M solution in hexanes (Aldrich), was stored at 4°C and titrated prior to use. Organic solutions were concentrated under reduced pressure on a rotary evaporator using a water bath. Chromatographic purification of products was accomplished using forced-flow chromatography on 230-400 mesh silica gel. Preparative thin-layer chromatography (PTLC) separations were carried out on 1000 μm SiliCycle silica gel F-254 plates. Thin-layer chromatography (TLC) was performed on 250 μm SiliCycle silica gel F-254 plates. Visualization of the developed chromatogram was performed by fluorescence quenching or by staining using $KMnO_4$, panisaldehyde, or ninhydrin stains.

$^{1}H$ and $^{13}C$ NMR spectra were obtained from the Emory University NMR facility and recorded on a Bruker Avance III HD 600 equipped with cryo-probe (600 MHz), INOVA 600 (600 MHz), INOVA 500 (500 MHz), INOVA 400 (400 MHz), VNMR 400 (400 MHz), or Mercury 300 (300 MHz) and are internally referenced to residual protio solvent signals. Data for $^{1}H$ NMR are reported as follows: chemical shift (ppm), multiplicity (s = singlet, d = doublet, t = triplet, q = quartet, m = multiplet, dd = doublet of doublets, dt = doublet of triplets, ddd= doublet of doublet of doublets, dtd= doublet of triplet of doublets, b = broad, etc.), coupling constant (Hz), integration, and assignment, when applicable. Data for decoupled $^{13}C$ NMR are reported in terms of chemical shift and multiplicity when applicable. Gas Chromatography Mass Spectrometry (GC-MS) was performed on an Agilent 5977A mass spectrometer with an Agilent 7890A gas chromatography inlet. Liquid Chromatography Mass Spectrometry (LC-MS) was performed on an Agilent 6120 mass spectrometer with an Agilent 1220 Infinity liquid chromatography inlet. Preparative High-Pressure Liquid chromatography (Prep-HPLC) was performed on an Agilent 1200 Infinity Series chromatograph using an Agilent Prep-C18 30 x 250 mm 10 µm column, or an Agilent Prep-C18 21.2 x 100 mm, 5 µm column.

**Evaluation of compound purity**

Purity of all tested compounds was determined by HPLC analysis, using one of the methods given below:

*Method A*: A linear gradient using water and 0.1 % formic acid (FA) (Solvent A) and MeCN and 0.1% FA (Solvent B); t = 0 min, 30% B, t = 4 min, 99% B (held for 1 min), then 50% B for 1 min, was employed on an Agilent Poroshell 120 EC-C18 2.7-micron, 3.0 mm x 50 mm column (flow rate 1 mL/min) or an Agilent Zorbax SB-C18 1.8-micron, 2.1 mm x 50 mm column (flow rate 0.8 mL/min). The UV detection was set to 254 nm. The LC column was maintained at ambient temperature.

*Method B*: A linear gradient using water and 0.1 % formic acid (FA) (Solvent A) and MeCN and 0.1% FA (Solvent B); t = 0 min, 70% B, t = 4 min, 99% B (held for 1 min), then 50% B for 1 min, was employed on an Agilent Poroshell 120 EC-C18 2.7-micron, 3.0 mm x 50 mm column (flow rate 1 mL/min) or an Agilent Zorbax SB-C18 1.8-micron, 2.1 mm x 50 mm column (flow rate 0.8148 mL/min). The UV detection was set to 254 nm. The LC column was maintained at ambient temperature.

*Method C:* A linear gradient using water and 0.1 % formic acid (FA) (Solvent A) and MeCN and 0.1% FA (Solvent B); t = 0 min, 5% B, t = 6 min, 95% B (held for 2 min), then 5% B for 1 min, was employed on an Agilent Poroshell 120 EC-C18 2.7-micron, 3.0 mm x 50 mm column (flow rate 1 mL/min) or an Agilent Zorbax SB-C18 1.8-micron, 2.1 mm x 50 mm column (flow rate 0.8 mL/min). The UV detection was set to 254 nm. The LC column was maintained at ambient temperature.

**GENERAL PROCEDURES**

*General cyclization procedure*

Hexahydropentalene formation was accomplished through slight modification of Whitby's procedure[1-4]. Prior to cyclization, all non-volatile reagents were dried by azeotropic removal of water using benzene. A dry round bottom flask containing bis(cyclopentadienyl)zirconium(IV) dichloride (1.2 equiv) under nitrogen, was dissolved in anhydrous, degassed tetrahydrofuran (THF, 8 mL/mmol enyne) and cooled to -78 °C. The resulting mixture was treated with n-BuLi (2.4 equiv) and the light-yellow solution was stirred for 45 minutes. A solution of ((5-(methoxymethoxy)hept-6-en-1-yn-1-yl)benzene) or (tertbutyldimethyl((7-phenylhept-1-en-6-yn-3-yl)oxy)silane) (1.0 equiv) in anhydrous, degassed THF (8 mL/mmol) was added. The resulting salmon-colored mixture was stirred at -78 °C for 45 minutes before the cooling bath was removed. The reaction mixture continued to stir at room temperature for an additional 2.5 hours. The reaction mixture was then cooled to -78 °C and the required 1,1-dibromoether tail (**S5** or **S10**) (1.1 equiv) was added as a solution in anhydrous THF (8 mL/mmol enyne) followed by freshly prepared lithium diisopropylamide (LDA, 1.0 M, 1.1 equiv.). After 15 minutes, a freshly prepared solution of lithium phenylacetylide (3.6 equiv) in anhydrous THF (8 mL/ mmol enyne) was added dropwise and the resulting rust-colored solution was stirred at -78 °C for 1.5 hours. The reaction was quenched with methanol and saturated aqueous sodium bicarbonate slowly warming to room temperature, affording a light-yellow slurry. The slurry was poured onto water and extracted with ethyl acetate four times. The combined organic layers were washed with brine, dried with $Na_2SO_4$, and concentrated *in vacuo*. The resulting yellow oil was passed through a short plug of silica (100% EtOAc eluent) and concentrated. The crude product was dissolved in THF and treated with either HCl or TBAF. The resulting solution stirred at room temperature for 16 h. The reaction mixture was concentrated, and the diastereomers were purified and separated by flash chromatography.

*General oxidation procedure*

To a solution starting material (1.0 equiv) in acetonitrile (0.1 M) was added tetrapropylammonium perruthenate (TPAP) (0.1 equiv), N-methylmorpholine N-oxide (NMO) (10 equiv), and water (10 equiv) and stirred at room temperature overnight. The reaction solution was then filtered through a pad of silica with (100% EtOAc) and concentrated. The resulting mixture was purified by flash chromatography to give the title compound.

*General methylation procedure*

To a solution of the desired carboxylic acid (1.0 equiv) in methanol (0.1 M) was added three drops of concentrated HCl and stirred at room temperature overnight. Reaction solution was then concentrated *in vacuo* and filtered through a pad of silica to collect the title compound.

*General reductive amination*

To a flame-dried screw top test tube charged with a stir bar backfilled (3x) was added carbonyl compound (1.0 equiv.) and ethanol (0.1 M). Ammonia (7 M in methanol, 5.0 or 20.0 equiv) then titanium(IV) isopropoxide (1.5 equiv) were added via syringe and stirred at room temperature for 6 hours. The test tube cap was then removed and sodium borohydride (3.0 equiv) added portion-wise. The resulting solution was stirred at room temperature for 30 minutes before being diluted with EtOAc. The solution was adjusted to a pH of 1 with 1M HCl. The layers were separated, and the aqueous layer was extracted 3x with EtOAc. The combined organic layers were then dried over $Na_2SO_4$, filtered, and concentrated *in vacuo* before being purified by flash chromatography to give the title compound.

*General deprotection procedure*

A reaction vial was charged with a stir bar and starting material (1.0 equiv) dissolved in dioxane and a few drops of concentrated HCl was then added. The reaction was gradually heated to 40 °C before being diluted with EtOAc and washed with 3 x 5 mL 0.5 M aqueous HCl, 5 ml water, and 5 mL brine. The organic layer was then dried over $Na_2SO_4$, filtered, and concentrated *in vacuo* to give the title compound after flash chromatography.

**SYNTHESIS OF INDIVIDUAL COMPOUNDS**

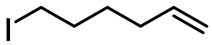

**6-iodohex-1-ene** (**S1**): To a round bottom flask under nitrogen flow was added sodium iodide (93.68 g, 5 equivalents, 625 mmol), acetone (300 mL) and 6-bromo-1-hexene (125 mmol, 1 eq, 16.75 mL). The suspension was stirred and heated to reflux for 18h. The reaction was then loaded onto a silica plug, eluted with hexanes, 1:1 hexanes and MTBE, and MTBE (75 mL each). The organics were concentrated to a colorless oil (21.9 g, 104 mmol, 83%).

**$^1$H NMR** (600 MHz, Chloroform-*d*) δ 5.76 (ddtd, *J* = 17.0, 10.2, 6.7, 1.5 Hz, 1H), 4.99 (dp, *J* = 17.1, 1.7 Hz, 1H), 4.94 (dp, *J* = 10.2, 1.6 Hz, 1H), 3.17 (td, *J* = 7.0, 1.5 Hz, 2H), 2.05 (qd, *J* = 7.3, 1.5 Hz, 2H), 1.85 – 1.76 (m, 2H), 1.48 (pd, *J* = 7.5, 1.5 Hz, 2H).

**$^{13}$C NMR** (151 MHz, Chloroform-d) δ 138.12, 115.00, 32.91, 32.62, 29.69, 6.82.

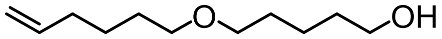

**5-(hex-5-en-1-yloxy)pentan-1-ol** (**S2**): Sodium hydride (60% in mineral oil, 3.46 g, 86 mmol, 1 eq) was added to a flame dried round bottom flask under positive nitrogen pressure followed by 115 mL of DMF and 115 mL of THF. This was cooled to 0 °C and then 1,5-pentane diol (27.2 mL, 3.0 eq, 260 mmol) was added dropwise. The resulting solution was stirred for 20 minutes, then 6- iodohex-1-ene **(S1)** was added dropwise. The reaction was stirred overnight and allowed to warm to room temperature. In the morning, saturated ammonium chloride was added to quench the reaction and then the mixture was extracted with ethyl acetate (3x 100 mL). The organics was combined, dried, and concentrated to provide a crude oil that was purified by silica chromatography (0-20% EtOAc:hexanes) to yield the title compound (8.01 g, 43 mmol, 50% yield).

**$^1$H NMR** (600 MHz, Chloroform-*d*) δ 5.80 (ddtd, *J* = 16.9, 10.1, 6.7, 0.9 Hz, 1H), 5.00 (dtd, *J* = 17.1, 2.6, 1.5 Hz, 1H), 4.94 (ddq, *J* = 10.1, 2.2, 1.1 Hz, 1H), 3.64 (td, *J* = 6.5, 0.9 Hz, 2H), 3.40 (tdd, *J* = 6.6, 3.8, 0.9 Hz, 4H), 2.06 (tdd, *J* = 7.8, 6.4, 1.3 Hz, 2H), 1.63 – 1.55 (m, 6H), 1.47 – 1.40 (m, 4H).

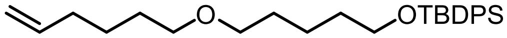

***tert*-butyl((5-(hex-5-en-1-yloxy)pentyl)oxy)diphenylsilane** (**S3**): 8.01 g of 5-(hex-5-en-1-yloxy)pentan-1-ol **(S2)** (43 mmol) was dissolved in 400 mL of dry THF, followed by 4.39 g imidazole (1.5 eq, 64.5 mmol). The mixture was stirred for 5 minutes until it was homogeneous and then TBDPS-Cl (1.2 eq, 51.6 mmol, 13.42 mL) was added dropwise. Stirred overnight at room temperature. The reaction was concentrated to approximately 150 mL and passed through a celite plug to remove imidazolium salts. The filtrate was then concentrated to a clear oil that was purified by silica chromatography (1-7% EtOAc/hexanes) (12.9 g, 30.37 mmol, 70% yield).

**$^1$H NMR** (600 MHz, Chloroform-*d*) δ 7.69 – 7.64 (m, 4H), 7.44 – 7.35 (m, 6H), 5.80 (ddt, *J* = 16.9, 10.2, 6.7 Hz, 1H), 5.00 (dq, *J* = 17.1, 1.7 Hz, 1H), 4.94 (ddt, *J* = 10.2, 2.5, 1.2 Hz, 1H), 3.65 (t, *J* = 6.5 Hz, 2H), 3.38 (dt, *J* = 9.3, 6.6 Hz, 4H), 2.10 – 2.03 (m, 2H), 1.62 – 1.51 (m, 6H), 1.48 – 1.37 (m, 4H), 1.04 (s, 9H).

**$^{13}$C NMR** (151 MHz, Chloroform-d) δ 138.83, 135.59, 134.15, 129.50, 127.59, 114.47, 70.87, 70.73, 63.89, 33.60, 32.42, 29.52, 29.26, 26.89, 25.53, 22.45, 19.23.

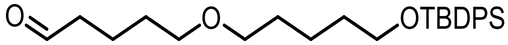

**5-((5-((*tert*-butyldiphenylsilyl)oxy)pentyl)oxy)pentanal** (**S4**): 9.59 mmol (4.06 g) of **S3** was dissolved in 150 mL of dry DCM and the reaction cooled to -78 °C. Next, ozone was bubbled through the reaction while stirring until a blue color persisted. Oxygen was then bubbled through the reaction to flush remaining ozone and then 3 equivalents (7.73 g) of triphenylphosphine was added. The reaction was warmed to room temperature and stirred for 3 hours. The solvent was removed *in vacuo* and the resulting oil purified by silica chromatography (0-20% EtOAc/hexanes). 3.083 g, 76%

**$^1$H NMR** (600 MHz, Chloroform-*d*) δ 9.76 (t, *J* = 1.8 Hz, 1H), 7.71 – 7.60 (m, 4H), 7.46 – 7.33 (m, 6H), 3.66 (t, *J* = 6.5 Hz, 2H), 3.39 (dt, *J* = 18.5, 6.5 Hz, 4H), 2.45 (td, *J* = 7.3, 1.8 Hz, 2H), 1.75 – 1.67 (m, 2H), 1.64 – 1.51 (m, 6H), 1.45 – 1.36 (m, 2H), 1.04 (s, 9H).

**$^{13}$C NMR** (101 MHz, Chloroform-d) δ 202.67, 135.58, 134.11, 129.53, 127.60, 70.96, 70.26, 63.85, 43.66, 32.40, 29.48, 29.18, 26.88, 22.44, 19.24, 19.00.

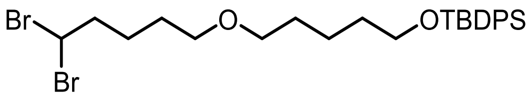

***tert*-butyl((5-((5,5-dibromopentyl)oxy)pentyl)oxy)diphenylsilane** (**S5**): Under nitrogen, a solution of triphenylphosphite (2.35 mL, 9 mmol, 1.1 equiv) in DCM (50 mL) was cooled to -78 °C. Bromine (0.48 mL, 9 mmol, 1.1 equiv) and triethylamine (1.32 mL, 9 mmol, 1.1 equiv) were sequentially added dropwise at -78 °C. The reaction was stirred for 5 minutes, then **S4** (5.8 g, 14 mmol, 1.0 equiv) was added as a solution in DCM (10 mL) via syringe at -78 °C. The reaction was stirred for 5 h and allowed to warm to ambient temperature. The whole reaction was then poured over a pad of silica. The filtrate was concentrated and purified on a short plug of silica with 5% MTBE in hexanes to afford the title compound as a clear, colorless oil (3.35 g, 72%).

**$^1$H NMR** (600 MHz, Chloroform-*d*) δ 7.64 (dt, *J* = 8.0, 1.4 Hz, 4H), 7.44 – 7.31 (m, 6H), 5.67 (td, *J* = 6.2, 1.1 Hz, 1H), 3.63 (td, *J* = 6.5, 1.2 Hz, 2H), 3.42 – 3.29 (m, 4H), 2.38 (dddt, *J* = 7.6, 6.1, 4.2, 2.0 Hz, 2H), 1.66 – 1.46 (m, 9H).

**$^{13}$C NMR** (151 MHz, Chloroform-*d*) δ 135.58, 129.51, 128.35,127.59, 70.98, 70.28, 63.87, 46.01, 45.20, 32.39, 29.48, 28.38, 26.89, 25.10, 22.45, 19.24.

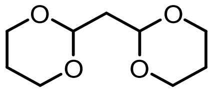

**di(1,3-dioxan-2-yl)methane** (**S6**): The title compound was prepared according to the reported procedure (17.6 g, 94%) and the NMR data were consistent with those previously reported [5].

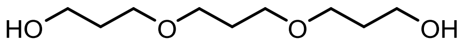

**3,3'-(propane-1,3-diylbis(oxy))bis(propan-1-ol)** (**S7**): A dry 3-neck round bottom flask charged with a stir bar was removed from a drying oven and sealed with septum stoppers before being connected to a vacuum line. The flask was allowed to cool to room temperature under vacuum. Once cool, Di(1,3-dioxan-2-yl)methane (21 g, 111.85 mmol, 1.0 equiv) and freshly prepared $Cu^{II}(OTf)_2$ (810 mg, 2.24 mmol, 2% loading) were added to the flask under positive nitrogen pressure. The flask was resealed with a septum stopper. The atmosphere was exchanged by applying vacuum and backfilling with $N_2$ (this process was conducted a total of three times). Degassed tetrahydrofuran (THF, 65 ml/mmol Di(1,3-dioxan-2-yl)methane) was delivered to the flask and the resulting solution was cooled to 0 °C. Once cool, $BH_3$·DMS (37.1 ml, 391.5 mmol, 3.5 equiv) was added. The resulting mixture was allowed to stir overnight while warming to room temperature. The reaction was quenched with MeOH and subjected to a short celite plug. The resulting crude mixture was concentrated *in vacuo* and subjected to silica gel chromatography (5-15% MeOH/EtOAc) to afford the title compound (13.3 g, 62%). The title compound's NMR data were consistent with those previously reported [6].

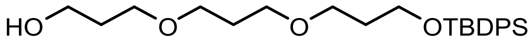

**2,2-dimethyl-3,3-diphenyl-4,8,12-trioxa-3-silapentadecan-15-ol** (**S8**): To a solution of tripropylene glycol (13.3 g, 69.2 mmol, 1.3 equiv) and imidazole (3.62 g, 53.2 mmol, 1 equiv) in anhydrous THF (800 ml) in an oven-dried flask under nitrogen at r.t. was added *tert*-butyl(chloro)diphenylsilane (13.85 ml, 53.2 mmol, 1 equiv) and stirred for 2.5 h. The resulting solution was concentrated *in vacuo* and subjected to silica gel chromatography (20-50% EtOAc/Hexanes) to afford the title compound (12.3 g, 54%).

**$^{1}$H NMR** (600 MHz, Chloroform-*d*) δ 7.67 – 7.65 (m, 4H), 7.43 – 7.35 (m, 6H), 3.77 – 3.73 (m, 4H), 3.61 – 3.58 (m, 2H), 3.54 (t, *J* = 6.4 Hz, 2H), 3.49 (t, *J* = 6.4 Hz, 2H), 3.46 (t, *J* = 6.3 Hz, 2H), 2.42 (s, 1H), 1.85 – 1.78 (m, 6H), 1.04 (d, *J* = 0.6 Hz, 9H).

**$^{13}$C NMR** (151 MHz, Chloroform-*d*) δ 135.56, 133.99, 129.54, 127.60, 70.46, 68.46, 67.76, 67.63, 62.35, 60.84, 32.72, 31.96, 30.07, 26.86, 19.24.

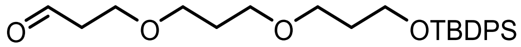

**2,2-dimethyl-3,3-diphenyl-4,8,12-trioxa-3-silapentadecan-15-al** (**S9**): To a solution of 2,2-dimethyl-3,3-diphenyl-4,8,12-trioxa-3-silapentadecan-15-ol (**S8**) (12.3 g, 28.6 mmol, 1 equiv) in anhydrous THF (500 ml) in an oven-dried flask under nitrogen at 0 °C was added Dess-Martin Periodinane (14.5 g, 34.3 mmol, 1.2 equiv) and allowed to warm to room temperature and stirred for 1.5 h. The reaction mixture was then filtered through a pad of celite. The resulting solution was concentrated *in vacuo* and subjected to silica gel chromatography (20-50% EtOAc/Hexanes) to afford the title compound (10.5 g, 86%).

**$^{1}$H NMR** (600 MHz, Chloroform-*d*) δ 9.77 (td, *J* = 2.0, 0.7 Hz, 1H), 7.69 – 7.64 (m, 4H), 7.44 – 7.34 (m, 6H), 3.74 (dt, *J* = 8.1, 6.1 Hz, 4H), 3.54 (t, *J* = 6.4 Hz, 2H), 3.49 (t, *J* = 6.4 Hz, 2H), 3.45 (t, *J* = 6.3 Hz, 2H), 2.66 – 2.61 (m, 2H), 1.81 (dp, *J* = 8.0, 6.4 Hz, 4H), 1.04 (s, 9H).

**$^{13}$C NMR** (151 MHz, Chloroform-*d*) δ 201.30, 135.56, 133.99, 129.55, 127.61, 68.25, 67.58, 64.47, 60.83, 43.87, 32.75, 29.98, 26.85, 19.24.

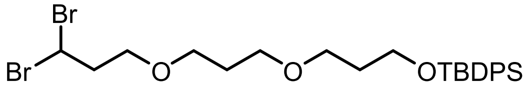

**15,15-dibromo-2,2-dimethyl-3,3-diphenyl-4,8,12-trioxa-3-silapentadecane** (**S10**): To a cold solution of triphenyl phosphite (7.1 g, 27.0 mmol, 1.1 equiv) in anhydrous dichloromethane (350 mL) maintained at -78 °C under N2 flow, bromine (1.4 ml, 27.0 mmol, 1.1 equiv) was dropped in. Anhydrous triethylamine (4.1 mL, 29.5 mmol, 1.2 equiv) and 2,2-dimethyl-3,3-diphenyl-4,8,12-trioxa-3-silapentadecan-15-al (10.5 g, 24.5 mmol, 1 equiv) were added to the faint orange solution. The reaction mixture was stirred for 4 h and allowed to warm to room temperature after 3 h. The resulting solution was concentrated *in vacuo* and subjected to silica gel chromatography (10-30% MTBE/Hexanes) to afford the title compound (10.9 g, 78%)

**$^1$H NMR** (600 MHz, Chloroform-*d*) *δ* 7.68 – 7.65 (m, 4H), 7.44 – 7.40 (m, 2H), 7.40 – 7.36 (m, 4H), 5.82 (t, *J* = 6.7 Hz, 1H), 3.75 (t, *J* = 6.1 Hz, 2H), 3.55 (t, *J* = 6.4 Hz, 2H), 3.52 – 3.44 (m, 6H), 2.61 (q, *J* = 5.9 Hz, 2H), 1.86 – 1.77 (m, 4H), 1.05 (s, 9H).

**$^{13}$C NMR** (151 MHz, Chloroform-*d*) *δ* 135.57, 133.98, 129.55, 128.35, 127.61, 77.23, 77.02, 76.81, 68.18, 68.05, 67.64, 67.58, 60.84, 45.59, 43.09, 32.76, 29.99, 27.25, 20.21.

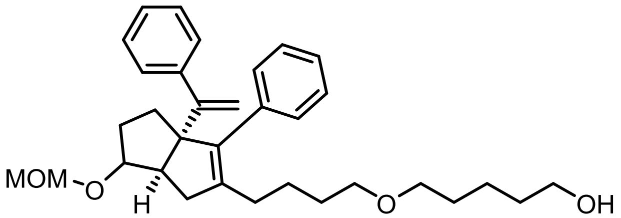

**5-(4-((3aR,6aR)-6-(methoxymethoxy)-3-phenyl-3a-(1-phenylvinyl)-1,3a,4,5,6,6a-hexahydropentalen-2-yl)butoxy)pentan-1-ol** (**S11**): According to the general cyclization procedure, **S5** (2.50 g, 4.4 mmol) was reacted with enyne (1.20 g, 4 mmol) before being treated with 5 equivalents of tetrabutylammonium fluoride to give the title compound (1.2 g, 5:1 dr, 60% yield over 2 steps) as a yellow oil after purification by flash chromatography (0-50% EtOAc/Hex eluent).

**$^1$H NMR** (600 MHz, Chloroform-*d*) δ 7.32 – 7.25 (m, 3H), 7.24 – 7.20 (m, 5H), 7.18 (dt, *J* = 8.1, 1.5 Hz, 2H), 5.02 (q, *J* = 2.5, 2.0 Hz, 1H), 4.98 (q, *J* = 2.4, 1.9 Hz, 1H), 4.60 – 4.54 (m, 2H), 3.79 – 3.73 (m, 1H), 3.66 – 3.57 (m, 4H), 3.34 (td, *J* = 6.6, 1.3 Hz, 2H), 3.28 (d, *J* = 1.3 Hz, 3H), 2.39 (dq, *J* = 9.6, 1.8 Hz, 1H), 2.26 (dddd, *J* = 9.8, 7.0, 3.1, 1.2 Hz, 1H), 2.03 – 1.96 (m, 3H), 1.60 – 1.52 (m, 8H), 1.49 – 1.34 (m, 8H).

**LRMS** (APCI) m/z[M+H]$^+$ calc'd for $C_{33}H_{44}O_4$: 505.3, found 473.3 (**13a** - CH3O$^•$)

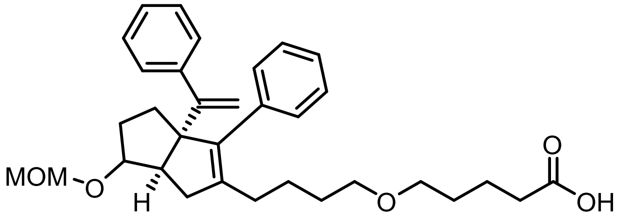

**5-(4-((3aR,6aR)-6-(methoxymethoxy)-3-phenyl-3a-(1-phenylvinyl)-1,3a,4,5,6,6a-hexahydropentalen-2-yl)butoxy)pentanoic acid** (**S12**): A solution of **S11** (504 mg, 1 eq) in 50 mL acetonitrile was treated with tetrapropylammonium perruthenate (TPAP, 0.1 equiv, 17.55 mg), N-Methylmorpholine-N-Oxide (NMO, 10 equiv, 1.17 g), and water (10 equiv, 180 µL). The reaction mixture was allowed to stir overnight at room temperature. The reaction mixture was passed through celite, concentrated and purified on silica (0-65% EtOAc (containing 0.1% AcOH)/hexanes eluent) to provide the title compound. 70 mg, 13%

**$^1$H NMR** (600 MHz, Chloroform-*d*) δ 7.38 – 7.28 (m, 4H), 7.28 – 7.25 (m, 3H), 7.25 – 7.22 (m, 2H), 5.06 (d, *J* = 1.2 Hz, 1H), 5.04 (t, *J* = 1.2 Hz, 1H), 4.64 (s, 2H), 3.81 (dd, *J* = 4.5, 2.6 Hz, 1H), 3.42 (q, *J* = 5.8 Hz, 2H), 3.37 (dd, *J* = 6.1, 1.2 Hz, 5H), 2.39 (t, *J* = 7.6 Hz, 2H), 1.71 – 1.59 (m, 4H), 1.56 – 1.41 (m, 5H), 1.36 – 1.23 (m, 10H).

**$^{13}$C NMR** (151 MHz, $CDCl_3$) δ 154.47, 144.05, 137.50, 129.58, 127.85, 127.71, 127.66, 126.65, 114.90, 104.61, 94.51, 86.79, 70.37, 70.21, 69.71, 55.06, 52.80, 40.46, 33.66, 33.58, 32.37, 31.58, 29.67, 29.43, 28.98, 28.62, 24.49, 21.80, 21.58.

**LRMS** (APCI) m/z[M+H]$^+$ calc'd for $C_{33}H_{42}O_5$: 519.3, found 487.3 (**S12** - CH3O•)

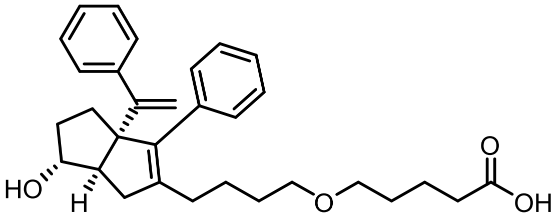

**5-(4-((3aR,6aR)-6-hydroxy-3-phenyl-3a-(1-phenylvinyl)-1,3a,4,5,6,6a-hexahydropentalen-2-yl)butoxy)pentanoic acid** (**3**): 70 mg of **S12** was dissolved in 200 mL DCM and then 5 drops of TFA was added. The reaction was stirred at room temp until complete by TLC and HPLC. The reaction was then concentrated and purified by silica chromatography (30-100% EtOAc/Hexanes w/ 1% AcOH). 27 mg, 42%

**$^1$H NMR:** (600 MHz, Chloroform-*d*) δ 7.35 – 7.28 (m, 5H), 7.24 – 7.18 (m, 5H), 5.06 (d, *J* = 1.3 Hz, 1H), 5.00 (d, *J* = 1.4 Hz, 1H), 3.95 (s, 1H), 3.39 (t, *J* = 6.2 Hz, 2H), 3.34 (t, *J* = 6.3 Hz, 2H), 2.39 (t, *J* = 7.3 Hz, 2H), 2.34 – 2.28 (m, 1H), 2.12 – 1.98 (m, 4H), 1.77 – 1.57 (m, 11H), 1.52 – 1.45 (m, 1H), 1.42 (q, *J* = 7.6, 7.2 Hz, 1H).

**$^{13}$C NMR** (151 MHz, $CDCl_3$) δ 154.57, 144.10, 140.82, 139.51, 137.32, 129.68, 127.80, 127.74, 127.70, 126.69, 126.68, 114.99, 82.03, 70.54, 70.28, 69.30, 55.79, 40.09, 33.99, 33.71, 32.06, 29.72, 29.65, 29.44, 28.99, 24.47, 21.67.

**LRMS** (APCI) m/z[M+H]$^+$ calc'd for $C_{31}H_{38}O_4$: 475.3, found 475.3

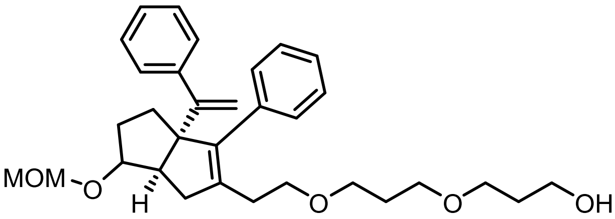

**3-(3-(2-((3aR,6aR)-6-(methoxymethoxy)-3-phenyl-3a-(1-phenylvinyl)-1,3a,4,5,6,6a-hexahydropentalen-2-yl)ethoxy)propoxy)propan-1-ol** (**S14**): According to the general cyclization procedure, **S10** (2.50 g, 4.4 mmol) was reacted with enyne (1.20 g, 4 mmol) before being treated with 5 equivalents of tetrabutylammonium fluoride to give the title compound (1.24 g, 7:1 dr, 61% yield over 2 steps) as a yellow oil after purification by flash chromatography (0-50% EtOAc/Hex eluent).

**$^1$H NMR**: (600 MHz, Chloroform-*d*) δ 7.32 (ddq, *J* = 6.0, 2.3, 1.3 Hz, 2H), 7.28 – 7.25 (m, 2H), 7.26 – 7.19 (m, 6H), 5.03 (q, *J* = 1.3 Hz, 1H), 4.98 (dd, *J* = 2.0, 1.3 Hz, 1H), 4.58 – 4.53 (m, 2H), 3.76 – 3.75 (m, 1H), 3.58 – 3.54 (m, 2H), 3.48 – 3.44 (m, 2H), 3.42 – 3.35 (m, 4H), 3.27 (d, *J* = 2.0 Hz, 3H), 2.40

(ddt, *J* = 11.0, 7.1, 1.7 Hz, 2H), 2.36 – 2.25 (m, 3H), 2.07 (d, *J* = 16.9 Hz, 1H), 1.78 (dddt, *J* = 7.8, 5.9, 3.4, 1.8 Hz, 5H), 1.67 – 1.58 (m, 2H).

**LRMS** (APCI) m/z$[M+H]^+$ calc'd for $C_{32}H_{42}O_5$: 507.3, found 475.3 (**S14** - CH3O$^{\bullet}$)

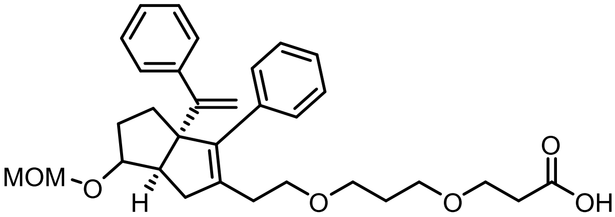

**3-(3-(2-((3aR,6aR)-6-(methoxymethoxy)-3-phenyl-3a-(1-phenylvinyl)-1,3a,4,5,6,6a-hexahydropentalen-2-yl)ethoxy)propoxy)propanoic acid** (**S15**): A solution of **S15** (504 mg, 1 eq) in 50 mL acetonitrile was treated with tetrapropylammonium perruthenate (TPAP, 0.1 equiv, 17.55 mg), N-Methylmorpholine-N-Oxide (NMO, 10 equiv, 1.17 g), and water (10 equiv, 180 µL). The reaction mixture was allowed to stir overnight at room temperature. The reaction mixture was passed through celite, concentrated and purified on silica (20-80% EtOAc (containing 0.1% AcOH)/hexanes eluent) to provide the title compound. (285 mg, 45%).

**$^1$H NMR** (600 MHz, Chloroform-*d*) δ 7.35 – 7.30 (m, 2H), 7.29 – 7.25 (m, 2H), 7.24 – 7.20 (m, 6H), 5.03 (d, *J* = 1.4 Hz, 1H), 4.98 (d, *J* = 1.4 Hz, 1H), 4.61 – 4.55 (m, 2H), 3.80 – 3.74 (m, 1H), 3.67 – 3.63 (m, 2H), 3.52 – 3.46 (m, 2H), 3.42 – 3.34 (m, 5H), 3.29 (t, *J* = 1.0 Hz, 2H), 2.57 (td, *J* = 6.2, 1.4 Hz, 2H), 2.39 (dt, *J* = 9.3, 1.6 Hz, 1H), 2.35 – 2.29 (m, 2H), 2.26 (dt, *J* = 13.8, 6.5 Hz, 1H), 2.11 – 2.08 (m, 1H), 2.08 – 2.04 (m, 2H), 2.01 – 1.96 (m, 1H), 1.78 (pd, *J* = 6.4, 1.6 Hz, 2H), 1.74 – 1.67 (m, 1H), 1.67 – 1.58 (m, 2H).

**LRMS** (APCI) m/z calc'd for $C_{32}H_{40}O_6$: 521.3, found 489.3 (**S15** - CH3O$^{\bullet}$)

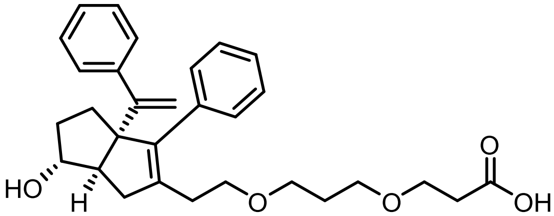

**3-(3-(2-((3aR,6aR)-6-hydroxy-3-phenyl-3a-(1-phenylvinyl)-1,3a,4,5,6,6a-hexahydropentalen-2-yl)ethoxy)propoxy)propanoic acid** (**5**): 125 mg of **S15** was dissolved in 10 mL acetonitrile and 5 drops of 12M HCl. Reaction stirred for 10 minutes, then concentrated and purified by prep HPLC (15-75% ACN in water, 0.1% formic acid modifier, 30 minute gradient). 97 mg, 85%
**$^1$H NMR** (600 MHz, Chloroform-*d*) δ 7.37 (dd, *J* = 6.6, 2.9 Hz, 2H), 7.32 (t, *J* = 7.3 Hz, 2H), 7.28 (d, *J* = 7.8 Hz, 6H), 5.09 (s, 1H), 5.03 (s, 1H), 4.01 – 3.93 (m, 1H), 3.70 (t, *J* = 6.2 Hz, 2H), 3.54 (t, *J* = 6.3 Hz, 2H), 3.44 (qd, *J* = 9.2, 4.6 Hz, 4H), 2.61 (t, *J* = 6.1 Hz, 2H), 2.42 – 2.33 (m, 2H), 2.31 (dd, *J* = 12.6, 6.9 Hz, 2H), 2.14 (d, *J* = 17.2 Hz, 1H), 1.83 (p, *J* = 6.3 Hz, 2H), 1.74 (tt, *J* = 10.9, 5.5 Hz, 1H), 1.49 (pd, *J* = 7.1, 3.4 Hz, 1H), 1.43 (q, *J* = 7.7, 6.6 Hz, 1H), 1.36 (p, *J* = 3.5 Hz, 1H).

**LRMS** (APCI) m/z$[M+H]^+$ calc'd for $C_{30}H_{36}O_5$: 477.3, found 477.3

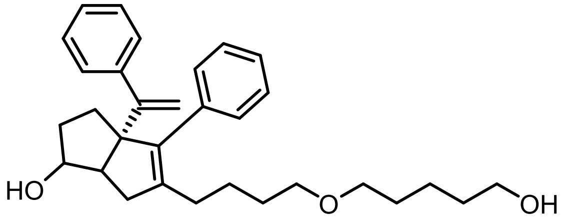

**(3a*R*)-5-(4-((5-hydroxypentyl)oxy)butyl)-4-phenyl-3a-(1-phenylvinyl)-1,2,3,3a,6,6a-hexahydropentalen-1-ol** (**S16**): According to the general cyclization procedure, **S5** (2.50 g, 4.4 mmol) was reacted with enyne (1.20 g, 4 mmol) before being treated with three drops of HCl to give the title compound (1.21 g, 66% yield over 2 steps) as a yellow oil after purification by flash chromatography (0-65% EtOAc/Hex eluent).

**$^1$H NMR** (600 MHz, Chloroform-*d*) δ 7.35 – 7.30 (m, 5H), 7.25 (m, 2H), 7.20 (t, *J* = 6.8 Hz, 3H), 5.07 (d, *J* = 1.6 Hz, 1H), 4.99 (dp, *J* = 1.5 Hz, 1H), 3.95 (m, 1H), 3.65 (t, *J* = 6.6 Hz, 3H), 3.38 (t, *J* = 6.4 Hz, 4H), 3.33 (t, *J* = 6.3 Hz, 2H), 2.64 (d, *J* = 17.3 Hz, 1H), 2.47 (t, *J* = 8.7 Hz, 1H), 2.42 – 2.24 (m, 2H), 2.14 – 1.94 (m, 6H), 1.76 – 1.64 (m, 4H), 1.59 (dd, *J* = 10.7, 4.6 Hz, 4H), 1.55 – 1.46 (m, 1H).

**LRMS** (APCI) m/z[M+H]$^+$ calc'd for $C_{31}H_{41}O_3$: 461.7, found 461.3

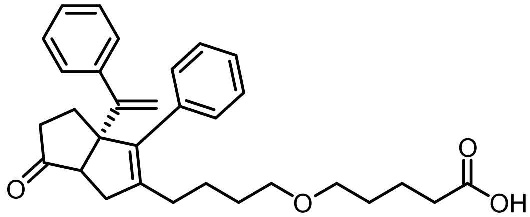

**5-(4-((3a*R*)-6-oxo-3-phenyl-3a-(1-phenylvinyl)-1,3a,4,5,6,6a-hexahydropentalen-2-yl)butoxy)pentanoic acid** (**S17**): According to the oxidation general procedure, **S16** (1.21 g, 2.63 mmol) was reacted with TPAP (92.4 mg, 0.26 mmol), NMO (3.07 g, 26 mmol), water (473 µL, 473 mg, 26 mmol), and acetonitrile (40 mL) to give the title compound (529 mg, 43% yield) as an oil after purification by flash chromatography (0-65% EtOAc/Hex).

**$^1$H NMR** (600 MHz, $CDCl_3$) δ 7.40 – 7.31 (m, 6H), 7.24 (m, 4H), 5.24 (dt, *J* = 3.8, 1.4 Hz, 1H), 5.13 (q, *J* = 1.6 Hz, 1H), 3.39 (tdd, *J* = 6.2, 3.6, 1.4 Hz, 2H), 3.32 (qd, *J* = 5.1, 1.9 Hz, 2H), 2.52 – 2.48 (m, 1H), 2.40 (tdd, *J* = 7.5, 3.7, 1.5 Hz, 2H), 2.37 – 2.29 (m, 2H), 2.15 – 2.08 (m, 4H), 2.07 – 1.99 (m, 2H), 1.93 (ddd, *J* = 16.5, 8.3, 2.9 Hz, 1H), 1.75 – 1.69 (m, 2H), 1.66 – 1.58 (m, 2H), 1.49 – 1.40 (m, 2H), 1.35 (qd, *J* = 7.9, 4.3 Hz, 2H).

**$^{13}$C NMR** (151 MHz, $CDCl_3$) δ 177.36, 153.15, 144.47, 142.49, 137.82, 136.55, 128.98, 128.28, 128.17, 127.65, 127.15, 127.02, 115.36, 70.51, 70.31, 65.48, 55.56, 38.64, 37.52, 33.47, 29.70, 28.91, 28.37, 24.32, 21.69.

**LRMS** (APCI) m/z[M+H]$^+$ calc'd for $C_{31}H_{37}O_4$: 473.6, found 473.3

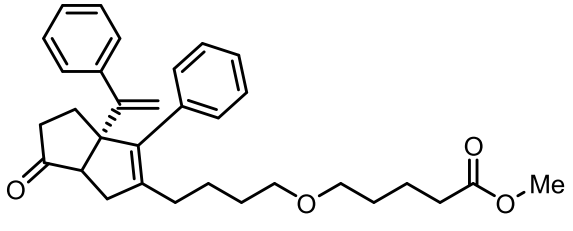

**Methyl 5-(4-((3a*R*)-6-oxo-3-phenyl-3a-(1-phenylvinyl)-1,3a,4,5,6,6a-hexahydropentalen-2-yl)butoxy)pentanoate** (**S18**): According to the methylation general procedure, **S17** (529 mg, 1.1 mmol) was reacted with HCl in methanol to give the title compound (545 mg, quantitative yield) as a yellow oil.

**$^1$H NMR** (600 MHz, $CDCl_3$) δ 7.40 – 7.36 (m, 2H), 7.33 (tt, *J* = 8.0, 1.4 Hz, 3H), 7.30 – 7.26 (m, 3H), 7.25 – 7.22 (m, 2H), 5.24 (d, *J* = 1.3 Hz, 1H), 5.13 (d, *J* = 1.3 Hz, 1H), 3.68 (s, 3H), 3.37 (t, *J* = 6.4 Hz, 2H), 3.32 – 3.29 (m, 2H), 2.49 (d, *J* = 7.8 Hz, 1H), 2.35 (t, *J* = 7.5 Hz, 3H), 2.33 – 2.27 (m, 1H), 2.17 – 2.08 (m, 4H), 2.05 (dd, *J* = 13.5, 6.4 Hz, 2H), 1.94 (dd, *J* = 16.5, 7.8 Hz, 1H), 1.73 – 1.66 (m, 2H), 1.62 – 1.55 (m, 3H), 1.44 (ddd, *J* = 13.1, 8.2, 5.8 Hz, 2H), 1.38 – 1.30 (m, 2H).

**$^{13}$C NMR** (151 MHz, $CDCl_3$) δ 174.05, 153.19, 144.47, 142.51, 137.80, 136.57, 128.99, 128.28, 128.16, 127.65, 127.14, 127.02, 115.35, 70.47, 70.32, 65.47, 55.52, 51.48, 38.71, 37.50, 33.83, 29.76, 29.73, 29.14, 28.39, 24.33, 21.75.

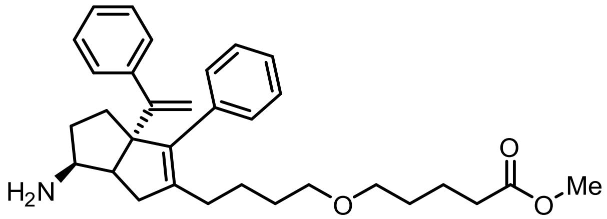

**Methyl 5-(4-((3a*R*,6*S*)-6-amino-3-phenyl-3a-(1-phenylvinyl)-1,3a,4,5,6,6a-hexahydropentalen-2-yl)butoxy)pentanoate** (**S19**): According to the reductive amination general procedure **S18** (545 mg, 1.13 mmol) was reacted with Ammonia (816 µL, 5.65 mmol), Ti(O*i*Pr)$_4$ (517 µL, 1.7 mmol), and $NaBH_4$ (128 mg, 3.4 mmol) in ethanol (11 mL) to give the title compound (235 mg, 42% yield) as a clear oil after purification by flash chromatography (0-10% MeOH/DCM).

**$^1$H NMR** (600 MHz, $CDCl_3$) δ 7.35 – 7.32 (m, 2H), 7.30 (d, *J* = 7.5 Hz, 2H), 7.27 (dd, *J* = 5.0, 2.0 Hz, 4H), 7.22 – 7.20 (m, 2H), 5.08 (d, *J* = 1.4 Hz, 1H), 4.95 (d, *J* = 1.5 Hz, 1H), 3.68 (s, 3H), 3.39 (t, *J* = 6.4 Hz, 2H), 3.35 (td, *J* = 6.3, 1.6 Hz, 2H), 3.33 – 3.29 (m, 1H), 2.50 (dd, *J* = 17.4, 2.2 Hz, 1H), 2.44 (td, *J* = 8.9, 2.2 Hz, 1H), 2.35 (t, *J* = 7.5 Hz, 2H), 2.16 – 2.00 (m, 4H), 1.85 – 1.79 (m, 1H), 1.70 (qd, *J* = 7.1, 5.0 Hz, 4H), 1.60 (dq, *J* = 9.8, 6.5 Hz, 2H), 1.55 – 1.45 (m, 6H).

**$^{13}$C NMR** (151 MHz, $CDCl_3$) δ 174.07, 155.05, 144.20, 142.49, 139.84, 137.08, 129.78, 127.75, 127.69, 127.62, 126.65, 126.58, 115.08, 70.56, 70.32, 69.52, 55.31, 51.49, 49.07, 34.48, 34.02, 33.83, 33.22, 29.81, 29.61, 29.15, 24.58, 21.78.

**LRMS** (APCI) m/z$[M+H]^+$ calc'd for $C_{32}H_{42}NO_3$: 488.7, found 488.3

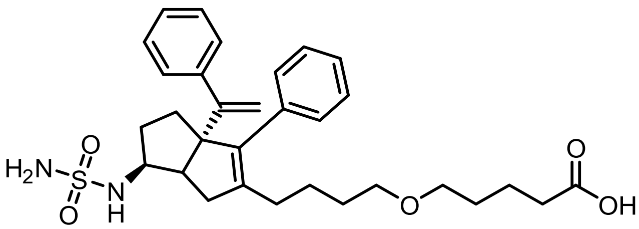

**5-(4-((3a*R*,6*S*)-3-phenyl-3a-(1-phenylvinyl)-6-(sulfamoylamino)-1,3a,4,5,6,6a-hexahydropentalen-2-yl)butoxy)pentanoic acid** (4, **ME610**): **S19** was added to a reaction vial in EtOH (0.1M). $H_2O$ (0.05 M) was added followed by sulfamide (5 equiv) and triethylamine (3 equiv). The reaction proceeded for 16 hours at reflux before being cooled to room temperature. The reaction mixture was concentrated via rotatory evaporation. The crude mixture was then purified via flash chromatography (10-20% MeOH/DCM) to give the title compound as a yellow oil.

**$^1$H NMR** (600 MHz, $CDCl_3$) δ 7.35 – 7.31 (m, 3H), 7.32 – 7.29 (m, 2H), 7.28 (d, *J* = 5.8 Hz, 3H), 7.20 – 7.17 (m, 2H), 5.71 – 5.66 (m, 1H), 5.13 (d, *J* = 1.2 Hz, 1H), 4.92 (d, *J* = 1.2 Hz, 1H), 4.76 – 4.66 (br. s, 2H), 3.81 (m, 1H), 3.52 – 3.40 (m, 4H), 2.66 (td, *J* = 9.1, 2.4 Hz, 1H), 2.49 (dd, *J* = 17.7, 2.4 Hz, 1H), 2.42 (td, *J* = 7.5, 1.3 Hz, 2H), 2.21 – 2.14 (m, 5H), 2.00 (dt, *J* = 9.8, 5.3 Hz, 2H), 1.84 (dq, *J* = 14.7, 7.3 Hz, 1H), 1.80 – 1.69 (m, 3H), 1.68 – 1.62 (m, 2H), 1.60 – 1.50 (m, 3H).

**$^{13}$C NMR** (151 MHz, $CDCl_3$) δ 177.19, 154.33, 143.88, 142.75, 139.44, 136.68, 129.85, 127.90, 127.69, 127.58, 126.88, 126.78, 115.76, 70.03, 69.58, 68.72, 57.06, 47.05, 35.37, 33.03, 32.77, 31.28, 30.94, 29.81, 29.71, 28.51, 24.93, 21.71.

**LRMS** (APCI) m/z[M+H]$^+$ calc'd for $C_{31}H_{41}N_2O_5S$: 553.7, found 553.3

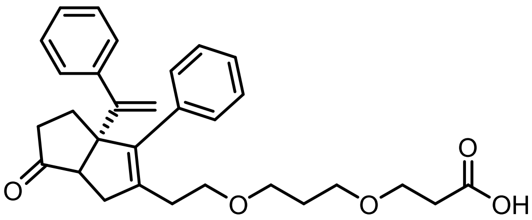

**3-(3-(2-((3a*R*)-6-oxo-3-phenyl-3a-(1-phenylvinyl)-1,3a,4,5,6,6a-hexahydropentalen-2-yl)ethoxy)propoxy)propanoic acid** (**S20**): According to the methylation general procedure, **S19** (403.4 mg, 0.85mmol) was reacted with HCl in methanol to give the title compound (299 mg, 70% yield) as a yellow oil after purification by flash chromatography (30% EtOAc/Hex).

**$^1$H NMR** (600 MHz, $CDCl_3$) δ 7.38 (d, *J* = 4.4 Hz, 4H), 7.36 – 7.31 (m, 1H), 7.31 – 7.28 (m, 3H), 7.25 (dd, *J* = 7.5, 1.9 Hz, 2H), 5.27 (s, 1H), 5.15 (s, 1H), 3.72 (t, *J* = 6.1 Hz, 2H), 3.52 (q, *J* = 5.9 Hz, 2H), 3.45 – 3.32 (m, 4H), 2.62 (t, *J* = 6.1 Hz, 2H), 2.50 (d, *J* = 7.8 Hz, 1H), 2.43 (dd, *J* = 14.2, 7.1 Hz, 1H), 2.39 (d, *J* = 16.6 Hz, 1H), 2.29 (ddt, *J* = 20.5, 13.4, 6.9 Hz, 2H), 2.19 – 2.08 (m, 2H), 2.08 – 2.01 (m, 1H), 1.97 (dd, *J* = 16.6, 7.8 Hz, 1H), 1.85 – 1.75 (m, 2H).

**$^{13}$C NMR** (151 MHz, $CDCl_3$) δ 174.03, 152.96, 142.48, 141.53, 139.42, 136.31, 129.07, 128.26, 128.17, 127.67, 127.25, 127.05, 115.51, 68.76, 68.21, 67.57, 65.46, 55.67, 38.89, 37.55, 34.70, 30.31, 29.92, 28.37.

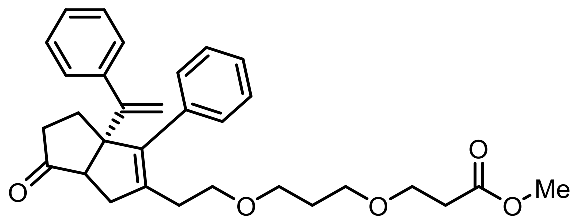

**Methyl 3-(3-(2-((3a*R*)-6-oxo-3-phenyl-3a-(1-phenylvinyl)-1,3a,4,5,6,6a-hexahydropentalen-2-yl)ethoxy)propoxy)propanoate** (**S21**): According to the methylation general procedure, **S20** (403.4 mg, 0.85 mmol) was reacted with HCl in methanol to give the title compound (299 mg, 70% yield) as a yellow oil after purification by flash chromatography (30% EtOAc/Hex).

**$^{1}$H NMR** (600 MHz, $CDCl_3$) δ 7.38 (d, *J* = 4.3 Hz, 4H), 7.34 (h, *J* = 4.4 Hz, 1H), 7.31 – 7.28 (m, 3H), 7.25 (dd, *J* = 7.6, 2.0 Hz, 2H), 5.27 (s, 1H), 5.15 (s, 1H), 3.70 (m, 4H), 3.48 (tt, *J* = 6.5, 3.2 Hz, 2H), 3.48 (dtt, *J* = 7.5, 3.1, 1.4 Hz) 1H), 3.37 (dddd, *J* = 31.9, 16.1, 9.2, 6.7 Hz, 4H), 2.58 (t, *J* = 6.4 Hz, 2H), 2.48 (d, *J* = 7.8 Hz, 1H), 2.44 – 2.35 (m, 2H), 2.30 (dq, *J* = 13.6, 6.8 Hz, 2H), 2.17 – 2.01 (m, 3H), 1.98 (dd, *J* = 16.5, 7.9 Hz, 1H), 1.78 (p, *J* = 6.4 Hz, 2H).

**$^{13}$C NMR** (151 MHz, $CDCl_3$) δ 172.10, 152.99, 142.52, 141.34, 139.50, 136.32, 129.07, 128.26, 128.15, 127.66, 127.23, 127.03, 115.50, 68.56, 68.01, 67.69, 66.12, 65.46, 55.59, 51.65, 50.92, 38.86, 37.51, 34.93, 30.40, 29.97, 28.40.

**LRMS** (APCI) m/z$[M+H]^+$ calc'd for $C_{31}H_{37}O_5$: 489.6, found 489.3

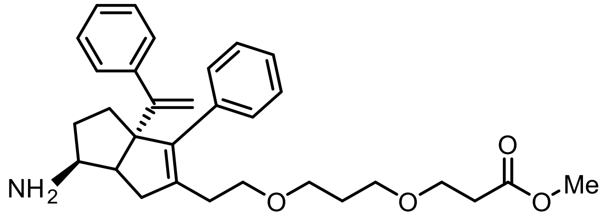

**Methyl 3-(3-(2-((3aR,6S)-6-amino-3-phenyl-3a-(1-phenylvinyl)-1,3a,4,5,6,6a-hexahydropentalen-2-yl)ethoxy)propoxy)propanoate (S22):** According to the reductive amination general procedure **S21** (299.1 mg, 0.6 mmol) was reacted with Ammonia (1.7 mL, 12.2 mmol), $Ti(O\textit{i}Pr)_4$ (272 µL, 261.4 mg, 0.92 mmol), and $NaBH_4$ (68.1 mg, 1.8 mmol) in ethanol (6 mL) to give the title compound (210.8 mg, 67% yield) as an orange oil after purification by flash chromatography (0-10% Methanol/DCM).

**$^{1}$H NMR** (600 MHz, Chloroform-d) δ 7.53 – 7.36 (m, 2H), 7.31 (dd, J = 8.3, 6.5 Hz, 3H), 7.29 – 7.25 (m, 3H), 7.19 (dq, J = 5.3, 2.9, 2.5 Hz, 2H), 5.18 (d, J = 1.5 Hz, 1H), 5.07 (d, J = 1.6 Hz, 1H), 3.93 – 3.86 (m, 1H), 3.75 – 3.66 (m, 3H), 3.65 (s, 3H), 3.60 – 3.51 (m, 3H), 3.48 (ddd, J = 9.2, 5.6, 3.2 Hz, 1H), 2.89 – 2.75 (m, 1H), 2.70 (dd, J = 9.7, 6.7 Hz, 1H), 2.66 – 2.58 (m, 2H), 2.54 (ddd, J = 16.2, 6.8, 4.4 Hz, 2H), 2.24 – 2.16 (m, 1H), 2.00 – 1.94 (m, 3H), 1.93 – 1.79 (m, 2H), 1.73 (dq, J = 15.9, 6.0 Hz, 2H), 1.49 – 1.38 (m, 1H).

**$^{13}$C NMR** (151 MHz, $CDCl_3$) δ 172.18, 143.78, 141.27, 139.69, 136.61, 129.70, 127.84, 127.70, 126.76, 115.24, 69.44, 68.83, 67.98, 67.66, 66.10, 51.68, 50.87, 34.91, 34.18, 32.69, 29.98, 29.93.

**LRMS** (APCI) m/z$[M+H]^+$ calc'd for $C_{31}H_{40}NO_4$: 490.6, found 490.3

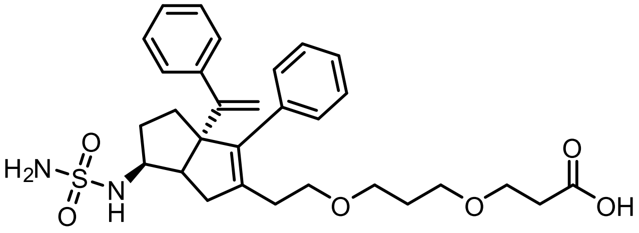

**3-(3-(2-((3aR,6S)-3-phenyl-3a-(1-phenylvinyl)-6-(sulfamoylamino)-1,3a,4,5,6,6a-hexahydropentalen-2-yl)ethoxy)propoxy)propanoic acid (6): S22** (26.3 mg, 0.05 mmol, 1.0 equiv) was added to a reaction vial in EtOH (500 µL). H2O (1.0 mL) was added followed by sulfamide (24 mg, 0.25 mmol, 5 equiv) and triethylamine (21 µL, 0.15 mmol, 3 equiv). The reaction proceeded for 16 hours at reflux before being cooled to room temperature. The reaction mixture was concentrated via rotatory evaporation. The crude mixture was then purified via flash chromatography (10-20% MeOH/DCM) to give the title compound (19.9 mg, 70% yield) as a yellow oil.

**$^1$H NMR** (600 MHz, $CDCl_3$) δ 7.41 – 7.39 (m, 2H), 7.33 (t, J = 7.2 Hz, 2H), 7.28 (t, J = 1.0 Hz, 8H), 5.45 (d, J = 9.0 Hz, 1H), 5.11 (d, J = 1.4 Hz, 1H), 5.09 (d, J = 1.6 Hz, 1H), 4.76 (br. s, 2H), 3.87 (t, J = 8.7 Hz, 1H), 3.77 – 3.72 (m, 2H), 3.69 (td, J = 9.6, 5.3 Hz, 1H), 3.64 (dt, J = 9.7, 6.3 Hz, 2H), 3.58 (dt, J = 10.5, 6.3 Hz, 1H), 3.53 (td, J = 9.6, 4.9 Hz, 2H), 2.63 (dddd, J = 30.5, 17.3, 9.8, 6.6 Hz, 4H), 2.53 (d, J = 17.2 Hz, 1H), 2.05 (tt, J = 12.4, 4.9 Hz, 2H), 1.96 – 1.84 (m, 3H), 1.42 – 1.36 (m, 2H).

**$^{13}$C NMR** (151 MHz, $CDCl_3$) δ 175.22, 175.12, 153.75, 143.31, 140.71, 140.25, 136.41, 129.47, 128.02, 127.81, 127.68, 126.87, 126.83, 115.11, 68.76, 68.69, 67.69, 67.57, 65.84, 56.69, 34.78, 34.60, 32.69, 31.45, 29.78, 29.73, 20.40.

**LRMS** (APCI) m/z[M+H]$^+$ calc'd for $C_{30}H_{39}N_2O_6S$: 555.7, found 555.3

**Solubility and Metabolic Stability**

Studies were conducted by Pharmaron, Inc, using protocols detailed below.

*Solubility*

Stock solutions of test compounds were prepared in DMSO at 50 mM and then diluted with DMSO to 30 mM. Stock solution of control compound (progesterone) was prepared in DMSO at 30 mM. 10 µL stock solution of test compounds were placed in 96-well rack, followed by 990 µL PBS (pH 7.4) into each vial of the solubility sample plate. One stir stick was added to each vial and then vials were sealed using a molded PTDE/SIL 96-well plate cover. The solubility sample plate was transferred to the thermomixer comfort plate shaker and incubated at room temperature for two hours (1100 rpm). All samples from the solubility sample plate were transferred into the filter plate and filtered by using the vacuum manifold. Filtered samples were diluted with methanol (dilution factor may be changed according to the solubility value and the LC/MS signal response). Compound stock solutions were diluted with DMSO to 300 µM and then diluted with methanol to obtain 300 nM STDs. The solution filtered was analyzed and quantified against a standard of known concentration in DMSO using liquid chromatography coupled with mass spectral peak identification and quantitation. This study was performed in duplicate. Solubility values of the test compounds were calculated as follows:

$$[Sample] = \frac{AREA_{Sample} \times INJVOL_{Std} \times DF_{Sample} \times [STD]}{AREA_{Std} \times INJVOL_{Sample}}$$

*Metabolic stability*

The master solution was prepared according to following:

| Reagent | [Stock] | Volume (µL) | [Final] |
|---|---|---|---|
| $PO_4$ buffer | 200 mM | 200 | 100 mM |
| Ultra-pure $H_2O$ | - | 66 | - |
| $MgCl_2$ solution | 50 mM | 40 | 5 mM |
| Alamethacin | 5 mg/mL | 2 | 25 µg/mL |
| Mouse liver microsomes, pooled | 20 mg/mL | 10 | 0.5 mg/mL |

40 µL of 10 mM NADPH solution and 40 µL of 20 mM UDPGA solution were added to each well. Final concentrations of NADPH and UDPGA were 1 mM and 2 mM, respectively. The mixture was pre-warmed at 37 °C for 5 minutes. The negative control samples were prepared by replacing cofactors (NADPH and UDPGA) solution with 80 µL of ultra-pure H2O. Negative control was used to exclude the misleading factor that resulted from instability of chemical itself. Samples with cofactors were prepared in duplicate. Negative controls were prepared in singlet. The reaction was started with the addition of 2 µL of 400 µM control compound or test compound solutions. Diclofenac was used as a positive control in this study. The final concentration of test compound or control compound was 2 µM. Aliquots of 50 µL were taken from the reaction solution at 0, 15, 30, 45 and 60 minutes. The reaction was stopped by the addition of 4 volumes of cold acetonitrile with internal standards (100 nM alprazolam, 200 nM imipramine, 200 nM labetalol and 2 µM ketoprofen). Samples were centrifuged at 3,220 g for 40 minutes. 90 µL of the supernatant were mixed with 90 µL of ultra-pure $H_2O$ and then used for LC-MS/MS analysis. Peak areas were determined from extracted ion chromatograms.

**Pharmacokinetics**

Conducted by Pharmaron, Inc, as follows:

Compounds were dissolved in DMSO and diluted in water before being given to mice at a single dose of 1 mg/kg compound (intravenous cohort) or 20 mg/kg (IP cohorts); Vehicle was 5% DMSO, and 95% water, pH adjusted to ~8.0 with 0.1 N NaOH. To quantify exposure, analyte stock solutions were serially diluted in 50% acetonitrile in water solution. 5 µL of working solutions (1, 2, 4, 10, 20, 100, 200, 1000, 2000 ng/mL) were added to 10 µL of the blank male C57BL/6J Mouse plasma to achieve calibration standards of 0.5~1000 ng/mL (0.5, 1, 2, 5, 10, 50, 100, 500, 1000 ng/mL) in a total volume of 15 µL. Four quality control samples at 1 ng/mL, 2 ng/mL, 50 ng/mL and 800 ng/mL for plasma were prepared independently of those used for the calibration curves. These QC samples were prepared on the day of analysis in the same way as calibration standards. 15 µL standards, 15 µL QC samples and 15 µL unknown samples （10 µL plasma with 5 µL blank solution) were added to 150 µL of acetonitrile containing IS mixture for precipitating protein respectively. Then the samples were vortexed for 30 s. After centrifugation at 4°C, 4000 rpm for 15 min, the supernatant was diluted 3 times with water. 20 µL of diluted supernatant was injected into the LC/MS/MS system for quantitative analysis. Bioavailability (F%) was calculated as following: $AUC_{last}$-PO/$AUC_{INF}$-PO > 80%: F=($AUC_{INF}$-PO*$Dose_{IV}$)/(mean $AUC_{INF}$-IV*DosePO); $AUC_{last}$-PO/$AUC_{INF}$-PO ≤ 80% or $AUC_{INF}$ was not available: F=($AUC_{last}$-PO*DoseIV)/(mean $AUC_{last}$-IV*DosePO)

Supplementary tables

**Table S1. Summary of pharmacokinetic parameters**

**Table S1a. IV administration, ME610**

**IV Dose 1 mg/kg**

| PK parameters | Unit | Mean | SD | CV(%) |
|---|---|---|---|---|
| Cl_obs | mL/min/kg | 408 | 86 | 21.1 |
| $T_{1/2}$ | h | 1.00 | 0.10 | 10.5 |
| $C_0$ | ng/mL | 174 | 78 | 44.9 |
| $AUC_{last}$ | h*ng/mL | 39.4 | 8.7 | 22.0 |
| $AUC_{Inf}$ | h*ng/mL | 42.1 | 9.0 | 21.5 |
| $AUC_{_\%Extrap_}$obs | % | 6.31 | 2.57 | 40.7 |
| $MRT_{Inf_}$obs | h | 0.85 | 0.24 | 28.2 |
| $AUC_{last}$/D | h*ng/mL | 39.4 | 8.7 | 22.0 |
| $V_{ss}$_obs | L/kg | 20.8 | 6.9 | 33.1 |

**Table S1b. IP administration, ME610**

**IP Dose 20 mg/kg**

| PK parameters | Unit | Mean | SD | CV(%) |
|---|---|---|---|---|
| $T_{1/2}$ | h | 1.28 | 0.19 | 15.0 |
| $T_{max}$ | h | 0.333 | 0.144 | 43.3 |
| $C_{max}$ | ng/mL | 2960 | 348 | 11.8 |
| $AUC_{last}$ | h*ng/mL | 3760 | 306 | 8.15 |
| $AUC_{Inf}$ | h*ng/mL | 3791 | 324 | 8.55 |
| AUC_%Extrap_obs | % | 0.78 | 0.41 | 52.0 |
| $MRT_{Inf_}$obs | h | 1.21 | 0.21 | 17.6 |
| $AUC_{last}$/D | h*ng/mL | 188 | 15 | 8.15 |

**Table S1c. Oral administration, ME610**

**PO Dose 20 mg/kg**

| PK parameters | Unit | Mean | SD | CV(%) |
|---|---|---|---|---|
| $T_{1/2}$ | h | 3.88 | 5.09 | 131 |
| $T_{max}$ | h | 0.250 | 0.000 | 0.000 |
| $C_{max}$ | ng/mL | 203 | 28 | 13.6 |
| $AUC_{last}$ | h*ng/mL | 224 | 25 | 11.1 |
| $AUC_{Inf}$ | h*ng/mL | 243 | 47 | 19.4 |
| AUC_%Extrap_obs | % | 6.5 | 9.7 | 150 |
| $MRT_{Inf_}$obs | h | 2.58 | 2.03 | 78.6 |
| $AUC_{last}$/D | h*ng/mL | 11.2 | 1.2 | 11.1 |
| F | % | 28.8 | 5.6 | 19.4 |

**Table S1d. IV administration, compound 6**

**IV Dose 1 mg/kg**

| PK parameters | Unit | Mean | SD | CV(%) |
|---|---|---|---|---|
| Cl_obs | mL/min/kg | 185 | 20 | 10.9 |
| $T_{1/2}$ | h | 0.312 | 0.013 | 4.30 |
| $C_0$ | ng/mL | 431 | 27 | 6.35 |
| $AUC_{last}$ | h*ng/mL | 90 | 10 | 10.9 |
| $AUC_{Inf}$ | h*ng/mL | 91 | 10 | 10.9 |
| $AUC_{_\%Extrap_}$obs | % | 0.656 | 0.104 | 15.8 |
| $MRT_{Inf_}$obs | h | 0.253 | 0.017 | 6.72 |
| $AUC_{last}$/D | h*ng/mL | 90 | 10 | 10.9 |
| $V_{ss_}$obs | L/kg | 2.80 | 0.29 | 10.2 |

**Table S1e. IP administration, compound 6**

**IP Dose 20 mg/kg**

| PK parameters | Unit | Mean | SD | CV(%) |
|---|---|---|---|---|
| $T_{1/2}$ | h | 1.09 | 0.28 | 25.6 |
| $T_{max}$ | h | 0.250 | 0.000 | 0.000 |
| $C_{max}$ | ng/mL | 5580 | 3844 | 68.9 |
| $AUC_{last}$ | h*ng/mL | 3964 | 2189 | 55.2 |
| $AUC_{Inf}$ | h*ng/mL | 3969 | 2195 | 55.3 |
| AUC_%Extrap_obs | % | 0.104 | 0.064 | 60.8 |
| $MRT_{Inf_}$obs | h | 0.660 | 0.042 | 6.42 |
| $AUC_{last}$/D | h*ng/mL | 198 | 109 | 55.2 |

**Table S1f. Oral administration, compound 6**

**PO Dose 20 mg/kg**

| PK parameters | Unit | Mean | SD | CV(%) |
|---|---|---|---|---|
| $T_{1/2}$ | h | 1.71 | 0.14 | 8.00 |
| $T_{max}$ | h | 0.250 | 0.000 | 0.000 |
| $C_{max}$ | ng/mL | 289 | 65 | 22.4 |
| $AUC_{last}$ | h*ng/mL | 207 | 17 | 8.09 |
| $AUC_{Inf}$ | h*ng/mL | 212 | 14 | 6.39 |
| AUC_%Extrap_obs | % | 2.56 | 2.14 | 83.7 |
| $MRT_{Inf_}$obs | h | 1.29 | 0.53 | 40.9 |
| $AUC_{last}$/D | h*ng/mL | 10.3 | 0.8 | 8.09 |
| F | % | 11.7 | 0.7 | 6.39 |

**Table S2. Crystal statistics for the LRH-1 + ME610 + Tif2 complex.**

| **Data collection** | **LRH-1 + ME610 + Tif2** |
|---|---|
| Space group | P 32 2 1 |
| Cell dimensions | |
| a, b, c (Å) | 88.5, 88.5, 106.2 |
| α, β, γ (°) | 90, 90, 120 |
| Resolution (Å) | 40.84-2.70 (2.799-2.702) |
| Rpim | 0.093 (0.708) |
| I / σI | 11.0 (1.4) |
| $CC_{1/2}$ | 99.2 (61.7) |
| Completeness (%) | 98.7 (93.7) |
| Redundancy | 10.0 (9.0) |
| Wilson B-factor | 45.3 |
| **Refinement** | |
| Resolution (Å) | 2.7 |
| No. reflections | 13409 |
| Rwork / Rfree (%) | 17.8 / 21.1 |
| No. atoms | |
| Protein | 2042 |
| Ligand/ion | 39 |
| Water | 31 |
| B-factors | |
| Protein | 72.2 |
| Ligand/ion | 54.6 |
| Water | 57.7 |
| R.m.s. deviations | |
| Bond lengths (Å) | 0.004 |
| Bond angles (°) | 0.63 |
| Ramachandran favored (%) | 99.2 |
| Ramachandran outliers (%) | 0 |
| PDB accession code | 36FH |

**Table S3. Chromatography parameters: Gradient time and the percentage of solvent A and B in mobile phase**

| Time (min) | Solvent A: ACN:water (60:40) | Solvent B: IPA:ACN (90:10) |
|---|---|---|
| 0 | 87 | 13 |
| 0.1 | 87 | 13 |
| 2 | 57 | 43 |

| | | |
|---|---|---|
| 11.5 | 5 | 95 |
| 12 | 5 | 95 |
| 12.5 | 5 | 95 |
| 22.55 | 5 | 95 |
| 22.65 | 22 | 78 |
| 22.75 | 27 | 73 |
| 22.85 | 87 | 13 |
| 22.95 | 87 | 13 |
| 24.35 | 87 | 13 |

**Table S4. The mass spectrometry instrumental parameters for untargeted lipidomic assay**

| Parameter | Setting |
|---|---|
| Ion Source Type | VIP-HESI |
| Capillary Voltage in Positive/Negative Mode (V) | 4500/4500 |
| Scan m/z range | 100-1350 m/z |
| Scan Mode | PASEF |
| Numer of PASEF Runs | 2 |
| Ramp Time (ms) | 100 |
| Accumulation Time (ms) | 100 |
| Ramp Rate (Hz) | 9.43 |
| Nebulizer (bar) | 2 |
| Dry Gas Flow at 230˚C (L/min) | 8 |
| Sheath Gas Flow at 400˚C (L/min) | 4 |
| Collision Energy (eV) | 30 |
| Collision RF (Vpp) | 1100 |
| Funnel 1 RF (Vpp) | 500 |
| Funnel 2 RF (Vpp) | 250 |
| Mass Spectra Peak Absolute Threshold | 100 |
| Mobility Absolute Threshold | 100000 |
| Ion Mobility (1/K0, V*s/cm$^2$) | 0.55-1.9 during 0.32s cycle |
| Stepping | Not applied |

*RF: Radio Frequency

**Table S5. The tolerances and scoring of the lipid annotation**

| | Narrow (lower limit) | Wide (Upper limit) |
|---|---|---|
| m/z (ppm) | 2 | 5 |
| Retention time (min) | 0.05 | 0.1 |
| mSigma[1] | 50 | 1000 |
| MS/MS score[2] | 800 | 400 |
| CCS (%) | 1 | 5 |

[1] A score to describe the difference between the measured isotopic pattern and the theoretical isotopic pattern of the ion.
[2] MS2 patterns

**Table S6. Oxylipin chromatography gradient**

| Time (min) | Solvent A: 10mM Ammonium acetate in water | Solvent B: 10mM Ammonium acetate in Acetonitrile | Flow (mL/min) |
|---|---|---|---|
| 0 | 95 | 5 | 0.4 |
| 0.5 | 95 | 5 | 0.4 |
| 1 | 65 | 35 | 0.4 |
| 2 | 65 | 35 | 0.4 |
| 2.1 | 45 | 55 | 0.4 |
| 5 | 45 | 55 | 0.4 |
| 7 | 25 | 75 | 0.4 |
| 20 | 25 | 75 | 0.4 |
| 20.1 | 95 | 5 | 0.5 |
| 25 | 95 | 5 | 0.5 |

**Table S7. The mass spectrometry instrumental parameters for targeted oxylipin assay**

| | Negative | Positive |
|---|---|---|
| Gas Temperature (˚C) | 290 | 290 |
| Gas Flow (L/min) | 14 | 14 |
| Nebulizer (psi) | 20 | 20 |
| Sheath Gas Temperature (˚C) | 300 | 300 |
| Sheath Gas Flow (L/min) | 11 | 11 |
| Capillary Voltage (V) | 3000 | 3000 |
| Nozzle Voltage (V) | 1500 | 1500 |
| iFunnel High Pressure RF | 150 | 150 |
| iFunnel low Pressure RF | 60 | 60 |

*RF: Radio Frequency

**Table S8. Oxylipins transition list**

| Lipid | Q1 MASS | Q3 MASS |
|---|---|---|
| **11(12)-DiHET** | 337.2 | 319.3/167.1 |
| **11(12)-EET** | 319.2 | 301.3/167.1 |
| **12 R-HETE** | 319.2 | 301.3/179.1 |
| **12(13)-EpOME** | 295.2 | 195.1/183.1 |
| **13 HODE** | 295.2 | 277.2/195.1 |
| **13(S)HOTrE** | 293.2 | 275.2/195.4 |

| | | |
|---|---|---|
| **13(S)-HpODE** | 311.2 | 293.2/113.1 |
| **13-OxoODE (13-KODE)** | 293.2 | 195.1/113.1 |
| **14(15)-EET** | 319.2 | 257.2 |
| **14,15 DiHET** | 337.2 | 207.1/129.1 |
| **20-HETE** | 319.2 | 289.3/275.3 |
| **5 (S)HETE** | 319.2 | 301.3/257.2 |
| **5,6-DiHET** | 337.2 | 319.3/145 |
| **8(9)-DiHET** | 337.2 | 185.1/127.1 |
| **8(9)-EET** | 319.2 | 301.3/257.2 |
| **8-isoprostane** | 353.2 | 309.2/193.1 |
| **9(S)-HODE** | 295.2 | 277.2/141.1 |
| **9(S)HOTrE** | 293.2 | 275.2/171.2 |
| **9(S)-HpODE** | 311.2 | 293.2/185.1 |
| **9,10 DiHOME** | 313.2 | 277.3/201.1 |
| **9-HETE** | 319.2 | 167.1/151.1 |
| **leukotriene B4 (LTB4)** | 335.2 | 317.2/195.1 |
| **Maresin 2** | 359.2 | 232.1/221.1 |
| **Maresin1** | 359.2 | 250.1/93.1 |
| **PGE2** | 351.2 | 315.2/271.2 |
| **PGF2a** | 353.2 | 309.2/193.1 |
| **Resolvin D1** | 375.2 | 215.2/141 |
| **Resolvin D2** | 375.2 | 277.2/175.1 |
| **Resolvin D3** | 375.2 | 357.2/147.1 |
| **Resolvin E1** | 349.2 | 205.1/195.1 |
| **TXB2** | 369.2 | 169.1 |
| **2-Arachidonoyl Glycerol (2AG)** | 379.3 | 91/67.1 |
| **alpha-linolenoyl ethanolamide** | 322.3 | 81.1/62.2 |
| **Arachidonoyl-Ethanolamide (AEA)** | 348.3 | 91.1/62.2 |
| **dihomo-gamma-linolenoyl ethanolamide** | 350.3 | 81.1/62.2 |
| **Docosahexaenoyl Ethanolamide (DHEA)** | 372.3 | 91.1/62.2 |
| **docosanoyl ethanolamide** | 384.4 | 62.2/57.2 |
| **Docosatetraenoyl Ethanolamide (DEA)** | 376.3 | 79.1/62.2 |
| **Linoleoyl Ethanolamide (LEA)** | 324.3 | 62.2/55.2 |
| **oleamide** | 282.3 | 83.1/69.1 |
| **Oleoyl Ethanolamide (OEA)** | 326.3 | 62.2/55.2 |
| **oxy-Arachidonoyl Ethanolamide (oxy-AEA)** | 364.3 | 105.1/91.1 |
| **Palmitoyl Ethanolamide** | 298.3 | 69.2/62.2 |
| **PGE2 Ethanolamide (PGE2-EA)** | 396.3 | 105/79 |
| **Prostagladin** | 444.3 | 409.2/391.2/299.2 |
| **Stearoyl Ethanolamide (ceramid)** | 328.3 | 62.2/57.2 |

### Supplementary Figures

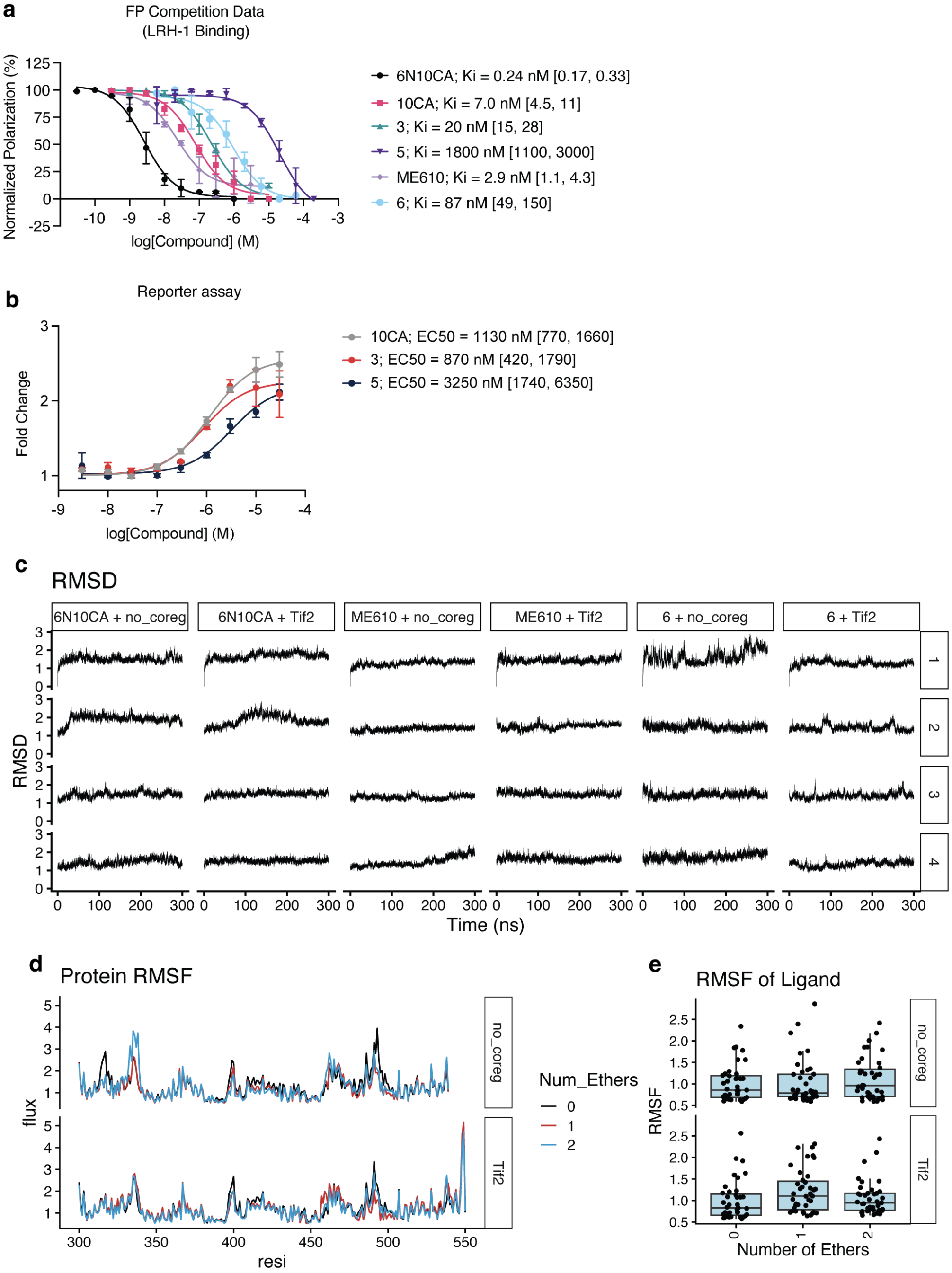

**Figure S1. Characterization of ether-containing compounds. a.** Normalized dose-response curves from the FP competition LRH-1 binding assay. Each point represents the mean +/- SD from two independent experiments. **b.** Dose-response curves from the LRH-1 luciferase reporter assay. Each point is the mean +/- SD from three experiments, each conducted in triplicate. *Insets* show the $EC_{50}$ values. **c-e.** Exploratory or QC plots from MD simulations with six LRH-1 complexes, showing root mean square deviation (RMSD) in *c,* and root mean square fluctuation (RMSF) of the protein (*d)* and ligand (*e*) components of the complexes. In *e,* each point is a ligand atom (hydrogens excluded); boxes represent the medians +/- the interquartile range (IQR); whiskers are 1.5 the IQR.

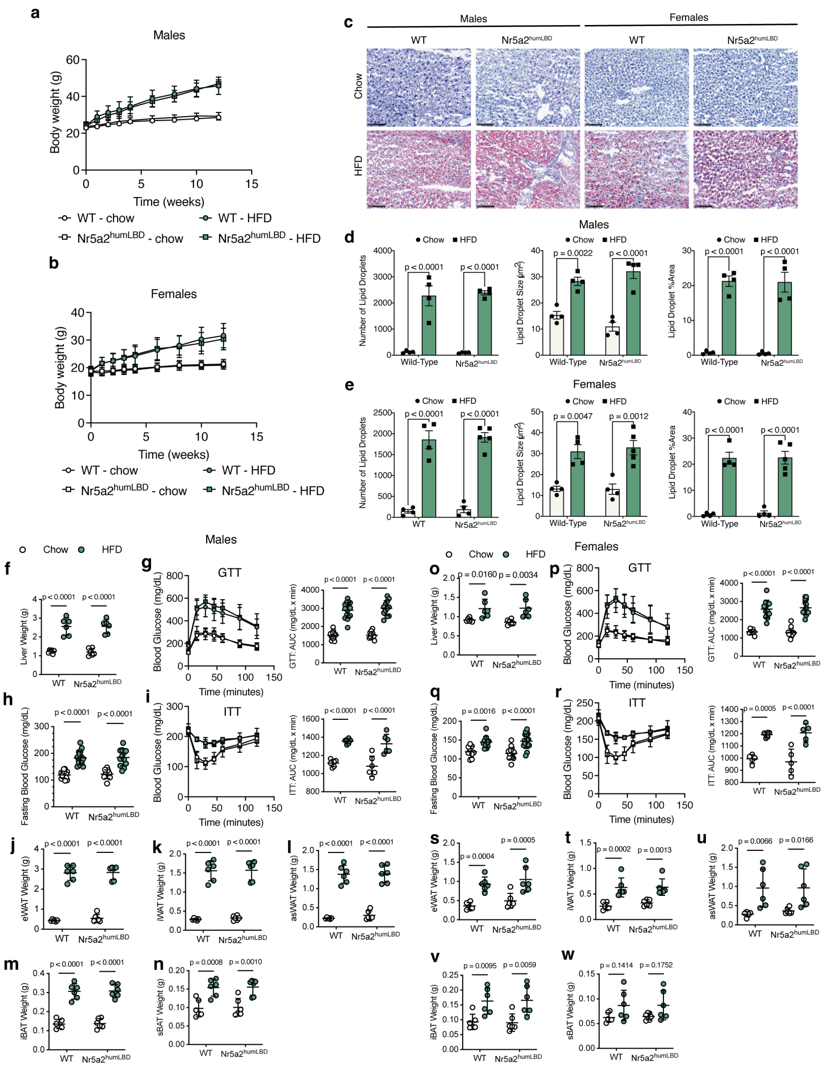

**Figure S2. *Nr5a2$^{humLBD}$* and WT mice respond similarly to HFD.** 10-week-old wild type (WT) C57Bl/6 mice or *Nr5a2$^{humLBD}$* mice were fed normal chow or HFD for 12 weeks. **a.** Body weights of male mice over time. Each point represents the mean +/- SEM for n = 16 (WT+chow); n = 11 (Nr5a2$^{humLBD}$+chow); n = 16 (WT+HFD); n = 12 (Nr5a2$^{humLBD}$+HFD). **b.** Body weights of female mice over time. Each point represents the mean +/- SEM for n = 9 (WT+chow); n = 16 (Nr5a2$^{humLBD}$+chow); n = 12 (WT+HFD); n = 17 (Nr5a2$^{humLBD}$+HFD). **c.** Representative images from liver tissues of mice fed chow or HFD and stained with oil red o (ORO) to detect lipid droplets. **d-e**. Quantification of lipid droplets from male (*d*) or female (*e*) mice. Each bar represents mean +/- SD (n = 4-5). **f,o.** Liver weights in male (*f*) and female (*o*) mice increase on HFD to a similar degree for both genotypes. Each point represents a different mouse; middle lines are the means, error bars are SD. **g,p**. *Left,* time-response curves and *right* AUC for mice in the oral glucose tolerance test (GTT) in males (*g*) and females (*p*). **h,q.** Fasting blood glucose (FBG) in HFD-fed versus chow-fed mice (males in *h*; females in *q*). Each point represents a different mouse; middle lines are the means, error bars are SD. **i,r.** Results from the insulin tolerance test (ITT), plotted as in *g* and *p*. **j-w**. Measurements of adipose depots in male (*j-n*) and female mice (*s-w*). Each point represents a different mouse; middle lines are the means, error bars are SD. Graph *insets* in panels *d-w* show p-values, calculated by two-way ANOVA; Sidak's multiple comparisons test.

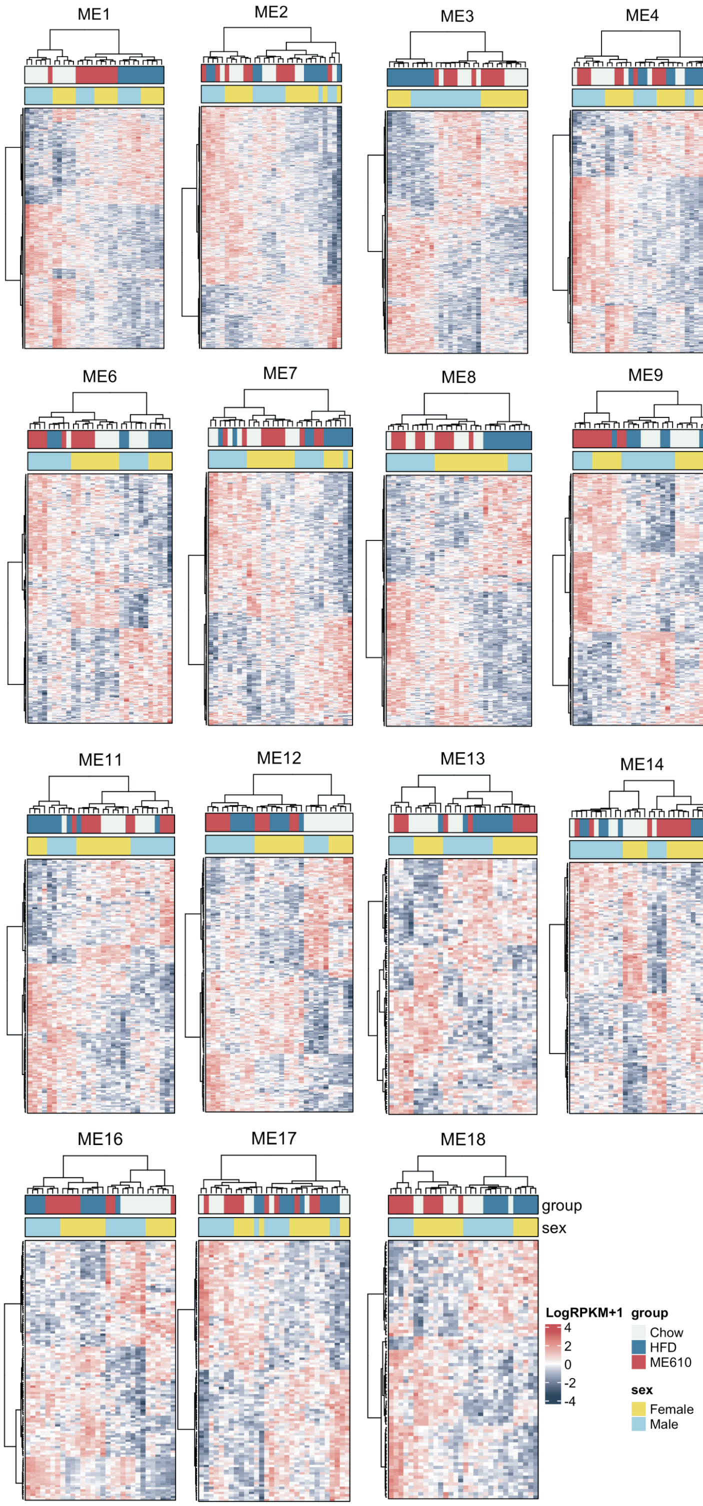

**Figure S3. Heatmaps from RNA-seq WGCNA modules.** $Log_2$ RPKM + 1 values for genes in each module were clustered by Euclidean distance and Ward's agglomeration. The clustering patterns were used to inspect whether mice in each module were separated mainly by diet (*e.g.,* ME5, ME12, where control group is separated from the two other groups) or whether the dominant trend showed ME610 as different from the HFD group (*e.g.,* ME3, ME8, ME9).

6. LaFrate AL, Carlson KE, Katzenellenbogen JA. Steroidal bivalent ligands for the estrogen receptor: design, synthesis, characterization and binding affinities. Bioorg Med Chem.

2009;17(10):3528-35. Epub 20090412. doi: 10.1016/j.bmc.2009.04.016. PubMed PMID: 19394231; PMCID: PMC3178464.
